# Distinct sequencing success at non-B-DNA motifs

**DOI:** 10.1101/2022.06.13.495922

**Authors:** Matthias H. Weissensteiner, Marzia A. Cremona, Wilfried Guiblet, Nicholas Stoler, Robert S. Harris, Monika Cechova, Kristin A. Eckert, Francesca Chiaromonte, Yi-Fei Huang, Kateryna D. Makova

## Abstract

Modern sequencing technologies are not error-free, and might have elevated error rates at some locations of the genome. A potential cause for such elevated error rates is the formation of alternative DNA structures (non-B DNA), such as G-quadruplexes (G4s), Z-DNA, or cruciform structures, during sequencing. Approximately 13% of the human genome has the potential to form such structures, which have been previously shown to affect the activity of DNA polymerases and helicases. Here we tested whether motifs with the potential to form non-B DNA (non-B motifs) influence the sequencing success of three major sequencing technologies—Illumina, Pacific Biosciences (PacBio) HiFi, and Oxford Nanopore Technologies (ONT). We estimated sequencing success by computing the rates of single-nucleotide, insertion, and deletion errors, as well as by evaluating mean read depth and mean base quality. Overall, all technologies exhibited altered sequencing success for most non-B motif types. Single-nucleotide error rates were generally increased for G-quadruplexes (G4s) and Z-DNA motifs in all three technologies. Illumina and PacBio HiFi deletion error rates were also increased for all non-B types except for Z-DNA motifs, while in ONT they were increased substantially only for G4 motifs. Insertion error rates for non-B motifs were highly elevated in Illumina, moderately elevated in PacBio HiFi, and only slightly elevated in ONT. Using Poisson regression modeling, we evaluated how non-B DNA motifs and other factors influence sequencing error profiles. Using the error rates at non-B motifs, we developed a probabilistic approach to determine the number of false-positive single-nucleotide variants (SNVs) in different sample size and variant frequency cutoff scenarios, as well as in previously generated sequencing data sets (1000Genomes, Simons Genome Diversity Project, and gnomAD). Overall, the effect of non-B DNA on sequencing should be considered in downstream analyses, particularly in studies with limited read depth—e.g., single-cell and ancient DNA sequencing, as well as sequencing of pooled population samples—and when scoring variants with low frequency (e.g., singletons). Because each sequencing technology analyzed has a unique error profile at non-B motifs, a combination of different technologies should be considered in future sequencing studies of such motifs, to maximize accuracy.

## Introduction

DNA conformations that deviate from the canonical right-handed double-helix with ten nucleotides per turn are collectively termed ‘non-B DNA’ (Zhao et al. 2010). The ability of the DNA molecule to fold into such alternative structures depends on the presence of certain sequence motifs (thereby called ‘non-B motifs’), which range in size from tens to hundreds of base pairs and account for a substantial portion of an organism’s genome (e.g., ~13% of the human genome)(Guiblet et al. 2018). Non-B DNA motifs can form distinct non-B DNA structures depending on their sequence (Fig. 1). A-phased repeat motifs, which consist of tracts of three to nine adenines or thymines (A-tract) separated by at least 4 bp (spacer), can facilitate bent double helix structures (Barbic, Zimmer, and Crothers 2003; Koo, Wu, and Crothers 1986). In G-quadruplex (G4) motifs, which consist of at least four blocks of at least three guanines separated by one to seven arbitrary bases, the guanines from different blocks can bind to each other via Hoogsteen hydrogen bonds forming stems, with the arbitrary bases forming loops (Sen and Gilbert 1988; Burge et al. 2006). Direct repeat motifs, which consist of two copies of the repeated unit separated by a non-repetitive spacer, can misalign leading to slipped-strand structures, with looped out bases (Sinden, Pytlos-Sinden, and Potaman 2007). Inverted repeat motifs, which consist of repetitive sequences complementary to each other (e.g., 5’-GACTGC and GCAGTC-3’) separated by a non-repetitive spacer, are capable of forming hairpins and cruciform structures (Nag and Petes 1991). Mirror repeat motifs that consist of stretches of homopurines:homopyrimidines arranged in a mirrored fashion, separated by a spacer, can form triple-helix (H-DNA) structures (Htun and Dahlberg 1988). Finally, Z-DNA motifs, which consist of alternating pyrimidines and purines, such as (CG:CG)_n_ or (CA:TG)_n_, can form left-handed zig-zag DNA structures (A. H.-J. Wang et al. 1979; Singleton et al. 1982).

**Figure 1.**
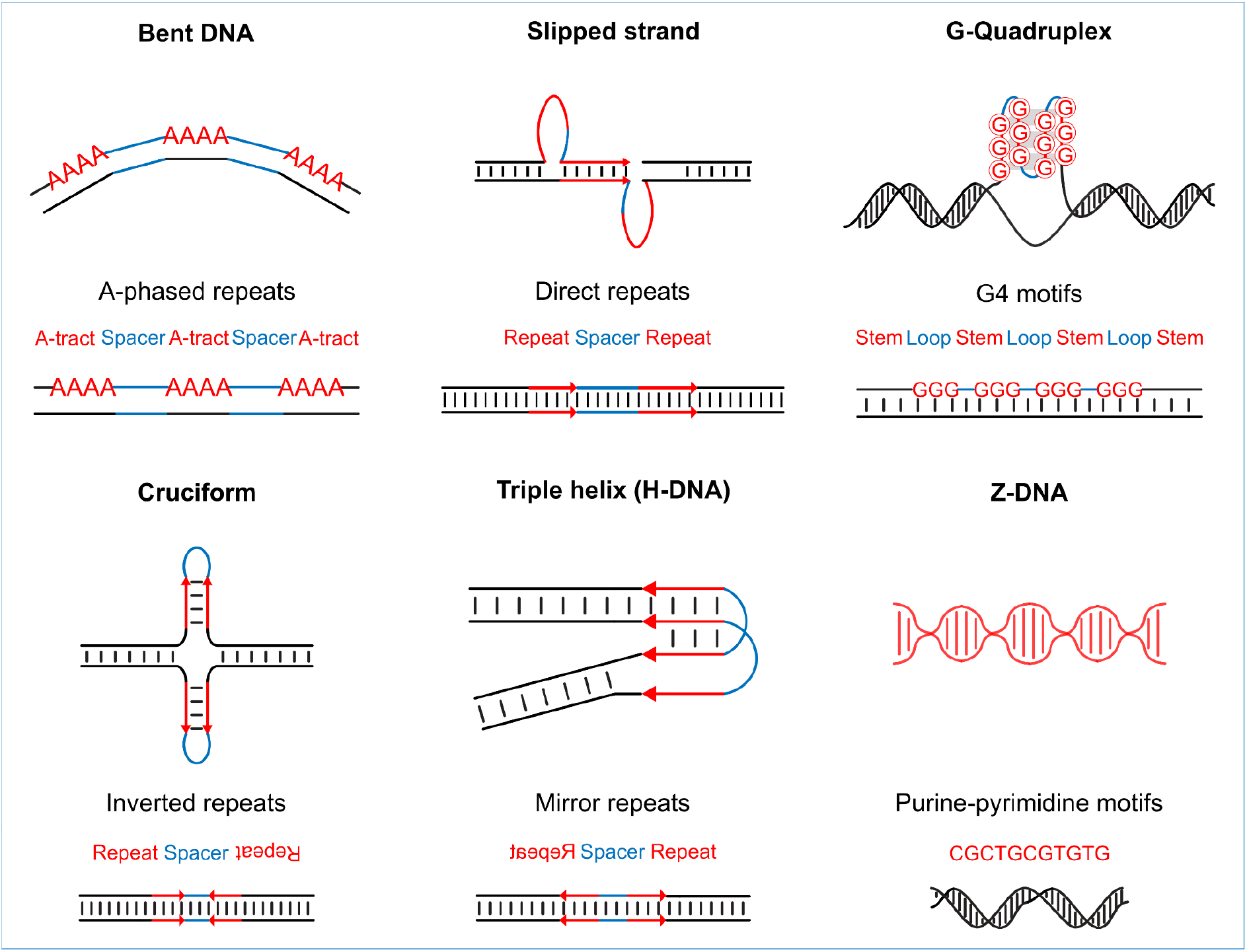
Types of non-B DNA structures. Shown are six types of alternative DNA structures (non-B DNA) with their respective underlying motifs and sequence arrangements. Mirror repeat motifs that form triple helix (H-DNA) structures consist of stretches of homopurines:homopyrimidines arranged in a mirrored fashion and separated by a spacer.

Non-B DNA structures can form *in vivo (Biffi et al. 2013; Hänsel-Hertsch et al. 2016; Shin et al. 2016)* and play an important role in essential cellular processes such as gene expression and DNA replication (Ghosh and Bansal 2003; A. Jain, Wang, and Vasquez 2008; G. Wang and Vasquez 2014). Non-B DNA may influence DNA synthesis because some non-B DNA structures have been shown to obstruct the progression and affect accuracy of DNA polymerases (Mirkin and Mirkin 2007; G. Wang and Vasquez 2014). For instance, hairpins, G4 structures, and triple helices have been linked to the inhibition of polymerase activity *in vitro* (Mirkin and Mirkin 2007). Furthermore, certain non-B-forming motifs were found to influence polymerase kinetics and affect sequencing errors of PacBio sequencing and to affect sequencing errors (Guiblet et al. 2018).

Two major high-throughput sequencing technologies—Illumina and PacBio—are based on synthesis with DNA polymerases. In Illumina sequencing, the *Pyrococcus*-derived Phusion polymerase (Quail et al. 2011) is involved in the bridge amplification of the template strand producing clusters, which is followed by sequencing by synthesis via the incorporation of fluorescently active nucleotides (Metzker 2010). In PacBio sequencing, an engineered bacteriophage phi29 DNA polymerase (Eid et al. 2009) incorporates fluorescently labeled nucleotides, whose sequence is then determined by a laser (Logsdon, Vollger, and Eichler 2020). In contrast, in Oxford Nanopore Technologies (ONT) sequencing, DNA polymerases are absent, and the nucleotide sequence is determined by changes in electric current caused by the passage of a single-stranded DNA through a protein nanopore, located in a synthetic membrane (Logsdon, Vollger, and Eichler 2020; M. Jain et al. 2016). While it is currently unknown what effect non-B DNA structures might have on the activity of the engineered T4 phage Dda helicase, the motor protein used to unwind and propel the double-stranded DNA molecule during ONT sequencing (Logsdon, Vollger, and Eichler 2020; Daniel and Deamer 2019), there are examples of helicases being involved in the resolution of non-B structures (e.g., A. Jain et al. 2010). Although the conceptual approach of nucleotide sequence determination is considerably different among these three sequencing technologies, it is conceivable that the enzymes they recruit—polymerases and helicases—contribute to the technology-specific sequence error profiles. Since non-B DNA structures have been shown to affect DNA-processing enzyme activity (Mirkin and Mirkin 2007), their effect on sequencing error profiles should be considered.

Errors have been a major concern ever since the invention of DNA sequencing, because they may drastically influence the downstream analysis and interpretation of sequencing data. Randomly occurring sequencing errors can be alleviated by increasing the read depth per site, enabling a consensus approach to identify the true nucleotide at a given locus (Nielsen et al. 2011). In cases where the amount and/or quality of input DNA (e.g., ancient DNA studies (Slatkin and Racimo 2016)) or budget constraints do not allow high read depth, sequencing errors may have a major effect on the downstream analyses (Shafer et al. 2017). Sequencing errors that occur non-randomly are expected to have an even greater impact, because such errors are expected to occur even with high read depth, resulting in them more likely to be identified as false positive genetic variants. Examples hereby are the coverage bias against GC-rich sequences in Illumina sequencing (Shafer et al. 2017; Aird et al. 2011), or the tendency of (earlier) versions of ONT sequencing to erroneously collapse homopolymer runs. Thus far, approaches to mitigate these issues have mostly included stringent computational filtering or the use of multiple independent sequencing technologies, which may drastically reduce the amount of usable data or may be cost-prohibitive, respectively.

In this study, we investigated a potential association between non-B DNA motifs and sequencing success for three major sequencing technologies (Illumina, HiFi mode of PacBio, and ONT). We compiled annotations of non-B-forming sequences in the human genome and used sequencing data from the Genome in a Bottle consortium (GIAB, Zook et al. 2016) to detect errors. We found sequencing success to be significantly altered for almost all types of non-B DNA motifs (as compared to canonical B DNA) in all three sequencing technologies studied. Thus, the presence of non-B DNA should be considered in downstream analyses, especially in low-depth studies.

## Results

We contrasted sequencing success, as measured by error rate, sequencing depth, and base quality, between motifs with non-B-DNA-forming potential (‘non-B motifs’) and control B DNA sequences. This was performed for sequencing reads generated with Illumina, PacBio, and ONT sequencing technologies. We utilized a dataset in which these three technologies were applied to the same sample, an Ashkenazim son (HG002) from the Genome in a Bottle Consortium (GIAB, Zook et al. 2016). For PacBio, the effects of non-B motifs on continuous long-read sequencing were evaluated previously (Guiblet et al. 2018), and thus we focused on the HiFi circular consensus reads, which achieve low error rates after multiple passes over the same template. We acquired the genomic coordinates of A-phased repeats, direct repeats, inverted repeats, mirror repeats, and Z-DNA motifs from the non-B DNA database (Cer et al. 2012). The motifs potentially forming G4 structures were annotated with Quadron (Sahakyan et al. 2017).

Prior to assessing sequencing success, we applied two filtering schemes that differed in stringency. With moderate filtering, we included the repetitive portions of the genome that are enriched in non-B motifs, while acknowledging that they might be affected by misalignment and ambiguous mapping of sequencing reads. With stringent filtering, we obtained ‘cleaner’ (largely void of artifacts), but smaller sets of non-B motifs by filtering out repeats and microsatellites, as well as overlapping motifs of different types. In the moderately filtered set, we restricted the size of motifs to the range from 10 to 1,000 bp (mean=26 bp, median=16 bp, Table S1), excluded motifs that overlapped with other non-B motifs of the same type, and excluded motifs with an average mappability below 1 (see Methods). As a result, we retained 5,360,356 non-B motifs, covering ~137 Mb of the genome (Table S1). In the stringently filtered sets, we additionally removed non-B DNA motifs and controls that had ≥1-bp overlap with a repetitive element or a microsatellite, were within 50 bp from another non-B motif of any type, or had an average base quality of a Phred score below 30 for Illumina or below 73.2 for HiFi (these are corresponding thresholds for the two technologies; see Methods). As a result, we retained 710,553 motifs (13.5 Mb), 570,217 motifs (10.9 Mb), and 721,479 motifs (13.9 Mb) in the Illumina, HiFi, and ONT data sets, respectively (Table S1). For each filtering scheme, each sequencing technology, and each type of non-B motifs, we also generated a set of random control sequences, matching the corresponding motif set in number and length, and excluding all non-B motifs, sequencing gaps, and ≥7-bp homopolymer runs. Prior to scoring sequencing errors, we removed biological variants as annotated in the GIAB true variant set (Zook et al. 2016).

### Single-nucleotide mismatch error rates

#### Illumina

We detected significantly higher per-motif single-nucleotide mismatch (SNM) error rates for direct repeats, G4 motifs, and Z-DNA motifs as compared to the respective controls for Illumina (average per-motif and per-control error rates shown as black circles are provided in Fig. 2A, see also Table S2A; Benjamini-Hochberg-corrected t-test p-values are provided in Table S3). This was the case for both moderately (1.35-, 2.39-, and 1.64-fold, respectively) and stringently (1.13-, 2.00-, and 1.49-fold, respectively) filtered sets (Fig. 2B). Error rates for A-phased and inverted repeats were significantly lower as compared to the respective controls (0.90- and 0.93-fold for moderate filtering and 0.90- and 0.92-fold for stringent filtering, respectively), whereas mirror repeats had error rates significantly higher (1.20-fold) than controls for the moderately filtered set and not significantly different from controls for the stringently filtered set. Here and below we also computed the aggregate error rates (sum of errors divided by sum of aligned nucleotides) in motifs and corresponding controls (orange triangles in Fig. 2A; Table S2A; Fig. S1), which in most cases were similar to the average per-motif and per-control error rates.

**Figure 2.**
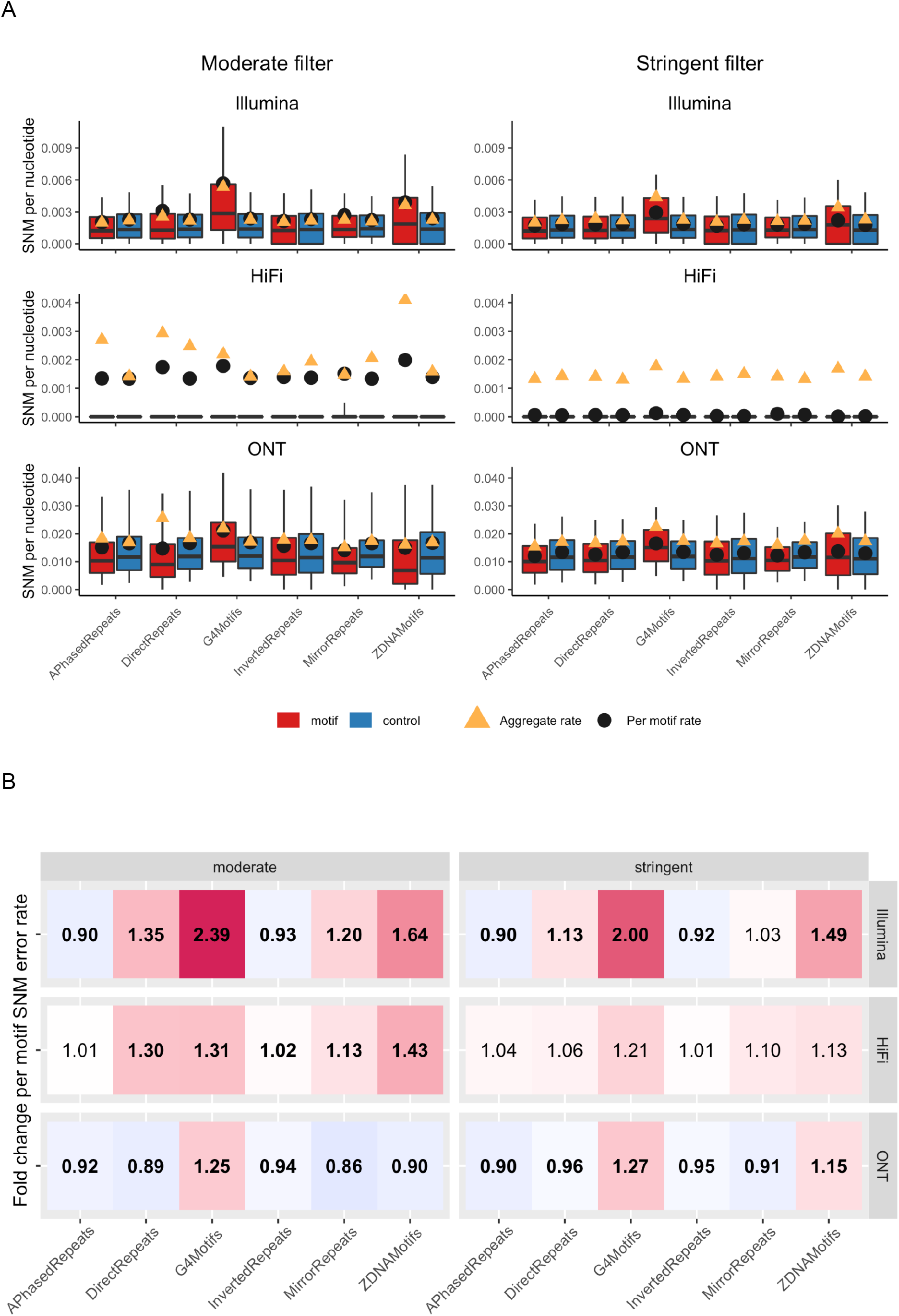
Single-nucleotide mismatch (SNM) error rates in non-B motifs. **(A)** Boxplots of per-motif SNM error rates. Values above the 90th percentile are excluded from the boxplots in order to better visualize the bulk of the distributions. The left panel shows the moderately filtered motif set, the right panel the stringently filtered set, and the three rows correspond to the different technologies (Illumina, HiFi, and ONT). Red and blue boxes correspond to motifs and controls, respectively, black dots mark per-motif means, and orange triangles aggregate error rates (sum of all errors divided by sum of all aligned nucleotides). Note that y-axes differ among technologies. **(B)** Heat maps visualizing fold changes in per-motif means of SNM error rates between motifs and corresponding controls. Red (green) shades indicate higher (lower) error rates in non-B motifs than in controls, with fold change values reported in each cell of the map. When these values are in bold, per-motif means were significantly different between motifs and controls (t-test p-values corrected for multiple testing smaller or equal to 0.05). Also here, left and right panels correspond to moderately and stringently filtered sets, respectively, and rows correspond to Illumina, HiFi, and ONT technologies, respectively.

To disentangle the factors contributing to the differences in average per-motif and per-control SNM error rates, we fit a Poisson regression model for each non-B motif type separately. In each model, the number of SNMs (in a motif or a control sequence) is the response, and the predictors are an indicator of ‘non-B motif vs. control’, the nucleotide composition, the motif length, the occurrence of 3-7-bp homopolymer runs, and the total number of sequenced nucleotides across all reads mapping to a motif (see Methods for details). The results were similar between the moderately and stringently filtered sets (Table S4); below we discuss results for the former. Deviance explained, a measure of the explanatory power provided by the model, ranged from 1.7% (for A-phased repeats) to 16.7% (for G4 motifs; Table S4). To estimate the contribution of each predictor, we followed the approach described in Kelkar et al. (2011) and fitted reduced models where we left out individual predictors one at a time, and calculated the reduction in deviance explained compared to the full model. Note however that, due to potential interactions between predictors, effects might not be additive. The removal of the ‘non-B motif vs. control’ predictor reduced the deviance explained the most for models for Z-DNA motifs (by 31.5%), direct repeats (by 11.9%), and G4 motifs (by 5.3%). Removing nucleotide composition from the model substantially reduced deviance explained in models for mirror repeats (by 97.8%) and A-phased repeats (by 89.4%). Here and below such percentages among models should be compared by taking into account the overall deviance explained by each model (Table S4).

To add an orthogonal approach to evaluate sequencing errors, we calculated SNM error rates as mismatches in the overlaps between Illumina read pairs in non-B motifs of the moderately filtered set, using sequencing data of the same individual as above (HG002)(Stoler and Nekrutenko 2021). Given that this approach identifies far fewer errors, because it includes only the overlapping part of the reads (see Methods for details), the resulting data set was admittedly smaller than the original one: in total, we analyzed between 494,961 (Z-DNA motifs) and 1,395,058 (inverted repeats) of overlapping nucleotides in non-B motifs (Table S5). Overall, this analysis showed similar trends (Table S5) to the analysis presented above. In particular, G4 and Z-DNA motifs exhibited substantially higher SNM error rates than controls, with 3.60-fold and 1.90-fold increases, respectively (Fig. S5), while SNM error rates for the other non-B motif types were more similar to controls, and even significantly decreased for direct and inverted repeats (Chi-square test, Table S5).

#### PacBio HiFi

Even though SNM error rates for HiFi data were overall low (Table S2B), they were significantly elevated for several non-B motif types compared to controls in the moderately filtered set (Fig. 2). Similar to Illumina, for HiFi we found significantly elevated per-motif SNM error rates compared to controls for direct repeats, G4 motifs, inverted repeats, mirror repeats, and Z-DNA motifs (1.30-, 1.31-, 1.02-, 1.13-, and 1.43-fold increases). The percentage of deviance explained for the Poisson regression models was usually low (Table S4), and thus the contribution of each individual predictor to the variation in SNM error rates could not be reliably determined. For the stringently filtered set, we found no significant differences between per-motif vs. per-control SNM error rates (Fig. 2).

#### ONT

The overall ONT SNM error rate was an order of magnitude higher than that for Illumina or HiFi (Table S2B). All comparisons of ONT SNM per-motif vs. per-control rates, for both moderately and stringently filtered sets, were statistically significant (Fig. 2). Similar to Illumina and HiFi data, ONT reads exhibited sizably higher per-motif SNM error rates in G4 motifs than in controls (1.25-fold and 1.27-fold for moderately and stringently filtered sets, respectively). However, fold differences of SNM error rates for the other non-B motifs vs. controls were relatively small in magnitude (from 0.89-fold to 1.15-fold) and for Z-DNA were inconsistent between the moderately and stringently filtered data sets. The explanatory power of the Poisson regression models for the ONT data was low (Table S4), thus the contribution of individual predictors could not be reliably determined.

#### SNM error rates for different parts of non-B motifs

We found a conspicuous pattern of variation in SNM error rates between different parts of non-B motifs, such as repeat arms and spacers in the repeat motifs, and stems and loops in the G4 motifs (Fig. 3; Table S6). These patterns were qualitatively consistent between filtering schemes for all non-B DNA types and technologies (Fig. 3). Below we present fold-differences for the moderately filtered set. Per-spacer error rates were significantly higher (Table S6) than per-repeat-tract rates for A-phased repeats across all three technologies analyzed, with 1.40-fold for Illumina, 1.05-fold for HiFi, and 1.45-fold for ONT. Likewise, error rates were significantly elevated in spacers compared to repeat arms for direct repeats (1.39-fold for Illumina, 1.19-fold for HiFi, and 1.18-fold for ONT). For inverted repeats, the rates were also significantly elevated in spacers compared to repeat arms in all three technologies, but the fold-increases were small (≤1.10). For mirror repeats, error rates were significantly elevated in spacers compared to repeat arms for Illumina and ONT (1.23-fold and 1.16-fold, respectively), but were the same between these two parts of repeats for HiFi. For G4 motifs, we observed contrasting patterns among technologies; while error rates were significantly elevated in loops compared to stems in Illumina and HiFi (5.76- and 1.20-fold, respectively), they were decreased in loops vs. stems (0.83-fold) in ONT. The pattern observed in ONT likely reflects the known elevated error rates at homopolymers (Bowden et al. 2019), which are present in G4 stems.

**Figure 3.**
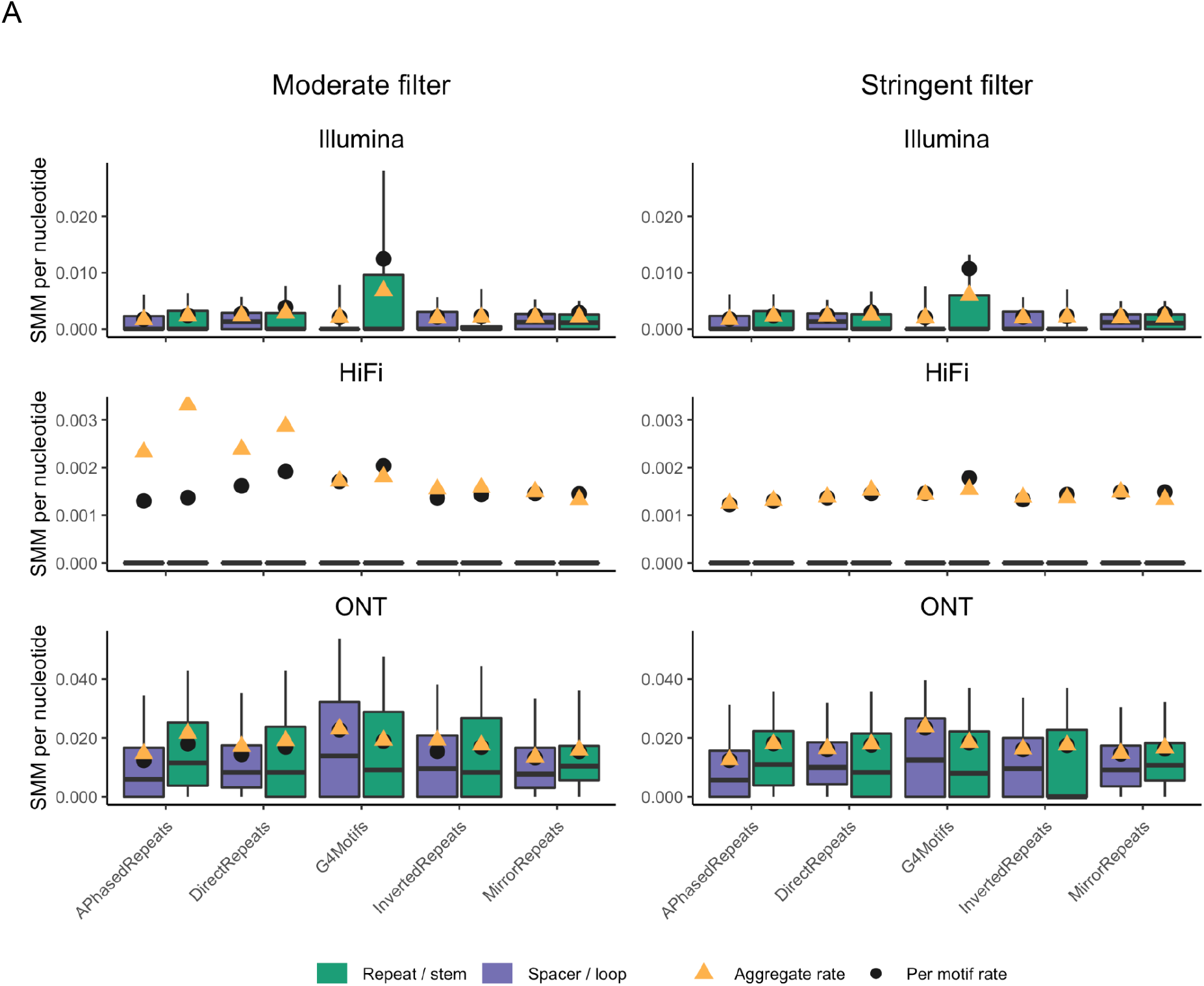
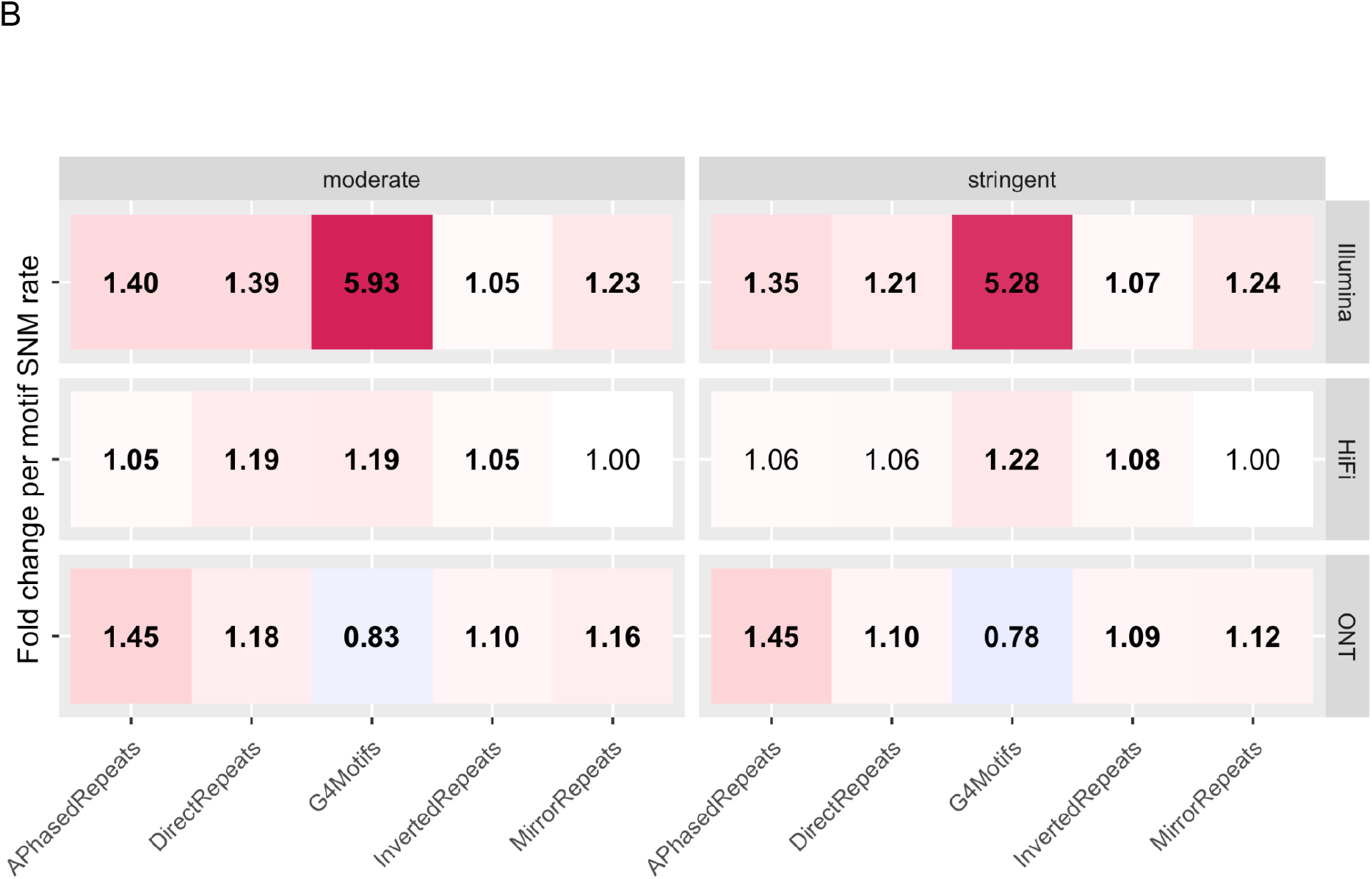
Single-nucleotide error rates in non-B motif sub-regions. **(A)** Boxplots of per-motif SNM error rates of sub-regions of non-B motifs. A-phased, direct, inverted, and mirror repeats are divided into repeat arms and spacer, G4 motifs are divided into stem (G-tract) and loop. Values above the 90th percentile are excluded from the boxplot in order to better visualize the bulk of the distributions. The left panel shows the moderately filtered set, the right panel the stringently filtered set, and the three rows correspond to the different technologies (Illumina, HiFi, and ONT). Purple and green boxes correspond to repeat/stem and spacer/loop subregions, respectively, black dots mark values for per-motif means, orange triangles aggregate error rates (sum of all errors divided by sum of all aligned nucleotides). Note that y-axes differ among technologies. **(B)** Heat maps visualizing fold-changes in per-motif means of SNM error rates between different sub-regions of non-B motifs. Red (green) shades indicate higher (lower) error rates in loops and spacers than in stems and repeat arms, with fold change values reported in each cell of the map. When these values are in bold, per-motif means were significantly different between motifs and controls (t-test p-values corrected for multiple testing smaller or equal to 0.05). Also here, left and right panels correspond to moderately and stringently filtered sets, respectively, and rows correspond to Illumina, HiFi, and ONT technologies, respectively.

### Deletion errors

#### Illumina

To compare deletion error rates between non-B motifs and controls, we divided the number of deletion errors by the number of aligned nucleotides (across all reads) of the motif or control sequence. The overall deletion error rates were low for the Illumina dataset (Table S2B). Deletion error rates were significantly higher for motifs than for controls in both stringently and moderately filtered sets for direct repeats (8.23- and 4.31-fold for the moderately and strongently filtered datasets, respectively), G4 motifs (3.00- and 3.02-fold), and inverted repeats (1.60- and 1.25-fold, Fig. 4, Table S3). Additionally, deletion error rates were significantly higher in motifs than in controls in the moderately filtered set for Z-DNA motifs (12.7-fold), mirror repeats (4.84-fold), and A-phased repeats (1.20-fold; Fig. 4, Table S2A). The percentage of deviance explained in the Poisson regression models for per-motif deletion error rates ranged from 0.4% (for A-phased repeats) to 16.3% (for Z-DNA motifs; Table S4). The removal of the ‘non-B motif vs. control’ predictor led to a reduction of deviance explained across all non-B motif types, with a 1.98-83.2% reduction depending on the non-B motif type, with particularly high contribution of this predictor for models concerning Z-DNA motifs (83.2%) and direct repeats (64.5%; Table S4).

**Figure 4.**
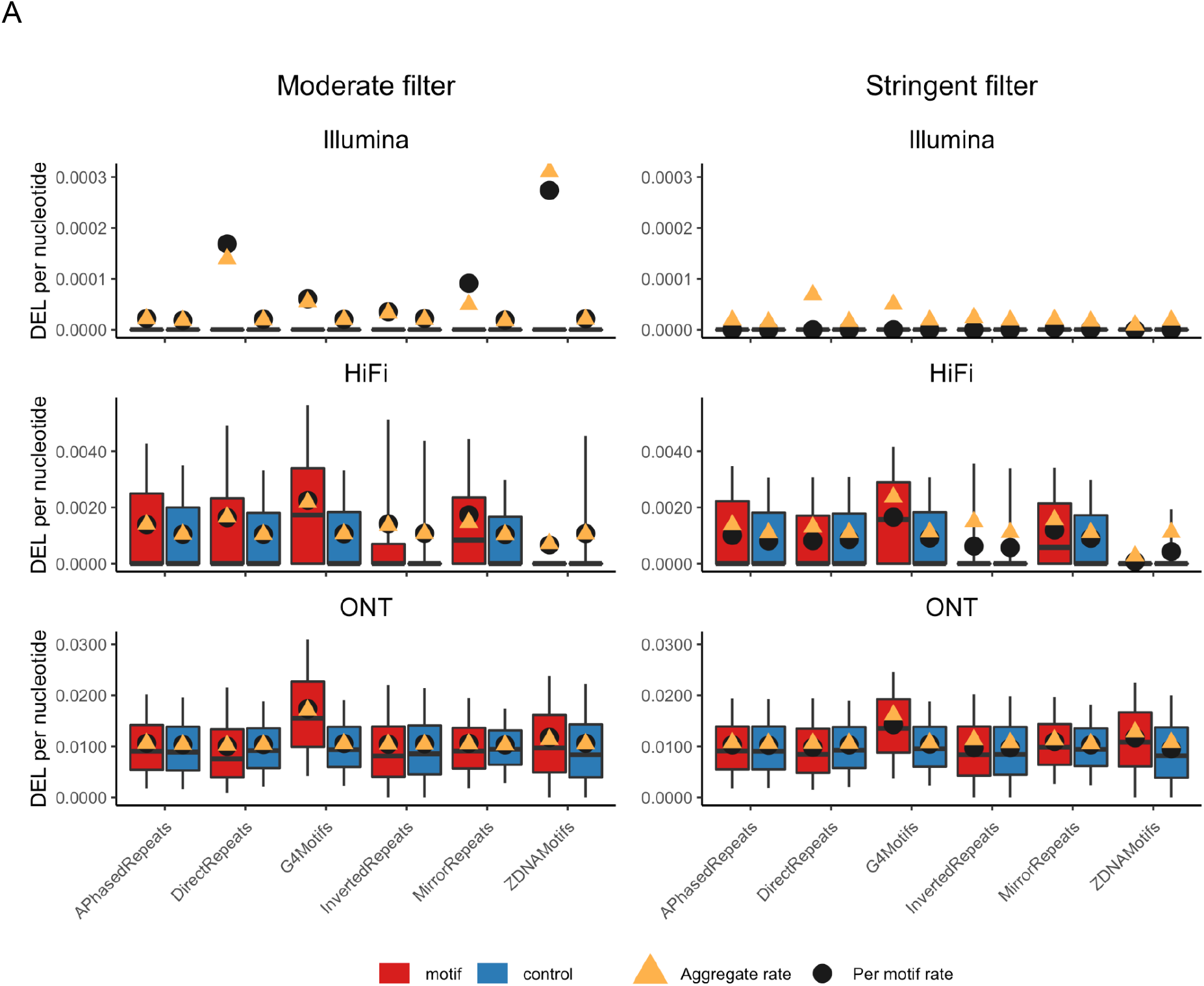
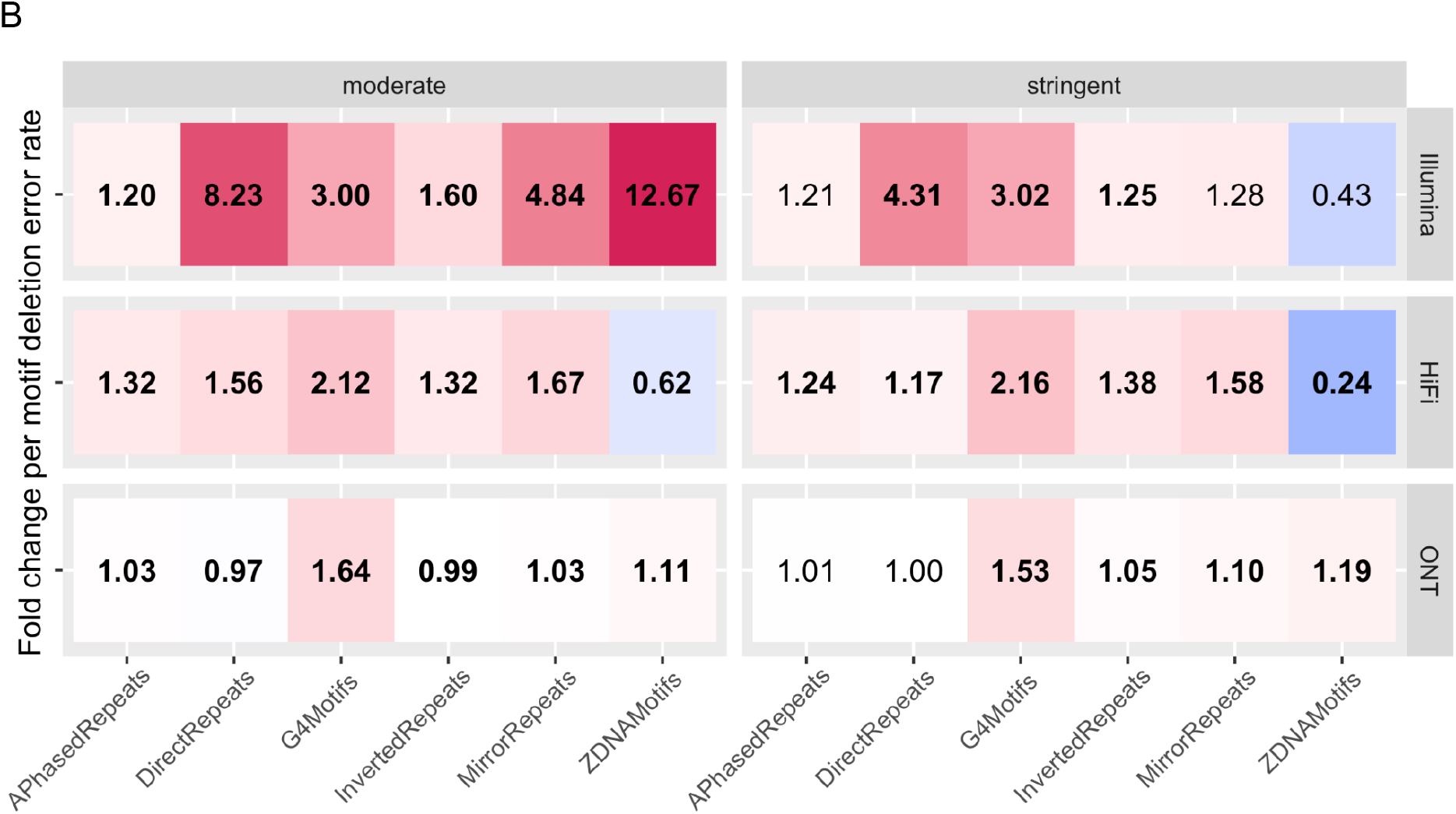
Deletion error rates in non-B motifs. **(A)** Boxplots of per-motif deletion error rates. Values above the 90th percentile are excluded from the boxplots in order to better visualize the bulk of the distributions. The left panel shows the moderately filtered motif set, the right panel the stringently filtered set, and the three rows correspond to the different technologies (Illumina, HiFi, and ONT). Red and blue boxes correspond to motifs and controls, respectively, black dots mark per-motif means, and orange triangles aggregate error rates (sum of all deletion errors divided by sum of all aligned nucleotides). Note that y-axes differ among technologies. **(B)** Heat maps visualizing fold changes in per-motif means of deletion error rates between motifs and corresponding controls. Red (green) shades indicate higher (lower) error rates in non-B motifs than in controls, with fold change values reported in each cell of the map. When these values are in bold, per-motif means were significantly different between motifs and controls (t-test p-values corrected for multiple testing smaller or equal to 0.05). Also here, left and right panels correspond to moderately and stringently filtered sets, respectively, and rows correspond to Illumina, HiFi, and ONT technologies, respectively.

#### PacBio HiFi

The overall deletion error rate for HiFi was approximately an order of magnitude higher than that for Illumina (Table S2B). All non-B motif types except for Z-DNA motifs exhibited significantly elevated per-motif deletion error rates compared to controls (Fig. 4), with 1.32-, 1.56-, 2.12-, 1.32-, and 1.67-fold increases for the moderately filtered set and 1.24-, 1.17-, 2.16-, 1.38-, and 1.58-fold increases for the stringently filtered set for A-phased repeats, direct repeats, G4s, inverted repeats, and mirror repeats, respectively (Fig. 2A; Table S3). Per-motif deletion error rates in Z-DNA motifs were significantly *reduced* compared to controls: 0.62-fold for the moderately filtered data and 0.24-fold for the stringently filtered data. The Poisson regression models for HiFi deletion error rates explained between 4.8% (Z-DNA motifs) and 12.0% (direct repeats) of deviance. Excluding the ‘non-B motif vs. control’ predictor led to a moderate reduction in percentage of deviance explained in models for direct repeats (17.5%) and mirror repeats (19.9%; Table S4). The presence of homopolymers was the most important predictor in all models; its removal led to the reduction of deviance explained between 57.9% and 91.7%.

#### ONT

The average per-motif deletion error rates were significantly different for all non-B motif types vs. controls for the moderately filtered set, and for G4s, inverted repeats, mirror repeats, and Z-DNA motifs for the stringently filtered set (Fig. 4). Sizable and consistent elevation in per-motif deletion rates was observed for G4 motifs over controls (1.64-fold and 1.53-fold for the moderately and stringently filtered sets, respectively). Differences in deletion rates between motifs and controls were smaller in magnitude for mirror repeats, Z-DNA motifs, and inverted repeats (from 0.99- to 1.19-fold). The explanatory power of the Poisson regression models explaining ONT deletion error rates ranged from 8.6% (for A-phased repeats) to 23.7% (for G4 motifs). The removal of the ‘non-B motif vs. control’ predictor led to a considerable reduction of deviance explained in a model for Z-DNA motifs (18.9%), whereas the removal of the predictor denoting the presence of a homopolymer led to the reduction in deviance explained by 68.6%, 53.8%, 61.8%, 34.8%, and 24.9% for A-phased, direct, inverted, mirror repeats, and Z-DNA motifs, respectively, but only 0.7% for G4 motifs.

### Insertion errors

#### Illumina

For Illumina, the insertion error rate was low and comparable to the deletion error rate (Table S2B). The per-motif insertion error rates were significantly and sizably elevated in all non-B motif types compared to controls for the moderately filtered set, and for A-phased repeats, direct repeats, and G4 motifs for the stringently filtered set (Fig. 5, Table S2A). A-phased repeats, direct repeats, and G4 motifs exhibited 1.50-, 29.2-, and 8.82-fold increase in insertion error rates in the moderately filtered, and 4.78-, 3.65-, and 7.15-fold increase in the stringently filtered set, respectively. In the Poisson regression models, the percentage of deviance explained ranged from 1.00% (for A-phased repeats) to 21.5% (for Z-DNA motifs). Importantly, the contribution of the ‘non-B motif vs. control’ predictor was significant in models for all non-B types, and its removal led to a reduction of the deviance explained by 21.6%, 47.7%, 85.2%, 39.5%, and 83.8% in models for A-phased repeats, direct repeats, inverted repeats, mirror repeats, and Z-DNA motifs, respectively, but only 1.50% for G4 motifs.

**Figure 5.**
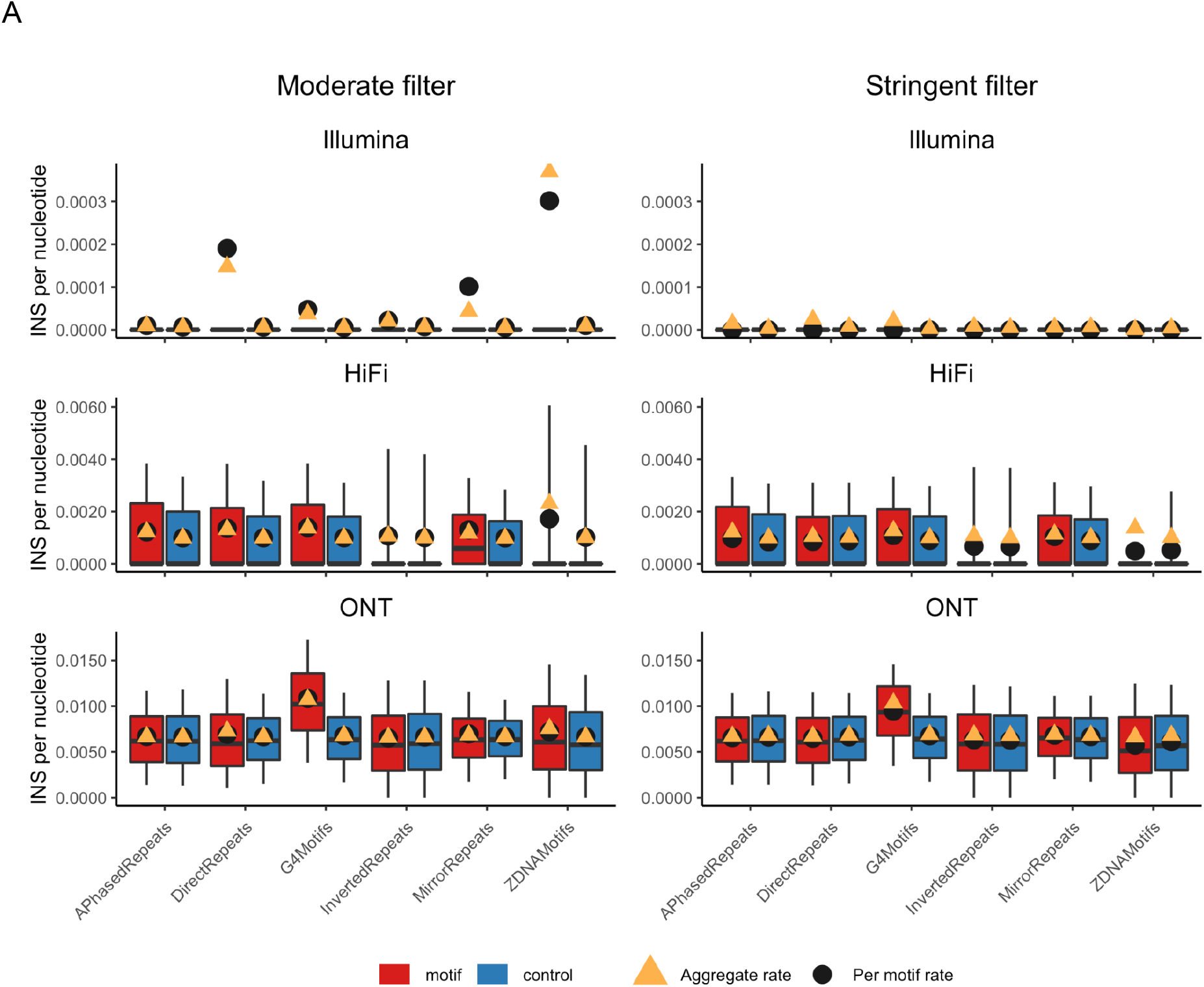
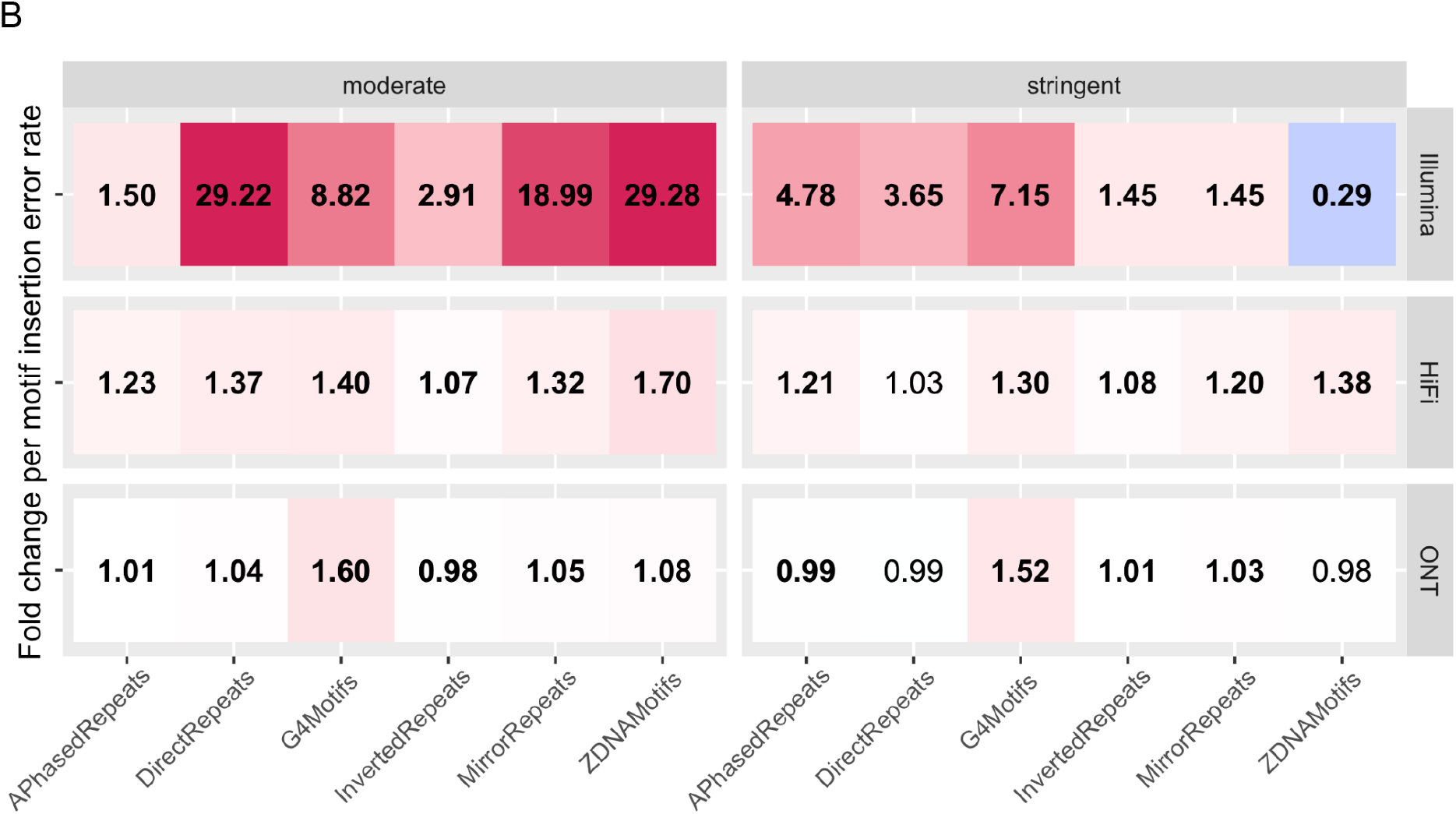
Insertion error rates in non-B motifs. **(A)** Boxplots of per-motif insertion error rates. Values above the 90th percentile are excluded from the boxplots in order to better visualize the bulk of the distributions. The left panel shows the moderately filtered motif set, the right panel the stringently filtered set, and the three rows correspond to the different technologies (Illumina, HiFi, and ONT). Red and blue boxes correspond to motifs and controls, respectively, black dots mark per-motif means, and orange triangles aggregate error rates (sum of all insertion errors divided by sum of all aligned nucleotides). Note that y-axes differ among technologies. **(B)** Heat maps visualizing fold changes in per-motif means of insertion error rates between motifs and corresponding controls. Red (green) shades indicate higher (lower) error rates in non-B motifs than in controls, with fold change values reported in each cell of the map. When these values are in bold, per-motif means were significantly different between motifs and controls (t-test p-values corrected for multiple testing smaller or equal to 0.05). Also here, left and right panels correspond to moderately and stringently filtered sets, respectively, and rows correspond to Illumina, HiFi, and ONT technologies, respectively.

#### PacBio HiFi

Similar to the deletion error rate, the insertion error rate for HiFi was approximately an order of magnitude higher than that for Illumina (Table S2B). Average per-motif insertion error rates for HiFi were significantly elevated in all non-B motif types compared to controls in the moderately filtered set, and in all non-B motifs but direct repeats in the stringently filtered set (Fig. 5; with 1.23-, 1.40-, 1.07-, 1.32-, and 1.70-fold increases for the moderately filtered set, and 1.21-, 1.30-, 1.08-, 1.20-, and 1.38-fold increases in A-phased repeats, G4 motifs, inverted repeats, mirror repeats, and Z-DNA motifs, respectively; Table S3). The explanatory power of the Poisson regression models did not exceed 5% in any of the models, rendering only limited information about the contribution of individual predictors (Table S4).

#### ONT

The insertion error rate in the ONT dataset was higher than for other technologies analyzed (Table S2B). Similar to the other technologies, for ONT we found significantly elevated per-motif insertion error rates in G4 motifs compared to controls (1.60- and 1.52-fold for the moderately and stringently filtered sets, respectively; Fig. 5). For the other non-B motifs, insertion error rates were similar in magnitude between motifs and controls (with fold-changes ranging between 0.98- and 1.08, Fig. 5), albeit often significantly different because of the large number of insertion events considered. Poisson regression models with ONT insertion error data explained between 1.60% (for inverted repeats) and 17.0% (for G4 motifs; Table S4) of deviance. The removal of the ‘non-B motif vs. control’ predictor led to a substantial reduction in deviance explained in the model for Z-DNA motifs (10.9%), and to <10% reduction in the other models.

### Sequencing depth and quality

For Illumina, we found lower average read depth in G4 and Z-DNA motifs than in controls (0.81- and 0.95-fold, respectively, in the moderately filtered set, and 0.82- and 0.90-fold in the stringently filtered set), and *higher* average read depth in A-phased, inverted, and mirror repeats than in controls (1.05-, 1.04- and 1.05-fold, respectively, in the moderately filtered set and 1.04-, 1.03- and 1.04-fold in the stringently filtered set; Fig. S3; Table S2A). Direct repeats did not affect average read depth. For HiFi, we observed only minimal differences in the aggregate read depth between non-B motifs and controls, ranging from 0.98- to 1.01-fold for the moderately filtered set and from 0.99 to 1.01-fold for the stringently filtered set (Fig. S3). For ONT, we found no relevant differences in aggregate read depth between non-B motifs and controls (Fig. S3, Table S3).

To evaluate potential effects of non-B DNA on sequencing base quality, we used the moderately filtered set. For Illumina, average base quality was lower in direct repeats, G4, and Z-DNA motifs with 0.99-, 0.97-, and 0.96-fold differences as compared to controls, respectively. Average base quality in A-phased repeats and inverted repeats motifs was slightly elevated as compared to that in controls (1.02- and 1.01-fold, respectively), whereas it was equivalent between mirror repeats motifs and controls (Fig. S4). For HiFi, average base quality was reduced in comparison to controls in direct repeats (0.99-fold) and Z-DNA motifs (0.96-fold), while it was increased in A-phased repeats (1.08-fold) and G4 motifs (1.08-fold). In inverted and mirror repeats motifs, average HiFi base quality was not different between motifs and controls. We could not measure the potential effects of non-B motifs on ONT base quality because no read base quality values were available in the data used (see Methods).

### False-positive single-nucleotide variants due to errors in non-B motifs

To evaluate the probability of identifying sequencing errors as variants (i.e., of false positive single-nucleotide variants, or SNVs) and to gauge the expected number of such false positives in motifs with non-B-forming potential, we developed a probabilistic model that takes into account several key parameters (see Methods). It incorporates the per-nucleotide error rates derived from the analyses presented above, as well as the number of haploid genomes, the average sequencing read depth per haploid genome, the minimum number of reads used to identify a variant, the minor variant frequency used to call an SNV, and the total number of base pairs covered by non-B motifs (either of a certain type or all together; see Methods for details).

We applied this model to three hypothetical scenarios with varying ‘sample sizes’, i.e. numbers of haploid genomes—200, 2,000, and 20,000 (corresponding to 100, 1,000, and 10,000 individuals)—and average read depths per haploid genome ranging from 3× to 30× (Fig. 6A). For each scenario, we estimated the expected number of false positive SNVs using the probabilistic model with the SNM error rates we computed above for the moderately filtered set in each of the three technologies. This was performed separately for motifs of each non-B type and for controls (B DNA), and considering different ways of scoring rare variants (i.e., by scoring variants present in a single haploid genome and three haploid genomes, and by using a minor variant frequency threshold of 0.01; Fig. 6A). Requiring a variant to occur in multiple (e.g. 3-5) haploid genomes is frequently used when studying rare variants (Wainschtein et al. 2022). For all three ways of scoring variants, the expected number of false positive SNVs is higher in all three technologies when considering non-B motifs of all types combined compared to an equally long stretch of B DNA sequence (solid and dashed lines, respectively, in Fig. 6A). However, above the read depth of 21× for Illumina and HiFi, the expected number of false positives approaches zero even for singletons in all three sample size scenarios. In contrast, ONT sequencing may not be advisable when investigating singleton variants due to its substantial expected false positives even at higher read depths. For tripletons and variants with a minor frequency above 0.01, expected false positives are much lower overall. Indeed, already at an average depth of 6× virtually no false positive SNVs are expected when requiring a minor frequency above 0.01 for sample sizes larger than 100 individuals in Illumina and HiFi (for ONT, this is achieved at ~9× read depth). Whereas motifs of all non-B DNA types were combined above, expected false positives computed separately for different non-B DNA types (Fig. S5) depend, for each non-B motif type, on its error rate and on the number of nucleotides it occupies in the genome. In addition to calculating expected false positives, we also ran Monte Carlo simulations with our probabilistic model to gauge the variability of false positive values under the different scenarios considered (see Methods)–drawing congruent conclusions (Fig. S6).

**Figure 6.**
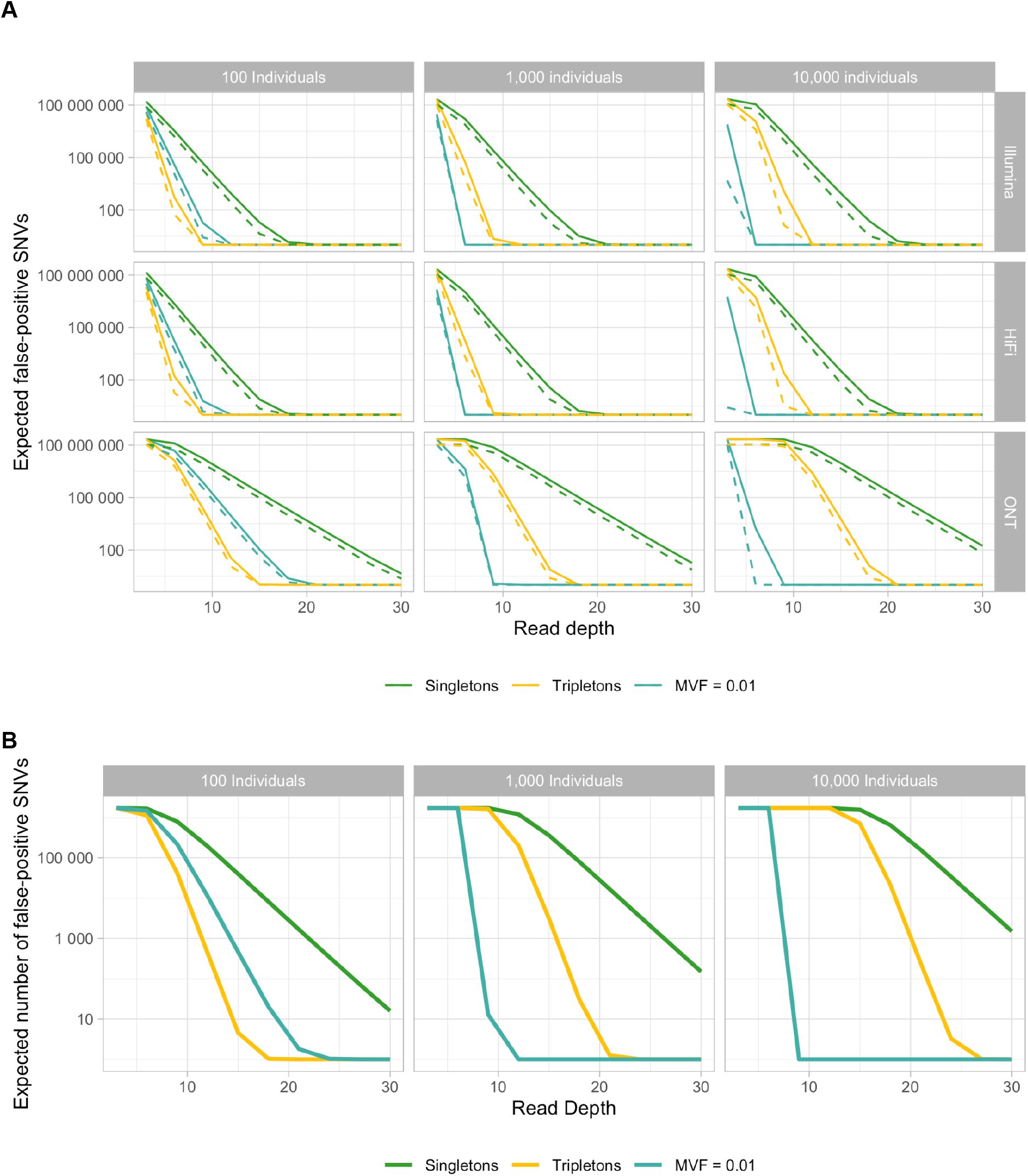
False-positive SNVs in non-B motifs. **(A)** Scenarios with different technologies corresponding to rows and numbers of diploid individuals (100, 100, and 10,000) corresponding to columns. The average read depth per haploid genome is plotted on the x-axis, whereas the expected number of false-positive SNVs due to errors is shown on the y-axis. Colors indicate different variant frequency filters (singletons, tripletons, and 1%), whereby the solid line corresponds to the cumulative number of all false-positive SNVs across non-B types, and the dashed line to the number of false-positive SNVs in an equally long stretch of B-DNA. **(B)** Expected false-positive SNVs in middle guanines in guanine-triplets at G4 motifs in Illumina sequencing. Within G4 motifs, there are 1,715,082 bp that fit the requirements of a middle guanine in a guanine triplet, which may have an extremely high error rate (Schirmer et al. 2016). Plotted are the numbers of expected false-positive SNVs, with the same coloring scheme and numbers of diploid individuals as (A).

We then investigated a special case of an exceptionally high error rate potentially occurring in G4 motifs, namely the middle guanines in guanine triplets. Schirmer et al. (Schirmer et al. 2016) have shown that, in Illumina sequencing, the second positions of guanine triplets exhibit error rates orders of magnitude higher than genome-wide averages. Using this drastically elevated error rate (0.035 errors per site) in our probabilistic model for all potential middle Gs in the G4 motifs analyzed in this study (a total of 1,715,082 bp), we obtain large expected false positive values even at higher read depths (Figure 6B) and at higher minor variant frequency cutoffs. Indeed, false positives for singletons do not approach zero even at sequencing depth of 30×, and they approach zero at the depth of 23× and 27× for tripletons scored for 1,000 and 10,000 individuals, respectively. This indicates that for analyses of variants contained in G4 motifs, substantially higher read depths and/or more stringent minor variant filters are necessary to discern between true and false positive variants.

To further illustrate the potential impact of non-B motifs on false positive SNVs, we used our probabilistic model with parameters derived from three publicly available sequencing data sets: the 1000 Genomes project (1000G) (1000 Genomes Project Consortium et al. 2015), the Simons Genome Diversity Project (SGDP) (Mallick et al. 2016), and the Genome Aggregation Database (gnomAD) (Karczewski et al. 2020). These data sets are all based on Illumina sequencing technology and have vastly different sample sizes as well as considerably different average sequencing read depths. In the 1000G example, which has an average read depth of 2× (1× per haploid genome) and a high number of individuals (5,008 haploid genomes), singleton and tripleton variants cannot be reliably distinguished from sequencing errors, while requiring minor variant frequencies ≥0.01 drastically reduces the expected number of false positives (Table S8). In contrast, in the SGDP example (21× read depth per haploid genome, 600 individuals), the sequencing depth is such that, according to our probabilistic model, false positive variants are not expected regardless of the minor variant frequency. In the gnomAD example (15× read depth per haploid genome, 152,312 haploid genomes), 11,044 errors are expected to be falsely identified as singletons among all non-B motifs, compared to 2,481 errors among an equally long stretch of B DNA. These numbers are dramatically reduced (leading to virtually no expected false positives according to our probabilistic model) in tripletons and variants with frequencies ≥0.01. Overall, these results are in line with previous findings on the relationship between read depth, minor variant frequency, and the occurrence of false positives. They highlight the need for sufficient depth and quality control in error-prone regions (Kishikawa et al. 2019; Tabangin, Woo, and Martin 2009), to which we now add non-B DNA, and suggest that investigations incorporating rare variants in such regions should be carried out with additional caution and sufficient read depth.

## Discussion

Identifying biases in sequencing accuracy and predicting sequencing success are paramount for genomic studies. By using publicly available data, we demonstrated that non-B-DNA-forming motifs are associated with altered sequencing success across three major sequencing technologies. Thus, such motifs should be taken into consideration when interpreting existing sequencing studies and designing new ones. As our Poisson models suggested, the association of non-B motifs with error rates can be caused by the co-occurrence of other attributes, such as biased nucleotide composition or the presence of homopolymers. Yet, previous studies suggested that a variety of non-B DNA structures are affecting the function of DNA polymerases *in vivo (Mirkin and Mirkin 2007)* and this might be the case also when these enzymes are employed in sequencing instruments.

### SNM rates

Overall, we found moderate associations between non-B motifs and SNM error rates. The largest was observed for G4 motifs, which exhibited consistently elevated SNM error rates across all technologies and both filtering schemes (Fig. 2). The magnitude of this elevation was lower for HiFi and ONT than for Illumina (Fig. 2). For HiFi, when we restricted attention to the non-repetitive portion of the genome through our ‘stringent’ filtering, we observed no significant differences in SNM error rates between non-B motifs and controls (Fig. 2). For Illumina, our Poisson regression models suggest that in some cases altered SNM error rates are mainly associated with the presence of non-B DNA motifs themselves (e.g., Z-DNA, direct repeats, and G4 motifs) and in others with their peculiar nucleotide composition (e.g., mirror repeats and A-phased repeats). The latter is in line with previous studies indicating that GC-content affects sequencing depth and errors in Illumina (Aird et al. 2011). We also found higher SNM error rates in spacers/loops than in repeat arms/stems, with particularly strong elevations for the Illumina technology and for G4 motifs (Fig. 3). This heterogeneity in SNM error rates within motifs should also be taken into account when analyzing variants located in non-B DNA. It suggests that, while average error rates over an entire non-B motif might be only slightly elevated compared to control regions, certain subregions within the motif may be more prone to sequencing errors than others. This was also evident when we analyzed the expected number of false positives at the middle Gs in G4s’ stems (Fig. 6B).

### Deletion error rates

The association of non-B motifs with deletion error rates was stronger than that with SNM error rates. Particularly for the Illumina technology, fold-differences between non-B motifs and controls were larger in magnitude for deletion than for SNM error rates. G4 motifs had elevated deletion error rates over controls across technologies and filtering schemes, with highest fold-increases for Illumina, intermediate for HiFi, and lowest for ONT. Our models indicated that, in addition to the presence of non-B motifs, deletion error rates are strongly associated with the presence of homopolymers. While ONT sequencing is known to show a bias in homopolymer regions (Bowden et al. 2019), the ongoing improvement of both base-calling algorithms and sequencing chemistry is expected to further reduce this bias in the future. Notably, while fold-differences in deletion error rates between non-B motifs and controls were smaller in magnitude for HiFi than for Illumina, and were smallest for ONT, both long-read technologies exhibited higher overall deletion error rates than the short-read Illumina.

### Insertion error rates

We found that, compared to deletion error rates, insertion error rates (albeit low) were more strongly affected by the presence of non-B motifs for Illumina, less for HiFi, and similarly largely unaffected for ONT. The fold-increases in insertion error rates due to the presence of non-B motifs for the Illumina technology were the largest across all the analyses we conducted. For this technology, even after applying stringent filtering, we found significant increases in insertion error rates. Again for Illumina, and for all non-B motif types but G4 motifs, our models indicated a sizable contribution of the motif presence to explaining variability in insertion error rates. These contributions were only moderate for HiFi, and minor for ONT.

### Effects of moderate vs. stringent filtering

We found that applying moderate vs. stringent filtering dramatically alters Z-DNA deletion and insertion error rates for the Illumina technology. When the moderate filter is applied, Z-DNA exhibits highly elevated deletion and insertion error rates compared to controls, while these rates are reduced when the stringent filter is applied. Given that in the stringently filtered data set any overlap with microsatellites is removed, we suspect that microsatellites are driving the signal in the moderately filtered data set. We also note that our reanalysis of the Simons Diversity Project data (Illumina sequencing) to compare diversity between B DNA and non-B DNA indicated G4 motifs stand out in terms of SNV diversity even when stringent filtering is applied (Fig. S7).

### Depth and quality

Across technologies, Illumina clearly showed the largest differences in average read depth between non-B motifs and controls. Importantly, read depth was reduced in G4 and Z-DNA motifs. For G4 motifs in particular, this reduction was sizable—20% and 18% decrease in sequencing depth compared with controls for the moderately and stringently filtered sets, respectively. Interestingly, read depth was hardly altered at all for the two long-read technologies HiFi and ONT. For Illumina and HiFi technologies, the associations between non-B motifs and sequencing quality appear to be minor.

### Conclusions and recommendations

Our results suggest that the relationship between non-B motifs and sequencing accuracy differs across technologies and sequencing error types. Therefore, the choice of technology depends on the type of errors one is trying to avoid at non-B motifs. To minimize single-nucleotide errors at non-B motifs, we recommend using PacBio HiFi, particularly for the non-repetitive portion of the genome. Both HiFi and ONT display low SNM error biases at non-B DNA motifs as compared to Illumina, however, HiFi has lower overall SNM error rates as compared to ONT. If minimizing deletion and insertion error biases at non-B motifs is of interest, the choice should be between HiFi and ONT, which exhibit comparatively lower biases at non-B motifs, with a preference towards HiFi which has overall low indel rates. Illumina has low insertion and deletion error rates, but its insertion and deletion error biases at non-B motifs are substantial, making it a suboptimal choice.

If one is to choose one sequencing technology to obtain the most accurate results for non-B motifs minimizing all three types of sequencing errors, we would recommend HiFi, which balances out low error rates with relatively low biases at non-B motifs. The ONT technology, while having higher overall rates for all errors considered, does not appear to be affected by non-B DNA motifs as much as Illumina and PacBio HiFi, indicating that ONT may carry less bias in non-B motifs. This is consistent with the fact that this technology is not polymerase-based (Daniel and Deamer 2019). However, G4 motifs are the only type of non-B that exhibit a considerable increase of mismatch errors–which is somewhat surprising since the helicase employed in ONT sequencers should in principle be less susceptible to structures formed in single-stranded DNA (G4 motifs and cruciforms). Overall, given a sufficient read depth, ONT might be less prone to false positive variants at non-B motifs, which, in addition to its ability to generate ultra-long sequencing reads (M. Jain et al. 2018), makes it attractive.

The use of multiple technologies facilitates error detection for any type of mutation, since true variants should be present in all technologies, each with its own sequencing error profile. In addition to combining different technologies, our calculation of expected false positive SNVs based on SNM error rates in non-B motifs suggests that read depth and variant frequency cutoffs are critical–especially for rare variants. The high amount of expected false positive singleton variants in scenarios with low read depth, and particularly in subregions of G4 motifs, highlights the importance of treating these regions with extra caution in downstream analyses, and of applying rigorous quality filters. Moreover, results obtained from our probabilistic model with parameters derived from the 1000Genomes, SGDP, and gnomAD datasets highlight how investigating rare variants with low read depth may be particularly problematic. In some extreme cases (for example middle guanines in guanine triplets), with insufficient read depth variants may become indistinguishable from errors if they are only occurring once or only a few times among studied individuals.

Overall, in cases where abundant read depth is available, the increased error rate in non-B motifs might not lead to mistaking errors for biological variants (i.e., to false positives), especially when stringent filters are applied in terms of read quality and the avoidance of repetitive regions. However, when read depth is low (e.g. in ancient DNA or pooled population sample studies), variants identified in non-B motifs require additional scrutiny before being used in downstream analyses, and it may be advisable to restrict attention to variants identified by multiple technologies (when available) to avoid technology-specific biases.

## Methods

### Data

We used publicly available sequencing data generated for the Genome in a Bottle (GIAB) Consortium ((Zook et al. 2016), https://www.nist.gov/programs-projects/genome-bottle). We downloaded Illumina, Pacbio HiFi, and ONT data for one individual—the son of the Ashkenazim trio (HG002, NA24385). We downsampled the existing Illumina (2×150 bp, generated with Illumina HiSeq 2500 Rapid SBS) alignment file of 300× to ~100×. For PacBio HiFi, we downloaded 167 Gb of raw read data (30× consensus read depth, generated with the PacBio Sequel instrument). Since the ONT base-calling is under constant development, we used a dataset available at https://nanoporetech.github.io/ont-open-datasets/, which employs a significantly improved base-calling algorithm (Bonito v0.3.0) on the same raw dataset with sequencing depth of 57× (HG002, NA24385).

### Read mapping

In all our analyses, we used hg19 as a reference, as the most comprehensive genomic annotations and other resources are available for this version. For Illumina, we downloaded alignment files from the GIAB homepage (https://github.com/genome-in-a-bottle/giab_data_indexes). For PacBio and ONT, we aligned reads to hg19 using *minimap2* with the Pacbio HiFi and ONT specific parameters, respectively (Li 2018). In all alignment files, we removed duplicates using *picard tools* (http://broadinstitute.github.io/picard/), and sorted and split reads into forward and reverse and by chromosome using *samtools* (Li et al. 2009).

### Non-B DNA annotations

For all non-B-DNA motifs except G4 motifs, we used annotations available at the non-B DNA Database (https://nonb-abcc.ncifcrf.gov), which are based on the human reference hg19. To predict potentially G4-forming loci, we used *Quadron (Sahakyan et al. 2017),* which provides predicted stability values for each motif, with default parameters. We then downloaded the mappability track for hg19 based on 36-mers from the UCSC Genome Browser (http://genome.ucsc.edu/), used the R package *genomicRanges* (Lee and Schatz 2012; Lawrence et al. 2013) to calculate mean mappability values, and used *bedtools nuc* (Quinlan and Hall 2010) to obtain nucleotide composition (abundance of each of the four bases) for all motifs. For base quality, we first obtained values for each read and position within each motif using *samtools mpileup* (version 1.9, (Li et al. 2009)) and then calculated the average for each motif. Since Illumina and PacBio technologies use different quality score encoding, these results are not directly comparable. For the ONT dataset, there are no quality scores available in base-called read data.

For each non-B motif type, we removed overlapping motifs (within the same non-B type), and those overlapping with a nucleotide homopolymer ≥7 bp, as well as any motif larger than 1,000 bp or with an average mappability lower than one. In addition, we recorded any overlap with a motif of another non-B type, a RepeatMasker annotation, a homopolymer ≤7 bp, or an annotated microsatellite. Microsatellite annotations were generated with STR-FM (Fungtammasan et al. 2015), identifying mono-, di-, tri-, and tetranucleotide repeats with a copy number of at least seven units. This set of non-B motifs, which we call the ‘moderately filtered set’, formed the basis for randomly generating control regions: independently for each non-B type, we constructed controls matching the number and the size of the motifs, and excluding reference gaps and all non-B-DNA motifs. For all control regions, we gathered features such as base quality, nucleotide composition, overlap with RepeatMasker annotations, etc. To form the ‘stringently filtered set’, we first excluded any motif that overlapped with a RepeatMasker or an STR annotation and any motif that either overlapped with, or was within 50 bp of, another non-B motif. Additionally, we excluded any motif with Phred quality score lower or equal to 30 (Illumina) or 73.17 (HiFi). To adjust controls, we subjected them to the same filters and then randomly subsampled them to match the size of the corresponding motif sets.

To assess variation in error rate within non-B motifs, we further partitioned annotated motifs into subregions. For A-phased, direct, inverted, and mirror repeats, we split the annotations into repeat arms and spacers, and for G4 motifs—into stems (the G-tract) and loops. To be consistent across motifs, we restricted analyses on subregions to G4 motifs with four stems and three loops, and to repeat motifs (A-phased, direct, inverted, and mirror repeats) with two repeat arms and one spacer.

### Error calling

Since variant detection tools are usually focused on detecting true biological variation and are optimized to avoid sequencing errors, we developed a script that, for each region of interest, counts the total number of aligned nucleotides and naively identifies mismatches directly from the CIGAR string of an alignment file. Any single-nucleotide, insertion, or deletion mismatch is recorded, unless it overlaps with a true biological variant present in the true variant set for HG002 (Zook et al. 2016).

To add an orthogonal approach in single-nucleotide error calling for Illumina, we also detected SNV sequencing errors by examining the overlaps between mates in Illumina read pairs as described in Stoler, Nekrutenko (2021), using the full (300×) data set described above. For each read pair, we restricted our analysis only to the errors located in the region between 50 and 60% of the full read length. For this analysis, we used the same sets of non-B motifs and controls as described above.

### Read depth and base quality

To assess potential biases in sequence read depth, we calculated the mean depth per bp (total number of aligned nucleotides divided by total length) and per motif (averaging the mean depth per bp across all motifs) for all non-B motif types and associated controls. Likewise, we computed mean base quality for motifs and controls for Illumina and HiFi sequencing data.

### Downstream statistical analysis

All statistical analyses were performed separately on the ‘moderately’ and ‘stringently’ filtered sets described above. T-test p-values for the comparisons between the average per-motif and per-control mismatch error rates were adjusted for multiple testing using the Benjamini-Hochberg correction (Benjamini and Hochberg 1995).

To study the effect of non-B-forming motifs on sequencing errors while taking into consideration the effects of other quantities such as nucleotide composition, motif length, and the presence of homopolymers, we fitted a Poisson regression model using the *glm* function in R with Poisson distribution and log link function. This model was chosen because errors, i.e. our responses, displayed extremely right-skewed distributions with an excess of zero. Specifically, we used the counts of single nucleotides mismatches, deletions, and insertions as responses in separate regressions, always including the logarithm of the total number of sequenced nucleotides as an offset term, i.e. a predictor with fixed coefficient equal to 1, to control for motif length and sequencing depth. In symbols, we employed the Poisson model

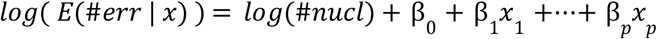

where *#err* is the error count, *#nucl* is the total number of sequenced nucleotides and *x*_1_,…,*x_p_* are the predictors. This model is mathematically equivalent to using the error rates as responses:

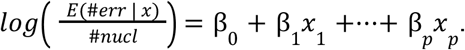

Nucleotide composition, as represented by the proportions of A, C, T, and G nucleotides in each motif or control, was transformed using the isometric log-ratio transform of Aitchison for compositional data (Aitchison 1982). For each regression, after fitting the full model with all predictors included, we excluded influential outliers (i.e. observations with a Cook’s distance >1, (Läuter 1985), fit the model again, and computed the share of deviance explained (*D_0_* - *D_m_*)/*D_0_*, in which *D_0_* is the null deviance, and *D_m_* the residual variance of the model. To estimate the contribution of each individual predictor, we then repeated the fit, excluding from the model one predictor at a time and calculating the reduction in the share of deviance explained (Kelkar et al. 2011); in symbols, ([*D_0_* − *D_m_*] − [*D_0_* − *D*_*m*{-}_])/(*D_0_* − *D_m_*), where *D_m_* and *D_m{-}_* are the residual deviances of the full and reduced model, respectively. All statistical analyses were performed with the R programming language and figures were produced with the *ggplot2* package *(Wickham 2011).*

### Probabilistic model for false-positive SNVs

We built a probabilistic model to quantify the contribution of sequencing errors in variant identification and to estimate the effect of non-B motifs on the number of false-positive SNVs called in several scenarios. Specifically, we modeled the presence of a sequencing error at a single site of a certain type (for example, a site belonging to a certain type of non-B motif) using the Bernoulli distribution, i.e. *X* ~ *B*(1, *r*) where *r* is the corresponding per-nucleotide sequencing error rate. Assuming independence among errors of different sequenced nucleotides mapping to the same site, the number of errors observed in a site sequenced at depth *d* is *Y* ~ *B*(*d*, *r*). Further assuming that all the sequencing errors in a site generate the same variant (worst case scenario), the probability of wrongly identifying a variant in a site of an haploid genome is *p_var_* = *P*(*Y* ≥ *min_reads_*) where *min_reads_* is the minimum number of reads required to call a variant. Finally, considering the total number of haploid genomes *g,* and assuming independence among their sequencing errors, we obtain that the number of haploid genomes with a variant in a site due to sequencing errors can be modeled as *V* ~ *B*(*g*, *p_var_*). Hence, the probability of identifying a variant in a site due to sequencing errors is *p_SNV_* = *P*(*V* ≥ *min_var_*), where *min_var_* is the minor variant frequency to call an SNV.

Using such probability *p_SNV_* and the total number of sites of the considered type, and assuming constant sequencing depth at all sites, we can compute the expected number of false positive SNVs as *p_SNV_*·*n_sites_*.

Since this calculation only provides an expected number (an average) of false positive SNVs, we also implemented a Monte-Carlo simulation study based on the same generative model to evaluate the corresponding variability in false positive values under various scenarios.

## Code Availability

All custom scripts used in our analyses are available on GitHub: https://github.com/makovalab-psu/nonB-Seq-Errors.

## Acknowledgments

This project was supported by NIH grants R01GM136684 (to KDM) and R01CA23715 (to KAE).

## Supplementary Figures

**Figure S1.**
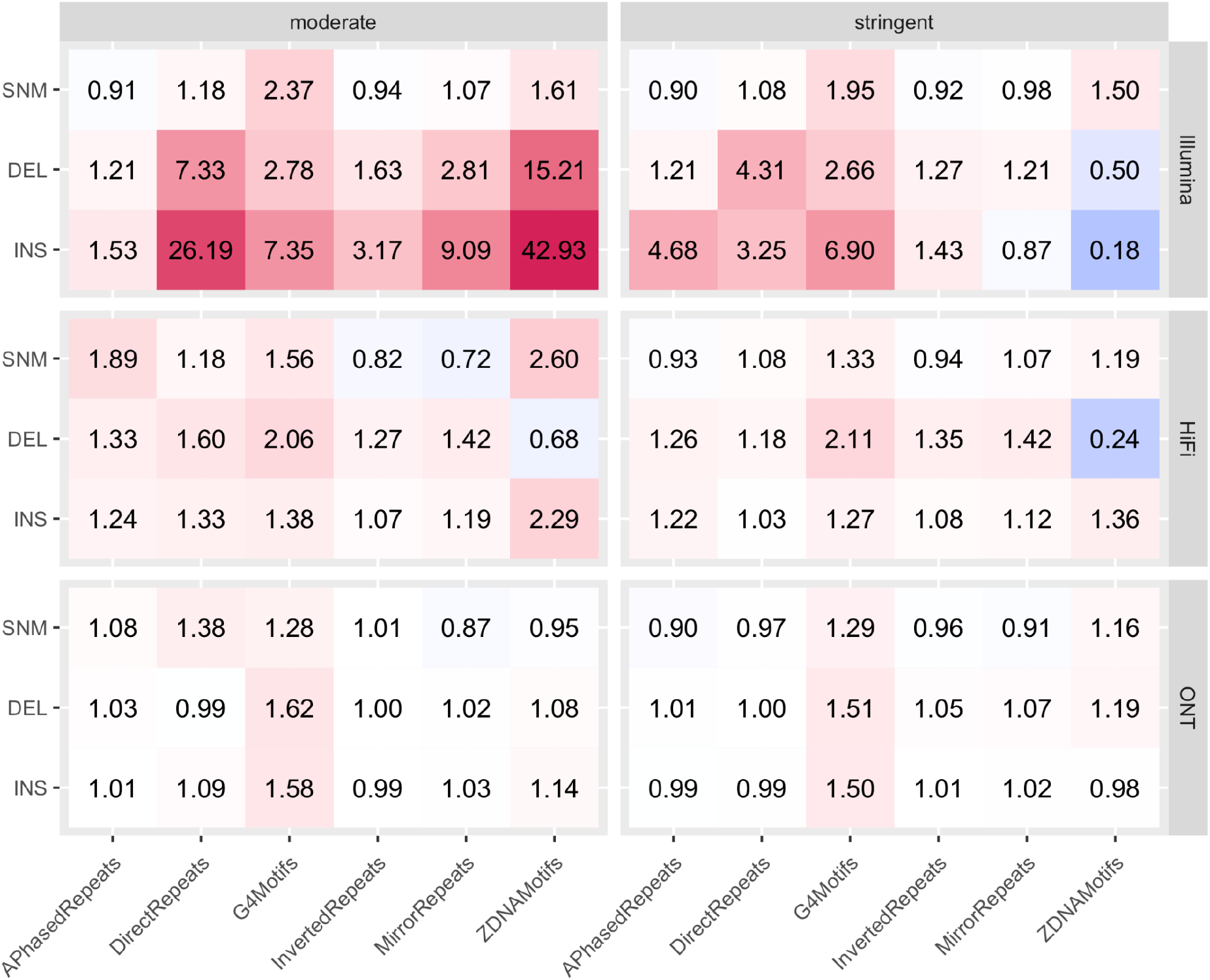
Heatmap of fold changes in aggregate error rates. Fold changes of different aggregate error rates (total number of mismatches divided by total number of aligned nucleotides) for single-nucleotide (SNM), insertion (INS), and deletion (DEL) errors. The left and right panels correspond to the moderately and stringently filtered motif sets, respectively, whereas rows correspond to Illumina, HiFi, and ONT technology. Shades of red and green indicate higher and lower error rate in non-B motifs than in controls, respectively. Note that these values are based on the aggregate, and not per-motif (and per-control) error rates (the latter are shown in the main manuscript).

**Figure S2.**
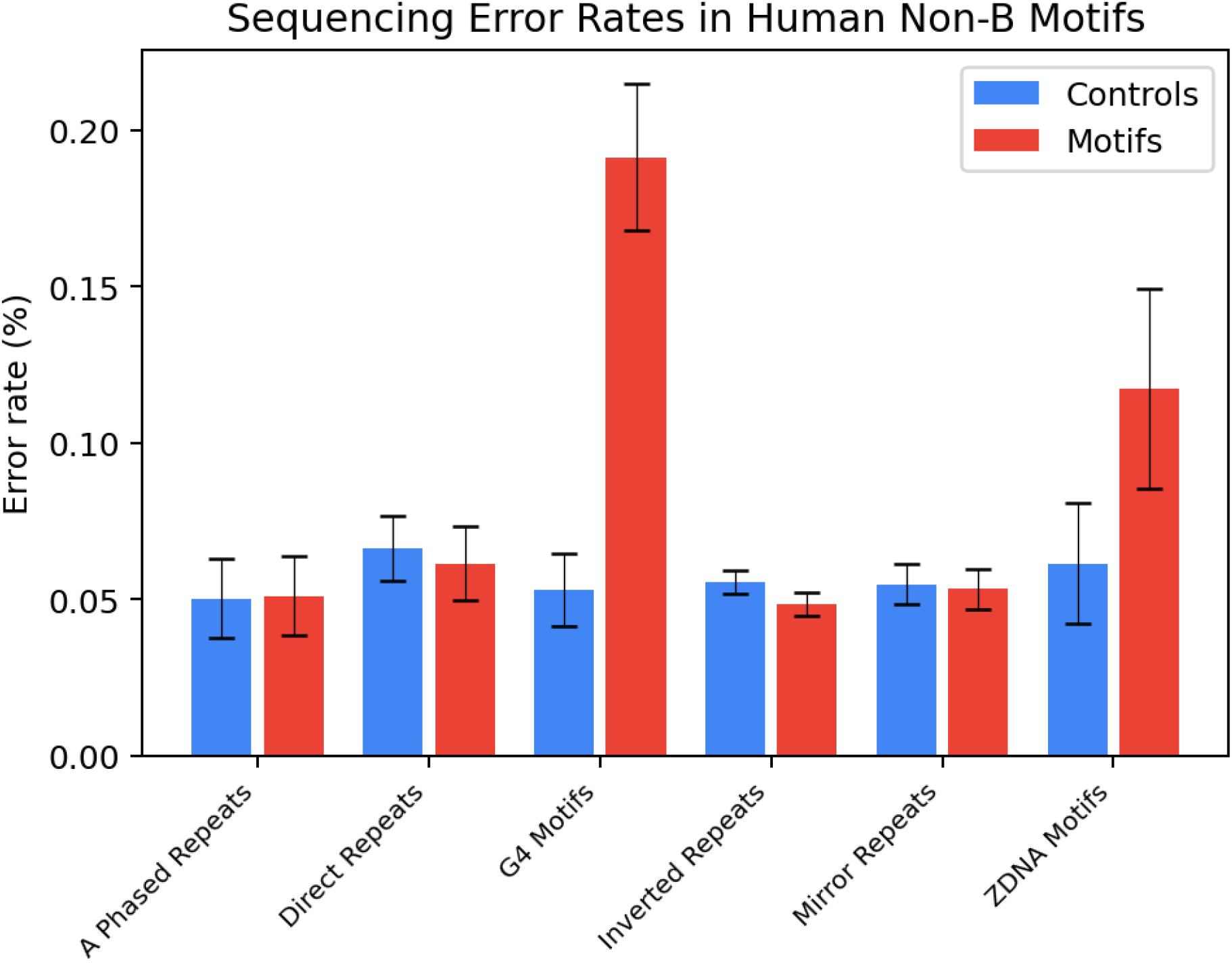
Single-nucleotide mismatch error rates in overlapping regions of paired-end reads. Shown are boxplots of SNM error rates derived from mismatches between pairs in overlapping Illumina read pairs. 95% confidence intervals were computed using the normal approximation of the binomial. To calculate the SNM error rate, the total number of mismatches was divided by the total number of nucleotides in the overlap. To minimize the effect of position in the read on the error rate, only the middle 10% of the reads were used. See Methods and Supplementary Table S5 for details.

**Figure S3.**
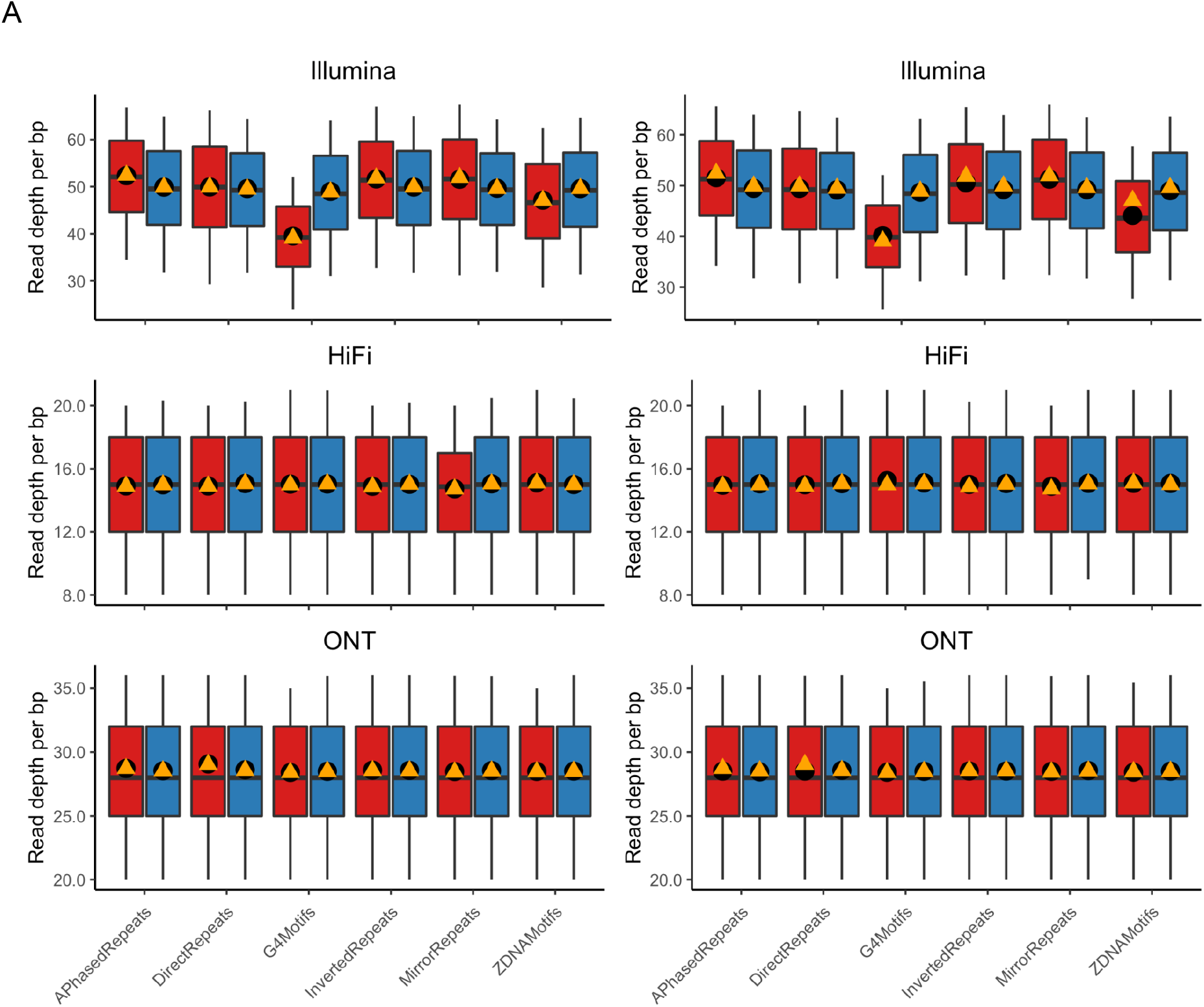
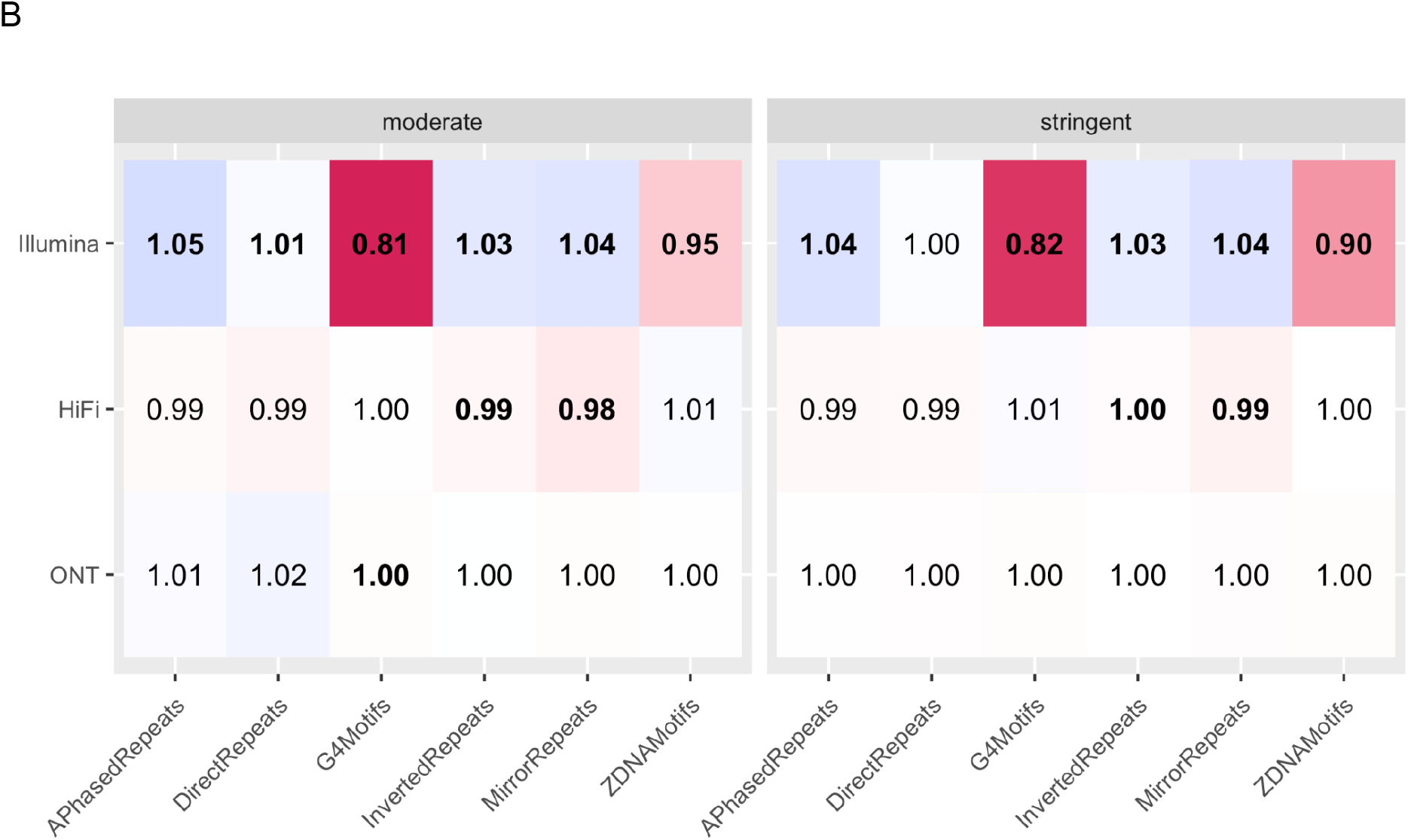
Read depth in non-B motifs. **(A)** Boxplots of per-bp read depth. Values above the 90th percentile were excluded from the figure to enable better visualization. The left panel shows results for the moderately filtered motif set, the right panel for the stringently filtered set, and the three rows correspond to the different technologies (Illumina, HiFi, and ONT). Black dots show values for per-motif means, orange triangles show overall error rates (sum of all aligned nucleotides divided by the total length of motifs / controls). **(B)** Heat map plot with fold changes of per-motif means of read depth. Red shades indicate lower values in non-B motifs than in controls, green shades indicate higher read depths in non-B motifs than in controls, with values also printed in rectangles. Values in bold represent fold-changes for which per-motif means were significantly different between motif and control (t-test p-values corrected for multiple testing). Left and right panels correspond to moderately and stringently filtered motif sets, respectively, and rows correspond to Illumina, HiFi, and ONT technologies.

**Figure S4.**
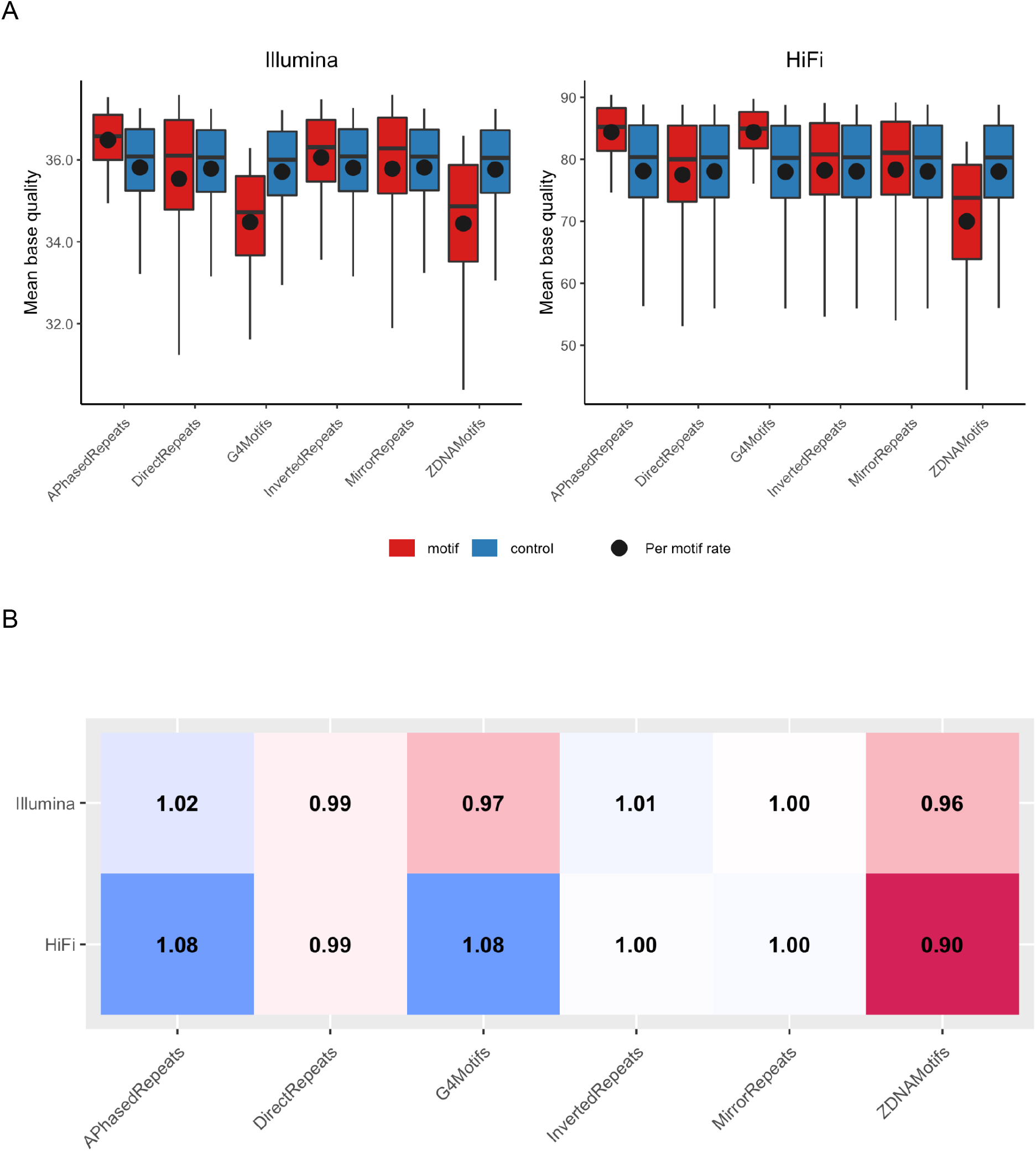
Base quality in non-B motifs. **(A)** Boxplots of average base quality. Values above the 90th percentile are excluded from the figure. Panels show results for Illumina and HiFi, respectively, and black dots show values for per-motif means. **(B)** Heat map plot with fold changes of per-motif means of base quality. Red shades indicate lower values in non-B motifs than in controls, green shades indicate higher base quality in non-B motifs than in controls, with values also printed in rectangles. Values in bold represent fold-changes for which per-motif means were significantly different between motif and control (t-test p-values corrected for multiple testing). Since we filtered for base quality in the stringently filtered motif set, we only show the results for the moderately filtered set.

**Figure S5.**
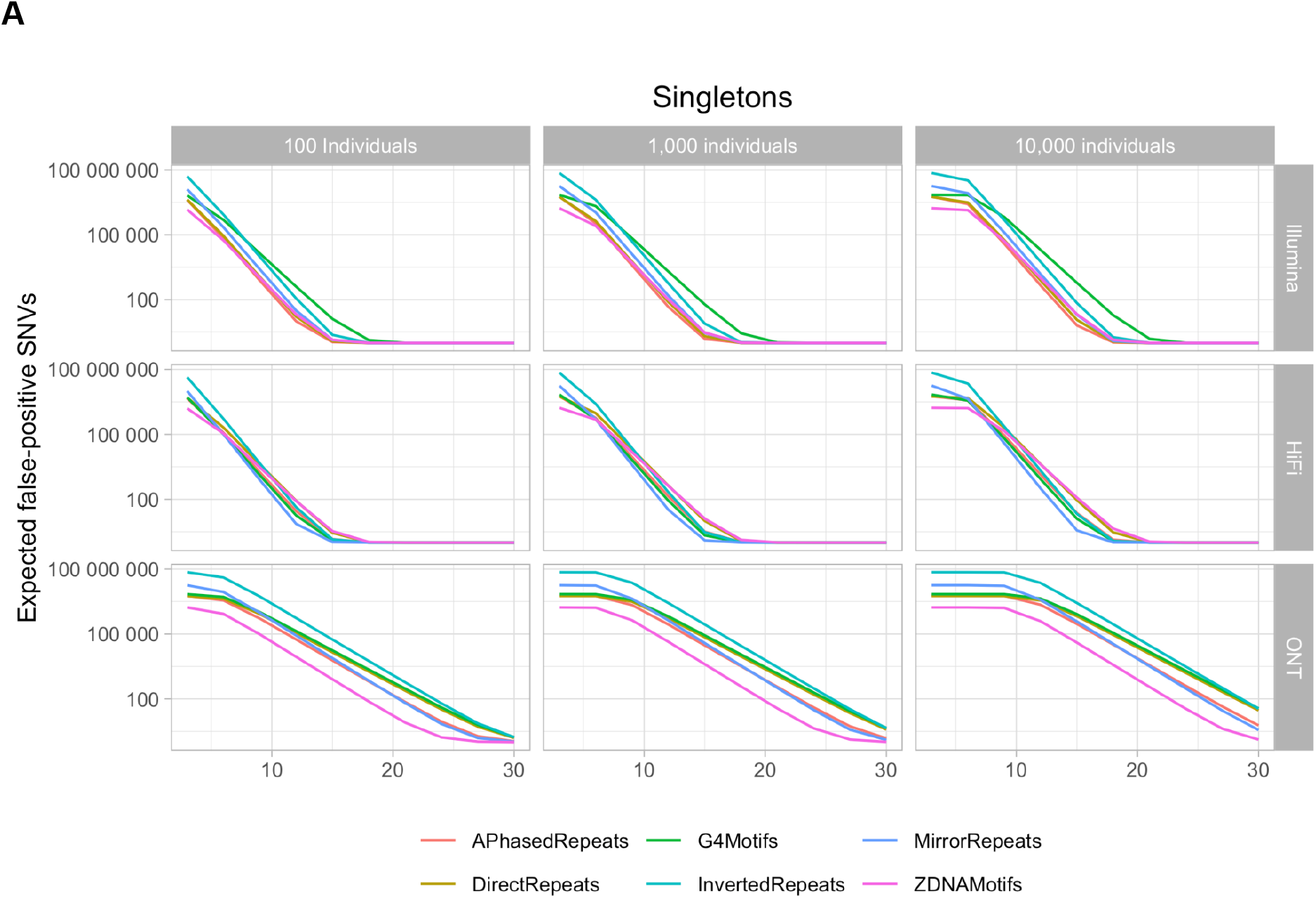
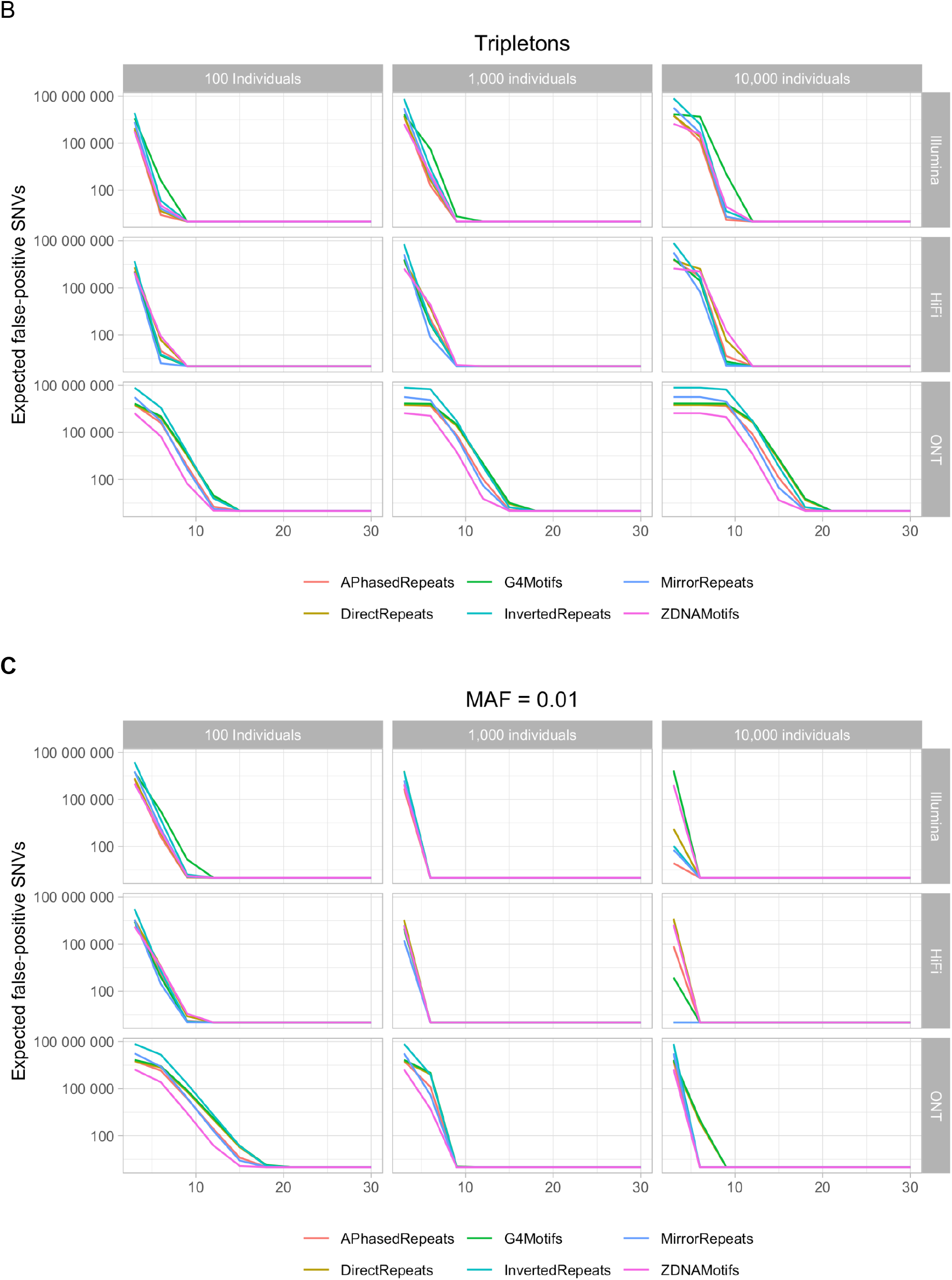
False-positive SNVs in non-B motifs. Shown are numbers of expected false-positive SNVs (y-axis) based on the probabilistic model and the SNM error rates derived from non-B motifs, for different haploid read depths (x-axis). For this analysis, we considered all bases annotated as a non-B motif of a particular type in the genome, i.e. the number of bases for each motif type differs among motif types. Different non-B motif types are shown in different colors. Columns correspond to different numbers of individuals, rows to the three technologies. (A), (B), and (C) show results for singleton, tripleton and 1% variant allele frequency cutoffs, respectively.

**Figure S6.**
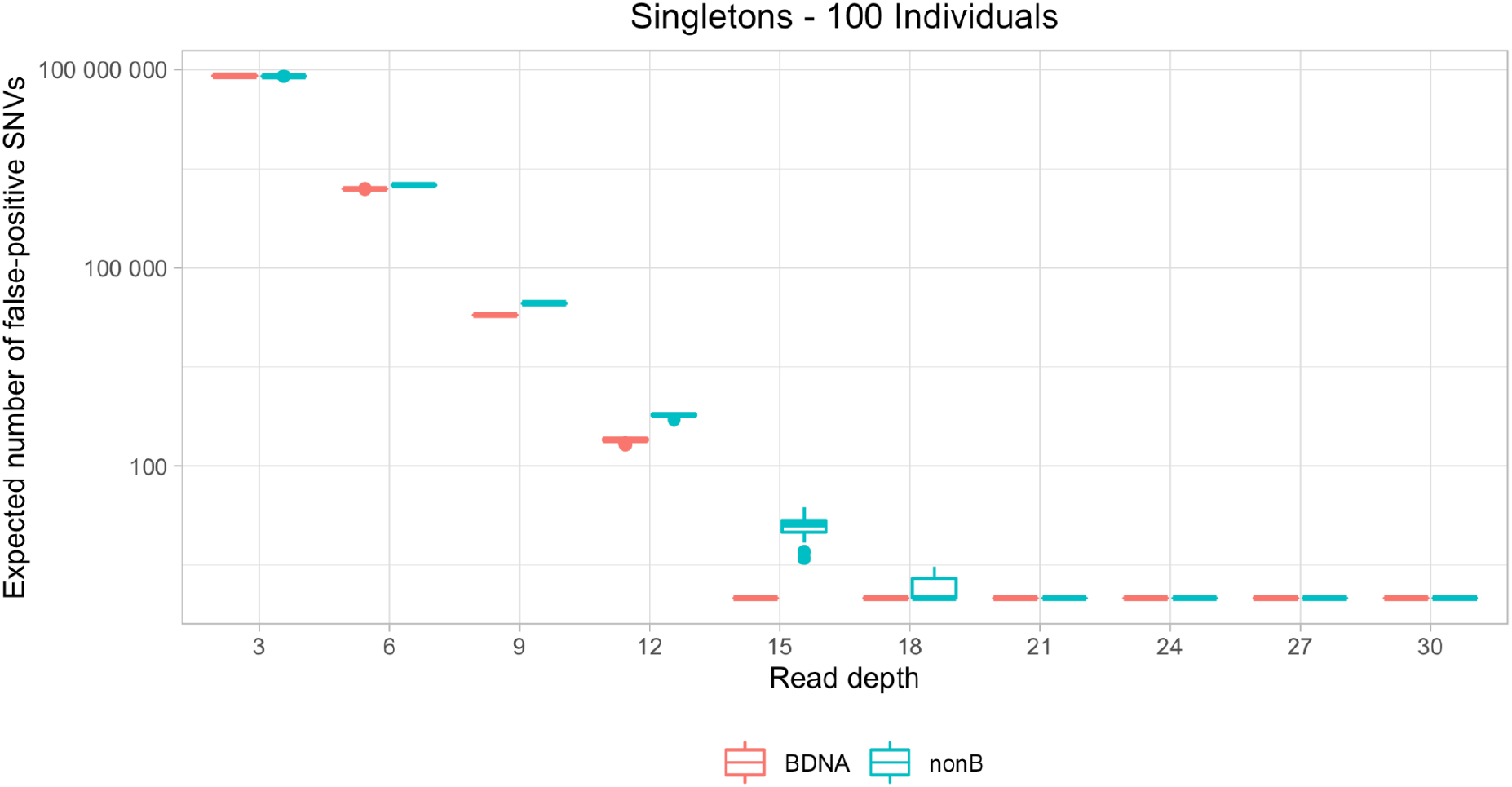
False-positive SNVs in non-B motifs, example of simulation results. Shown are numbers of expected false-positive SNVs in non-B motifs (red), and controls (blue), based on error rates in Illumina sequencing, and a minor variant frequency cutoff of one variant (singletons) in 100 diploid individuals. As opposed to results shown in Fig. 6, which only included the expected number of false positives, the boxplots of false positive values presented here are based on 100 Monte-Carlo simulations of each site.

**Figure S7.**
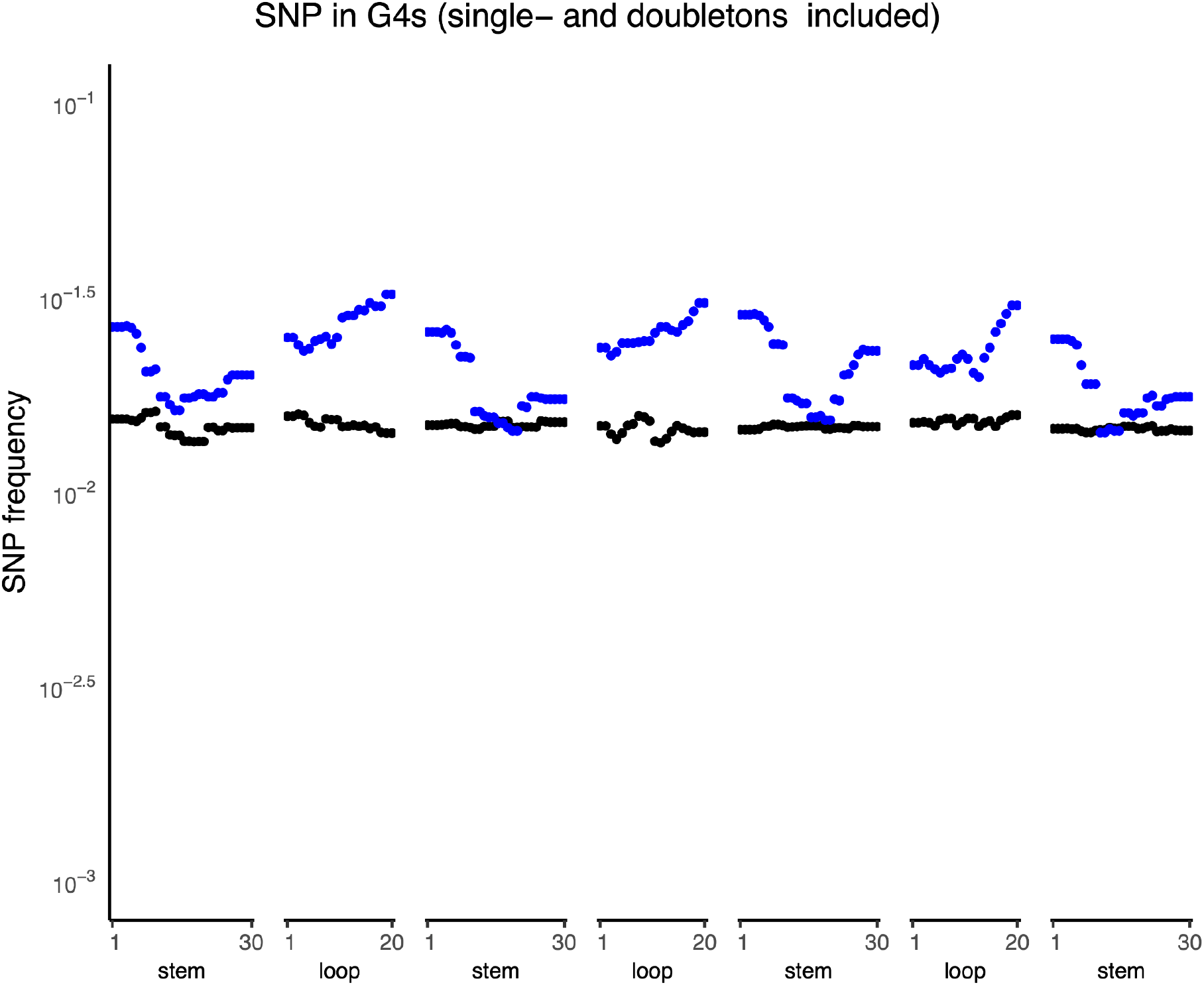
Genome-wide nucleotide substitution frequencies at G4 motifs and corresponding controls (for the stringent filtered data). The positions of nucleotide substitutions within motifs were scaled based on motif size. Stems are runs of guanines and loops are unspecified nucleotides between stems. For clarity of visualization, the Y-axis is displayed on a log scale. A comparison between all G4 loci (shown in blue) and control sequences (shown in black) for single-nucleotide polymorphism (SNP) frequencies based on the Simons Genome Diversity Project (Mallick et al. 2016) analyzed in (Guiblet et al. 2021).

