## Supplementary Tables for "Distinct sequencing success at non-B-DNA motifs"

**Table S1. Non-B motifs utilized in the study.** Shown are total motif set length, mean and median individual motif length, and number (n) of motifs for each non-B DNA type analyzed. Annotations are for the human reference genome *hg19*.

**Moderately filtered data set**

| Type | Total length (Mb) | Mean length (bp) | Median length (bp) | n |
|---|---|---|---|---|
| APhasedRepeats | 6.8 | 26.00 | 25 | 261,798 |
| DirectRepeats | 10.19 | 28.29 | 26 | 360,335 |
| G4Motifs | 8.59 | 35.37 | 33 | 243,017 |
| InvertedRepeats | 79.67 | 21.11 | 15 | 3,774,639 |
| MirrorRepeats | 29 | 58.25 | 51 | 498,033 |
| ZDNAMotifs | 2.76 | 12.44 | 11 | 222,534 |
| Total | 137.01 | | | 5,360,356 |

**Stringently filtered data set: Illumina**

| Type | Total length (Mb) | Mean length (bp) | Median length (bp) | n |
|---|---|---|---|---|
| APhasedRepeats | 1.00 | 25.76 | 25 | 38,851 |
| DirectRepeats | 0.77 | 28.30 | 27 | 27,093 |
| G4Motifs | 0.89 | 32.86 | 31 | 27,194 |
| InvertedRepeats | 9.84 | 16.59 | 15 | 592,876 |
| MirrorRepeats | 0.90 | 46.90 | 40 | 19,228 |
| ZDNAMotifs | 0.06 | 10.61 | 10 | 5,311 |
| Total | 13.46 | | | 710,553 |

**Stringently filtered data set: HiFi**

| Type | Total length (Mb) | Mean length (bp) | Median length (bp) | n |
|---|---|---|---|---|
| APhasedRepeats | 0.97 | 25.76 | 25 | 37,802 |
| DirectRepeats | 0.60 | 28.33 | 27 | 21,103 |
| G4Motifs | 0.90 | 32.84 | 31 | 27,320 |
| InvertedRepeats | 7.73 | 16.59 | 15 | 465,777 |
| MirrorRepeats | 0.72 | 47.04 | 40 | 15,294 |
| ZDNAMotifs | 0.03 | 10.61 | 10 | 2,921 |
| Total | 10.95 | | | 570,217 |

**Stringently filtered data set: ONT**

| Type | Total length (Mb) | Mean length (bp) | Median length (bp) | n |
|---|---|---|---|---|
| APhasedRepeats | 1.01 | 25.76 | 25 | 39,259 |
| DirectRepeats | 0.80 | 28.30 | 27 | 28,224 |
| G4Motifs | 0.92 | 32.85 | 31 | 28,058 |
| InvertedRepeats | 9.97 | 16.59 | 15 | 600,584 |
| MirrorRepeats | 0.93 | 46.90 | 40 | 19,778 |
| ZDNAMotifs | 0.06 | 10.61 | 10 | 5,576 |
| Total | 13.68 | | | 721,479 |

**Table S2A. Sequencing success measures in non-B motifs and controls.** In each row, aggregate (total number of errors divided by total number of nucleotides) and per-motif (or per-control) means of SNM rates, insertion rates, deletion rates, sequencing depth, and sequencing quality are shown for each non-B type, motifs and controls, technology, and filtering level.

| Non-B type | Source | Measure of sequencing success | Technology | Filter level | Aggregate mean | Per-motif (or per-control) mean |
|---|---|---|---|---|---|---|
| APhasedRepeats | motif | Single-nucleotide mismatches | Illumina | moderate | 0.00199 | 0.00204 |
| APhasedRepeats | control | Single-nucleotide mismatches | Illumina | moderate | 0.00219 | 0.00228 |
| DirectRepeats | motif | Single-nucleotide mismatches | Illumina | moderate | 0.00256 | 0.00308 |
| DirectRepeats | control | Single-nucleotide mismatches | Illumina | moderate | 0.00217 | 0.00228 |
| G4Motifs | motif | Single-nucleotide mismatches | Illumina | moderate | 0.00535 | 0.00567 |
| G4Motifs | control | Single-nucleotide mismatches | Illumina | moderate | 0.00225 | 0.00238 |
| InvertedRepeats | motif | Single-nucleotide mismatches | Illumina | moderate | 0.00205 | 0.00214 |
| InvertedRepeats | control | Single-nucleotide mismatches | Illumina | moderate | 0.00217 | 0.00231 |
| MirrorRepeats | motif | Single-nucleotide mismatches | Illumina | moderate | 0.00225 | 0.00269 |
| MirrorRepeats | control | Single-nucleotide mismatches | Illumina | moderate | 0.00210 | 0.00223 |
| ZDNAMotifs | motif | Single-nucleotide mismatches | Illumina | moderate | 0.00363 | 0.00390 |
| ZDNAMotifs | control | Single-nucleotide mismatches | Illumina | moderate | 0.00226 | 0.00238 |
| APhasedRepeats | motif | Insertion mismatches | Illumina | moderate | 0.00001 | 0.00001 |
| APhasedRepeats | control | Insertion mismatches | Illumina | moderate | 0.00001 | 0.00001 |
| DirectRepeats | motif | Insertion mismatches | Illumina | moderate | 0.00015 | 0.00019 |
| DirectRepeats | control | Insertion mismatches | Illumina | moderate | 0.00001 | 0.00001 |
| G4Motifs | motif | Insertion mismatches | Illumina | moderate | 0.00004 | 0.00005 |
| G4Motifs | control | Insertion mismatches | Illumina | moderate | 0.00001 | 0.00001 |
| InvertedRepeats | motif | Insertion mismatches | Illumina | moderate | 0.00002 | 0.00002 |
| InvertedRepeats | control | Insertion mismatches | Illumina | moderate | 0.00001 | 0.00001 |
| MirrorRepeats | motif | Insertion mismatches | Illumina | moderate | 0.00004 | 0.00010 |
| MirrorRepeats | control | Insertion mismatches | Illumina | moderate | 0.00000 | 0.00001 |
| ZDNAMotifs | motif | Insertion mismatches | Illumina | moderate | 0.00037 | 0.00030 |
| ZDNAMotifs | control | Insertion mismatches | Illumina | moderate | 0.00001 | 0.00001 |
| APhasedRepeats | motif | Deletion mismatches | Illumina | moderate | 0.00002 | 0.00002 |
| APhasedRepeats | control | Deletion mismatches | Illumina | moderate | 0.00002 | 0.00002 |
| DirectRepeats | motif | Deletion mismatches | Illumina | moderate | 0.00014 | 0.00017 |
| DirectRepeats | control | Deletion mismatches | Illumina | moderate | 0.00002 | 0.00002 |
| G4Motifs | motif | Deletion mismatches | Illumina | moderate | 0.00005 | 0.00006 |
| G4Motifs | control | Deletion mismatches | Illumina | moderate | 0.00002 | 0.00002 |
| InvertedRepeats | motif | Deletion mismatches | Illumina | moderate | 0.00003 | 0.00004 |
| InvertedRepeats | control | Deletion mismatches | Illumina | moderate | 0.00002 | 0.00002 |
| MirrorRepeats | motif | Deletion mismatches | Illumina | moderate | 0.00005 | 0.00009 |
| MirrorRepeats | control | Deletion mismatches | Illumina | moderate | 0.00002 | 0.00002 |
| ZDNAMotifs | motif | Deletion mismatches | Illumina | moderate | 0.00031 | 0.00027 |
| ZDNAMotifs | control | Deletion mismatches | Illumina | moderate | 0.00002 | 0.00002 |
| APhasedRepeats | motif | Average read depth per bp | Illumina | moderate | 52.43 | 52.36 |
| APhasedRepeats | control | Average read depth per bp | Illumina | moderate | 49.90 | 49.90 |
| DirectRepeats | motif | Average read depth per bp | Illumina | moderate | 49.93 | 49.94 |
| DirectRepeats | control | Average read depth per bp | Illumina | moderate | 49.47 | 49.60 |
| G4Motifs | motif | Average read depth per bp | Illumina | moderate | 39.09 | 39.54 |
| G4Motifs | control | Average read depth per bp | Illumina | moderate | 48.91 | 48.97 |
| InvertedRepeats | motif | Average read depth per bp | Illumina | moderate | 51.85 | 51.64 |
| InvertedRepeats | control | Average read depth per bp | Illumina | moderate | 49.93 | 49.91 |
| MirrorRepeats | motif | Average read depth per bp | Illumina | moderate | 51.95 | 51.56 |
| MirrorRepeats | control | Average read depth per bp | Illumina | moderate | 49.60 | 49.67 |
| ZDNAMotifs | motif | Average read depth per bp | Illumina | moderate | 47.12 | 47.13 |
| ZDNAMotifs | control | Average read depth per bp | Illumina | moderate | 49.60 | 49.59 |
| APhasedRepeats | motif | Mean basequality | Illumina | moderate | NA* | 36.48 |
| APhasedRepeats | control | Mean basequality | Illumina | moderate | NA | 35.81 |
| DirectRepeats | motif | Mean basequality | Illumina | moderate | NA | 35.54 |
| DirectRepeats | control | Mean basequality | Illumina | moderate | NA | 35.78 |
| G4Motifs | motif | Mean basequality | Illumina | moderate | NA | 34.48 |
| G4Motifs | control | Mean basequality | Illumina | moderate | NA | 35.71 |
| InvertedRepeats | motif | Mean basequality | Illumina | moderate | NA | 36.06 |
| InvertedRepeats | control | Mean basequality | Illumina | moderate | NA | 35.80 |
| MirrorRepeats | motif | Mean basequality | Illumina | moderate | NA | 35.78 |
| MirrorRepeats | control | Mean basequality | Illumina | moderate | NA | 35.81 |
| ZDNAMotifs | motif | Mean basequality | Illumina | moderate | NA | 34.44 |
| ZDNAMotifs | control | Mean basequality | Illumina | moderate | NA | 35.76 |
| APhasedRepeats | motif | Single-nucleotide mismatches | HiFi | moderate | 0.00270 | 0.00134 |
| APhasedRepeats | control | Single-nucleotide mismatches | HiFi | moderate | 0.00142 | 0.00134 |
| DirectRepeats | motif | Single-nucleotide mismatches | HiFi | moderate | 0.00292 | 0.00174 |
| DirectRepeats | control | Single-nucleotide mismatches | HiFi | moderate | 0.00247 | 0.00134 |
| G4Motifs | motif | Single-nucleotide mismatches | HiFi | moderate | 0.00220 | 0.00178 |
| G4Motifs | control | Single-nucleotide mismatches | HiFi | moderate | 0.00141 | 0.00136 |

| | | | | | | |
|---|---|---|---|---|---|---|
| InvertedRepeats | motif | Single-nucleotide mismatches | HiFi | moderate | 0.00159 | 0.00139 |
| InvertedRepeats | control | Single-nucleotide mismatches | HiFi | moderate | 0.00194 | 0.00137 |
| MirrorRepeats | motif | Single-nucleotide mismatches | HiFi | moderate | 0.00148 | 0.00151 |
| MirrorRepeats | control | Single-nucleotide mismatches | HiFi | moderate | 0.00206 | 0.00133 |
| ZDNAMotifs | motif | Single-nucleotide mismatches | HiFi | moderate | 0.00410 | 0.00199 |
| ZDNAMotifs | control | Single-nucleotide mismatches | HiFi | moderate | 0.00158 | 0.00139 |
| APhasedRepeats | motif | Insertion mismatches | HiFi | moderate | 0.00123 | 0.00122 |
| APhasedRepeats | control | Insertion mismatches | HiFi | moderate | 0.00099 | 0.00099 |
| DirectRepeats | motif | Insertion mismatches | HiFi | moderate | 0.00131 | 0.00136 |
| DirectRepeats | control | Insertion mismatches | HiFi | moderate | 0.00099 | 0.00099 |
| G4Motifs | motif | Insertion mismatches | HiFi | moderate | 0.00137 | 0.00139 |
| G4Motifs | control | Insertion mismatches | HiFi | moderate | 0.00099 | 0.00099 |
| InvertedRepeats | motif | Insertion mismatches | HiFi | moderate | 0.00106 | 0.00106 |
| InvertedRepeats | control | Insertion mismatches | HiFi | moderate | 0.00100 | 0.00100 |
| MirrorRepeats | motif | Insertion mismatches | HiFi | moderate | 0.00117 | 0.00130 |
| MirrorRepeats | control | Insertion mismatches | HiFi | moderate | 0.00098 | 0.00099 |
| ZDNAMotifs | motif | Insertion mismatches | HiFi | moderate | 0.00231 | 0.00172 |
| ZDNAMotifs | control | Insertion mismatches | HiFi | moderate | 0.00101 | 0.00101 |
| APhasedRepeats | motif | Deletion mismatches | HiFi | moderate | 0.00139 | 0.00139 |
| APhasedRepeats | control | Deletion mismatches | HiFi | moderate | 0.00104 | 0.00105 |
| DirectRepeats | motif | Deletion mismatches | HiFi | moderate | 0.00166 | 0.00163 |
| DirectRepeats | control | Deletion mismatches | HiFi | moderate | 0.00104 | 0.00105 |
| G4Motifs | motif | Deletion mismatches | HiFi | moderate | 0.00218 | 0.00225 |
| G4Motifs | control | Deletion mismatches | HiFi | moderate | 0.00106 | 0.00106 |
| InvertedRepeats | motif | Deletion mismatches | HiFi | moderate | 0.00135 | 0.00142 |
| InvertedRepeats | control | Deletion mismatches | HiFi | moderate | 0.00107 | 0.00108 |
| MirrorRepeats | motif | Deletion mismatches | HiFi | moderate | 0.00147 | 0.00173 |
| MirrorRepeats | control | Deletion mismatches | HiFi | moderate | 0.00103 | 0.00104 |
| ZDNAMotifs | motif | Deletion mismatches | HiFi | moderate | 0.00071 | 0.00066 |
| ZDNAMotifs | control | Deletion mismatches | HiFi | moderate | 0.00105 | 0.00107 |
| APhasedRepeats | motif | Average read depth per bp | HiFi | moderate | 14.90 | 14.91 |
| APhasedRepeats | control | Average read depth per bp | HiFi | moderate | 14.98 | 14.98 |
| DirectRepeats | motif | Average read depth per bp | HiFi | moderate | 14.90 | 14.91 |
| DirectRepeats | control | Average read depth per bp | HiFi | moderate | 15.06 | 15.07 |
| G4Motifs | motif | Average read depth per bp | HiFi | moderate | 14.98 | 15.03 |
| G4Motifs | control | Average read depth per bp | HiFi | moderate | 15.05 | 15.04 |
| InvertedRepeats | motif | Average read depth per bp | HiFi | moderate | 14.87 | 14.89 |
| InvertedRepeats | control | Average read depth per bp | HiFi | moderate | 15.02 | 15.02 |
| MirrorRepeats | motif | Average read depth per bp | HiFi | moderate | 14.72 | 14.73 |
| MirrorRepeats | control | Average read depth per bp | HiFi | moderate | 15.04 | 15.05 |
| ZDNAMotifs | motif | Average read depth per bp | HiFi | moderate | 15.10 | 15.12 |
| ZDNAMotifs | control | Average read depth per bp | HiFi | moderate | 15.00 | 15.00 |
| APhasedRepeats | motif | Mean basequality | HiFi | moderate | NA | 84.39 |
| APhasedRepeats | control | Mean basequality | HiFi | moderate | NA | 78.09 |
| DirectRepeats | motif | Mean basequality | HiFi | moderate | NA | 77.49 |
| DirectRepeats | control | Mean basequality | HiFi | moderate | NA | 78.04 |
| G4Motifs | motif | Mean basequality | HiFi | moderate | NA | 84.38 |
| G4Motifs | control | Mean basequality | HiFi | moderate | NA | 77.97 |
| InvertedRepeats | motif | Mean basequality | HiFi | moderate | NA | 78.20 |
| InvertedRepeats | control | Mean basequality | HiFi | moderate | NA | 78.05 |
| MirrorRepeats | motif | Mean basequality | HiFi | moderate | NA | 78.33 |
| MirrorRepeats | control | Mean basequality | HiFi | moderate | NA | 78.04 |
| ZDNAMotifs | motif | Mean basequality | HiFi | moderate | NA | 69.99 |
| ZDNAMotifs | control | Mean basequality | HiFi | moderate | NA | 78.02 |
| APhasedRepeats | motif | Single-nucleotide mismatches | ONT | moderate | 0.01835 | 0.01518 |
| APhasedRepeats | control | Single-nucleotide mismatches | ONT | moderate | 0.01704 | 0.01655 |
| DirectRepeats | motif | Single-nucleotide mismatches | ONT | moderate | 0.02561 | 0.01476 |
| DirectRepeats | control | Single-nucleotide mismatches | ONT | moderate | 0.01850 | 0.01666 |
| G4Motifs | motif | Single-nucleotide mismatches | ONT | moderate | 0.02197 | 0.02123 |
| G4Motifs | control | Single-nucleotide mismatches | ONT | moderate | 0.01721 | 0.01692 |
| InvertedRepeats | motif | Single-nucleotide mismatches | ONT | moderate | 0.01788 | 0.01567 |
| InvertedRepeats | control | Single-nucleotide mismatches | ONT | moderate | 0.01772 | 0.01660 |
| MirrorRepeats | motif | Single-nucleotide mismatches | ONT | moderate | 0.01517 | 0.01417 |
| MirrorRepeats | control | Single-nucleotide mismatches | ONT | moderate | 0.01743 | 0.01657 |
| ZDNAMotifs | motif | Single-nucleotide mismatches | ONT | moderate | 0.01617 | 0.01500 |
| ZDNAMotifs | control | Single-nucleotide mismatches | ONT | moderate | 0.01701 | 0.01671 |
| APhasedRepeats | motif | Insertion mismatches | ONT | moderate | 0.00668 | 0.00668 |
| APhasedRepeats | control | Insertion mismatches | ONT | moderate | 0.00664 | 0.00664 |
| DirectRepeats | motif | Insertion mismatches | ONT | moderate | 0.00728 | 0.00693 |

| | | | | | | |
|---|---|---|---|---|---|---|
| DirectRepeats | control | Insertion mismatches | ONT | moderate | 0.00667 | 0.00666 |
| G4Motifs | motif | Insertion mismatches | ONT | moderate | 0.01075 | 0.01085 |
| G4Motifs | control | Insertion mismatches | ONT | moderate | 0.00679 | 0.00677 |
| InvertedRepeats | motif | Insertion mismatches | ONT | moderate | 0.00660 | 0.00651 |
| InvertedRepeats | control | Insertion mismatches | ONT | moderate | 0.00664 | 0.00663 |
| MirrorRepeats | motif | Insertion mismatches | ONT | moderate | 0.00685 | 0.00698 |
| MirrorRepeats | control | Insertion mismatches | ONT | moderate | 0.00667 | 0.00667 |
| ZDNAMotifs | motif | Insertion mismatches | ONT | moderate | 0.00757 | 0.00722 |
| ZDNAMotifs | control | Insertion mismatches | ONT | moderate | 0.00666 | 0.00665 |
| APhasedRepeats | motif | Deletion mismatches | ONT | moderate | 0.01062 | 0.01061 |
| APhasedRepeats | control | Deletion mismatches | ONT | moderate | 0.01033 | 0.01033 |
| DirectRepeats | motif | Deletion mismatches | ONT | moderate | 0.01019 | 0.01002 |
| DirectRepeats | control | Deletion mismatches | ONT | moderate | 0.01032 | 0.01036 |
| G4Motifs | motif | Deletion mismatches | ONT | moderate | 0.01708 | 0.01735 |
| G4Motifs | control | Deletion mismatches | ONT | moderate | 0.01056 | 0.01055 |
| InvertedRepeats | motif | Deletion mismatches | ONT | moderate | 0.01043 | 0.01042 |
| InvertedRepeats | control | Deletion mismatches | ONT | moderate | 0.01040 | 0.01048 |
| MirrorRepeats | motif | Deletion mismatches | ONT | moderate | 0.01051 | 0.01060 |
| MirrorRepeats | control | Deletion mismatches | ONT | moderate | 0.01026 | 0.01029 |
| ZDNAMotifs | motif | Deletion mismatches | ONT | moderate | 0.01134 | 0.01175 |
| ZDNAMotifs | control | Deletion mismatches | ONT | moderate | 0.01053 | 0.01060 |
| APhasedRepeats | motif | Average read depth per bp | ONT | moderate | 28.72 | 28.73 |
| APhasedRepeats | control | Average read depth per bp | ONT | moderate | 28.54 | 28.54 |
| DirectRepeats | motif | Average read depth per bp | ONT | moderate | 29.03 | 29.07 |
| DirectRepeats | control | Average read depth per bp | ONT | moderate | 28.56 | 28.62 |
| G4Motifs | motif | Average read depth per bp | ONT | moderate | 28.40 | 28.41 |
| G4Motifs | control | Average read depth per bp | ONT | moderate | 28.49 | 28.49 |
| InvertedRepeats | motif | Average read depth per bp | ONT | moderate | 28.56 | 28.59 |
| InvertedRepeats | control | Average read depth per bp | ONT | moderate | 28.55 | 28.57 |
| MirrorRepeats | motif | Average read depth per bp | ONT | moderate | 28.45 | 28.48 |
| MirrorRepeats | control | Average read depth per bp | ONT | moderate | 28.53 | 28.56 |
| ZDNAMotifs | motif | Average read depth per bp | ONT | moderate | 28.48 | 28.49 |
| ZDNAMotifs | control | Average read depth per bp | ONT | moderate | 28.50 | 28.51 |
| APhasedRepeats | motif | Single-nucleotide mismatches | Illumina | stringent | 0.00200 | 0.00205 |
| APhasedRepeats | control | Single-nucleotide mismatches | Illumina | stringent | 0.00222 | 0.00229 |
| DirectRepeats | motif | Single-nucleotide mismatches | Illumina | stringent | 0.00237 | 0.00256 |
| DirectRepeats | control | Single-nucleotide mismatches | Illumina | stringent | 0.00219 | 0.00226 |
| G4Motifs | motif | Single-nucleotide mismatches | Illumina | stringent | 0.00440 | 0.00472 |
| G4Motifs | control | Single-nucleotide mismatches | Illumina | stringent | 0.00225 | 0.00236 |
| InvertedRepeats | motif | Single-nucleotide mismatches | Illumina | stringent | 0.00207 | 0.00215 |
| InvertedRepeats | control | Single-nucleotide mismatches | Illumina | stringent | 0.00224 | 0.00234 |
| MirrorRepeats | motif | Single-nucleotide mismatches | Illumina | stringent | 0.00213 | 0.00233 |
| MirrorRepeats | control | Single-nucleotide mismatches | Illumina | stringent | 0.00216 | 0.00227 |
| ZDNAMotifs | motif | Single-nucleotide mismatches | Illumina | stringent | 0.00343 | 0.00361 |
| ZDNAMotifs | control | Single-nucleotide mismatches | Illumina | stringent | 0.00229 | 0.00242 |
| APhasedRepeats | motif | Insertion mismatches | Illumina | stringent | 0.00002 | 0.00002 |
| APhasedRepeats | control | Insertion mismatches | Illumina | stringent | 0.00000 | 0.00000 |
| DirectRepeats | motif | Insertion mismatches | Illumina | stringent | 0.00002 | 0.00003 |
| DirectRepeats | control | Insertion mismatches | Illumina | stringent | 0.00001 | 0.00001 |
| G4Motifs | motif | Insertion mismatches | Illumina | stringent | 0.00002 | 0.00003 |
| G4Motifs | control | Insertion mismatches | Illumina | stringent | 0.00000 | 0.00000 |
| InvertedRepeats | motif | Insertion mismatches | Illumina | stringent | 0.00001 | 0.00001 |
| InvertedRepeats | control | Insertion mismatches | Illumina | stringent | 0.00001 | 0.00001 |
| MirrorRepeats | motif | Insertion mismatches | Illumina | stringent | 0.00001 | 0.00001 |
| MirrorRepeats | control | Insertion mismatches | Illumina | stringent | 0.00001 | 0.00001 |
| ZDNAMotifs | motif | Insertion mismatches | Illumina | stringent | 0.00000 | 0.00000 |
| ZDNAMotifs | control | Insertion mismatches | Illumina | stringent | 0.00000 | 0.00000 |
| APhasedRepeats | motif | Deletion mismatches | Illumina | stringent | 0.00002 | 0.00002 |
| APhasedRepeats | control | Deletion mismatches | Illumina | stringent | 0.00002 | 0.00002 |
| DirectRepeats | motif | Deletion mismatches | Illumina | stringent | 0.00007 | 0.00007 |
| DirectRepeats | control | Deletion mismatches | Illumina | stringent | 0.00002 | 0.00002 |
| G4Motifs | motif | Deletion mismatches | Illumina | stringent | 0.00005 | 0.00005 |
| G4Motifs | control | Deletion mismatches | Illumina | stringent | 0.00002 | 0.00002 |
| InvertedRepeats | motif | Deletion mismatches | Illumina | stringent | 0.00002 | 0.00002 |
| InvertedRepeats | control | Deletion mismatches | Illumina | stringent | 0.00002 | 0.00002 |
| MirrorRepeats | motif | Deletion mismatches | Illumina | stringent | 0.00002 | 0.00002 |
| MirrorRepeats | control | Deletion mismatches | Illumina | stringent | 0.00002 | 0.00002 |
| ZDNAMotifs | motif | Deletion mismatches | Illumina | stringent | 0.00001 | 0.00001 |
| ZDNAMotifs | control | Deletion mismatches | Illumina | stringent | 0.00002 | 0.00002 |

| | | | | | | |
|---|---|---|---|---|---|---|
| APhasedRepeats | motif | Average read depth per bp | Illumina | stringent | 51.62 | 51.56 |
| APhasedRepeats | control | Average read depth per bp | Illumina | stringent | 49.47 | 49.47 |
| DirectRepeats | motif | Average read depth per bp | Illumina | stringent | 49.40 | 49.46 |
| DirectRepeats | control | Average read depth per bp | Illumina | stringent | 49.16 | 49.17 |
| G4Motifs | motif | Average read depth per bp | Illumina | stringent | 40.03 | 40.17 |
| G4Motifs | control | Average read depth per bp | Illumina | stringent | 48.70 | 48.70 |
| InvertedRepeats | motif | Average read depth per bp | Illumina | stringent | 50.66 | 50.55 |
| InvertedRepeats | control | Average read depth per bp | Illumina | stringent | 49.25 | 49.25 |
| MirrorRepeats | motif | Average read depth per bp | Illumina | stringent | 51.32 | 51.27 |
| MirrorRepeats | control | Average read depth per bp | Illumina | stringent | 49.26 | 49.23 |
| ZDNAMotifs | motif | Average read depth per bp | Illumina | stringent | 44.10 | 44.12 |
| ZDNAMotifs | control | Average read depth per bp | Illumina | stringent | 49.05 | 49.02 |
| APhasedRepeats | motif | Mean basequality | Illumina | stringent | NA | 36.49 |
| APhasedRepeats | control | Mean basequality | Illumina | stringent | NA | 35.88 |
| DirectRepeats | motif | Mean basequality | Illumina | stringent | NA | 35.79 |
| DirectRepeats | control | Mean basequality | Illumina | stringent | NA | 35.86 |
| G4Motifs | motif | Mean basequality | Illumina | stringent | NA | 34.61 |
| G4Motifs | control | Mean basequality | Illumina | stringent | NA | 35.79 |
| InvertedRepeats | motif | Mean basequality | Illumina | stringent | NA | 36.11 |
| InvertedRepeats | control | Mean basequality | Illumina | stringent | NA | 35.88 |
| MirrorRepeats | motif | Mean basequality | Illumina | stringent | NA | 35.96 |
| MirrorRepeats | control | Mean basequality | Illumina | stringent | NA | 35.89 |
| ZDNAMotifs | motif | Mean basequality | Illumina | stringent | NA | 34.74 |
| ZDNAMotifs | control | Mean basequality | Illumina | stringent | NA | 35.85 |
| APhasedRepeats | motif | Single-nucleotide mismatches | HiFi | stringent | 0.00133 | 0.00132 |
| APhasedRepeats | control | Single-nucleotide mismatches | HiFi | stringent | 0.00143 | 0.00127 |
| DirectRepeats | motif | Single-nucleotide mismatches | HiFi | stringent | 0.00140 | 0.00138 |
| DirectRepeats | control | Single-nucleotide mismatches | HiFi | stringent | 0.00130 | 0.00130 |
| G4Motifs | motif | Single-nucleotide mismatches | HiFi | stringent | 0.00176 | 0.00168 |
| G4Motifs | control | Single-nucleotide mismatches | HiFi | stringent | 0.00133 | 0.00139 |
| InvertedRepeats | motif | Single-nucleotide mismatches | HiFi | stringent | 0.00141 | 0.00136 |
| InvertedRepeats | control | Single-nucleotide mismatches | HiFi | stringent | 0.00150 | 0.00134 |
| MirrorRepeats | motif | Single-nucleotide mismatches | HiFi | stringent | 0.00142 | 0.00150 |
| MirrorRepeats | control | Single-nucleotide mismatches | HiFi | stringent | 0.00133 | 0.00137 |
| ZDNAMotifs | motif | Single-nucleotide mismatches | HiFi | stringent | 0.00169 | 0.00163 |
| ZDNAMotifs | control | Single-nucleotide mismatches | HiFi | stringent | 0.00141 | 0.00144 |
| APhasedRepeats | motif | Insertion mismatches | HiFi | stringent | 0.00122 | 0.00121 |
| APhasedRepeats | control | Insertion mismatches | HiFi | stringent | 0.00100 | 0.00100 |
| DirectRepeats | motif | Insertion mismatches | HiFi | stringent | 0.00105 | 0.00106 |
| DirectRepeats | control | Insertion mismatches | HiFi | stringent | 0.00102 | 0.00102 |
| G4Motifs | motif | Insertion mismatches | HiFi | stringent | 0.00127 | 0.00130 |
| G4Motifs | control | Insertion mismatches | HiFi | stringent | 0.00101 | 0.00101 |
| InvertedRepeats | motif | Insertion mismatches | HiFi | stringent | 0.00108 | 0.00108 |
| InvertedRepeats | control | Insertion mismatches | HiFi | stringent | 0.00100 | 0.00100 |
| MirrorRepeats | motif | Insertion mismatches | HiFi | stringent | 0.00113 | 0.00121 |
| MirrorRepeats | control | Insertion mismatches | HiFi | stringent | 0.00101 | 0.00101 |
| ZDNAMotifs | motif | Insertion mismatches | HiFi | stringent | 0.00137 | 0.00139 |
| ZDNAMotifs | control | Insertion mismatches | HiFi | stringent | 0.00101 | 0.00101 |
| APhasedRepeats | motif | Deletion mismatches | HiFi | stringent | 0.00138 | 0.00137 |
| APhasedRepeats | control | Deletion mismatches | HiFi | stringent | 0.00110 | 0.00110 |
| DirectRepeats | motif | Deletion mismatches | HiFi | stringent | 0.00130 | 0.00130 |
| DirectRepeats | control | Deletion mismatches | HiFi | stringent | 0.00110 | 0.00111 |
| G4Motifs | motif | Deletion mismatches | HiFi | stringent | 0.00237 | 0.00243 |
| G4Motifs | control | Deletion mismatches | HiFi | stringent | 0.00112 | 0.00112 |
| InvertedRepeats | motif | Deletion mismatches | HiFi | stringent | 0.00149 | 0.00153 |
| InvertedRepeats | control | Deletion mismatches | HiFi | stringent | 0.00111 | 0.00111 |
| MirrorRepeats | motif | Deletion mismatches | HiFi | stringent | 0.00156 | 0.00175 |
| MirrorRepeats | control | Deletion mismatches | HiFi | stringent | 0.00110 | 0.00111 |
| ZDNAMotifs | motif | Deletion mismatches | HiFi | stringent | 0.00027 | 0.00027 |
| ZDNAMotifs | control | Deletion mismatches | HiFi | stringent | 0.00111 | 0.00112 |
| APhasedRepeats | motif | Average read depth per bp | HiFi | stringent | 14.95 | 14.96 |
| APhasedRepeats | control | Average read depth per bp | HiFi | stringent | 15.05 | 15.05 |
| DirectRepeats | motif | Average read depth per bp | HiFi | stringent | 14.98 | 14.99 |
| DirectRepeats | control | Average read depth per bp | HiFi | stringent | 15.06 | 15.06 |
| G4Motifs | motif | Average read depth per bp | HiFi | stringent | 15.23 | 15.25 |
| G4Motifs | control | Average read depth per bp | HiFi | stringent | 15.12 | 15.11 |
| InvertedRepeats | motif | Average read depth per bp | HiFi | stringent | 15.00 | 15.01 |
| InvertedRepeats | control | Average read depth per bp | HiFi | stringent | 15.07 | 15.07 |
| MirrorRepeats | motif | Average read depth per bp | HiFi | stringent | 14.89 | 14.89 |

| | | | | | | |
|---|---|---|---|---|---|---|
| MirrorRepeats | control | Average read depth per bp | HiFi | stringent | 15.08 | 15.08 |
| ZDNAMotifs | motif | Average read depth per bp | HiFi | stringent | 15.07 | 15.08 |
| ZDNAMotifs | control | Average read depth per bp | HiFi | stringent | 15.07 | 15.08 |
| APhasedRepeats | motif | Mean basequality | HiFi | stringent | NA | 84.83 |
| APhasedRepeats | control | Mean basequality | HiFi | stringent | NA | 82.74 |
| DirectRepeats | motif | Mean basequality | HiFi | stringent | NA | 82.73 |
| DirectRepeats | control | Mean basequality | HiFi | stringent | NA | 82.74 |
| G4Motifs | motif | Mean basequality | HiFi | stringent | NA | 84.72 |
| G4Motifs | control | Mean basequality | HiFi | stringent | NA | 82.61 |
| InvertedRepeats | motif | Mean basequality | HiFi | stringent | NA | 82.97 |
| InvertedRepeats | control | Mean basequality | HiFi | stringent | NA | 82.71 |
| MirrorRepeats | motif | Mean basequality | HiFi | stringent | NA | 83.16 |
| MirrorRepeats | control | Mean basequality | HiFi | stringent | NA | 82.70 |
| ZDNAMotifs | motif | Mean basequality | HiFi | stringent | NA | 79.24 |
| ZDNAMotifs | control | Mean basequality | HiFi | stringent | NA | 82.66 |
| APhasedRepeats | motif | Single-nucleotide mismatches | ONT | stringent | 0.01542 | 0.01533 |
| APhasedRepeats | control | Single-nucleotide mismatches | ONT | stringent | 0.01715 | 0.01702 |
| DirectRepeats | motif | Single-nucleotide mismatches | ONT | stringent | 0.01671 | 0.01641 |
| DirectRepeats | control | Single-nucleotide mismatches | ONT | stringent | 0.01728 | 0.01712 |
| G4Motifs | motif | Single-nucleotide mismatches | ONT | stringent | 0.02248 | 0.02211 |
| G4Motifs | control | Single-nucleotide mismatches | ONT | stringent | 0.01746 | 0.01738 |
| InvertedRepeats | motif | Single-nucleotide mismatches | ONT | stringent | 0.01663 | 0.01632 |
| InvertedRepeats | control | Single-nucleotide mismatches | ONT | stringent | 0.01734 | 0.01712 |
| MirrorRepeats | motif | Single-nucleotide mismatches | ONT | stringent | 0.01602 | 0.01557 |
| MirrorRepeats | control | Single-nucleotide mismatches | ONT | stringent | 0.01756 | 0.01717 |
| ZDNAMotifs | motif | Single-nucleotide mismatches | ONT | stringent | 0.02015 | 0.01992 |
| ZDNAMotifs | control | Single-nucleotide mismatches | ONT | stringent | 0.01738 | 0.01728 |
| APhasedRepeats | motif | Insertion mismatches | ONT | stringent | 0.00673 | 0.00672 |
| APhasedRepeats | control | Insertion mismatches | ONT | stringent | 0.00682 | 0.00682 |
| DirectRepeats | motif | Insertion mismatches | ONT | stringent | 0.00673 | 0.00676 |
| DirectRepeats | control | Insertion mismatches | ONT | stringent | 0.00683 | 0.00683 |
| G4Motifs | motif | Insertion mismatches | ONT | stringent | 0.01035 | 0.01047 |
| G4Motifs | control | Insertion mismatches | ONT | stringent | 0.00690 | 0.00689 |
| InvertedRepeats | motif | Insertion mismatches | ONT | stringent | 0.00687 | 0.00686 |
| InvertedRepeats | control | Insertion mismatches | ONT | stringent | 0.00681 | 0.00681 |
| MirrorRepeats | motif | Insertion mismatches | ONT | stringent | 0.00693 | 0.00703 |
| MirrorRepeats | control | Insertion mismatches | ONT | stringent | 0.00682 | 0.00683 |
| ZDNAMotifs | motif | Insertion mismatches | ONT | stringent | 0.00668 | 0.00669 |
| ZDNAMotifs | control | Insertion mismatches | ONT | stringent | 0.00682 | 0.00684 |
| APhasedRepeats | motif | Deletion mismatches | ONT | stringent | 0.01074 | 0.01071 |
| APhasedRepeats | control | Deletion mismatches | ONT | stringent | 0.01062 | 0.01062 |
| DirectRepeats | motif | Deletion mismatches | ONT | stringent | 0.01063 | 0.01064 |
| DirectRepeats | control | Deletion mismatches | ONT | stringent | 0.01063 | 0.01067 |
| G4Motifs | motif | Deletion mismatches | ONT | stringent | 0.01615 | 0.01641 |
| G4Motifs | control | Deletion mismatches | ONT | stringent | 0.01073 | 0.01073 |
| InvertedRepeats | motif | Deletion mismatches | ONT | stringent | 0.01130 | 0.01129 |
| InvertedRepeats | control | Deletion mismatches | ONT | stringent | 0.01075 | 0.01079 |
| MirrorRepeats | motif | Deletion mismatches | ONT | stringent | 0.01129 | 0.01168 |
| MirrorRepeats | control | Deletion mismatches | ONT | stringent | 0.01055 | 0.01058 |
| ZDNAMotifs | motif | Deletion mismatches | ONT | stringent | 0.01282 | 0.01287 |
| ZDNAMotifs | control | Deletion mismatches | ONT | stringent | 0.01078 | 0.01086 |
| APhasedRepeats | motif | Average read depth per bp | ONT | stringent | 28.50 | 28.51 |
| APhasedRepeats | control | Average read depth per bp | ONT | stringent | 28.45 | 28.45 |
| DirectRepeats | motif | Average read depth per bp | ONT | stringent | 28.51 | 28.51 |
| DirectRepeats | control | Average read depth per bp | ONT | stringent | 28.58 | 28.58 |
| G4Motifs | motif | Average read depth per bp | ONT | stringent | 28.41 | 28.42 |
| G4Motifs | control | Average read depth per bp | ONT | stringent | 28.49 | 28.49 |
| InvertedRepeats | motif | Average read depth per bp | ONT | stringent | 28.51 | 28.52 |
| InvertedRepeats | control | Average read depth per bp | ONT | stringent | 28.52 | 28.52 |
| MirrorRepeats | motif | Average read depth per bp | ONT | stringent | 28.49 | 28.49 |
| MirrorRepeats | control | Average read depth per bp | ONT | stringent | 28.56 | 28.55 |
| ZDNAMotifs | motif | Average read depth per bp | ONT | stringent | 28.43 | 28.43 |
| ZDNAMotifs | control | Average read depth per bp | ONT | stringent | 28.52 | 28.51 |
| Overall | Overall | Deletion mismatches | HiFi | moderate | 0.00124 | 0.00127 |
| Overall | Overall | Insertion mismatches | HiFi | moderate | 0.00107 | 0.00108 |
| Overall | Overall | Single-nucleotide mismatches | HiFi | moderate | 0.00188 | 0.00141 |
| Overall | Overall | Deletion mismatches | Illumina | moderate | 0.00003 | 0.00004 |
| Overall | Overall | Insertion mismatches | Illumina | moderate | 0.00002 | 0.00003 |
| Overall | Overall | Single-nucleotide mismatches | Illumina | moderate | 0.00224 | 0.00240 |

| | | | | | | |
|---|---|---|---|---|---|---|
| Overall | Overall | Deletion mismatches | ONT | moderate | 0.01062 | 0.01062 |
| Overall | Overall | Insertion mismatches | ONT | moderate | 0.00682 | 0.00673 |
| Overall | Overall | Single-nucleotide mismatches | ONT | moderate | 0.01789 | 0.01614 |
| Overall | Overall | Deletion mismatches | HiFi | stringent | 0.00133 | 0.00133 |
| Overall | Overall | Insertion mismatches | HiFi | stringent | 0.00106 | 0.00105 |
| Overall | Overall | Single-nucleotide mismatches | HiFi | stringent | 0.00145 | 0.00136 |
| Overall | Overall | Deletion mismatches | Illumina | stringent | 0.00002 | 0.00002 |
| Overall | Overall | Insertion mismatches | Illumina | stringent | 0.00001 | 0.00001 |
| Overall | Overall | Single-nucleotide mismatches | Illumina | stringent | 0.00222 | 0.00231 |
| Overall | Overall | Deletion mismatches | ONT | stringent | 0.01112 | 0.01110 |
| Overall | Overall | Insertion mismatches | ONT | stringent | 0.00695 | 0.00690 |
| Overall | Overall | Single-nucleotide mismatches | ONT | stringent | 0.01714 | 0.01683 |

NA* - not applicable or not available

**Table S2B. Overall mismatch error rates.** Shown are mismatch (single-nucleotide, insertion, and deletion) error rates across all motifs and controls of the moderately filtered set.

| Technology | Single-nucleotide | Insertion | Deletion |
|---|---|---|---|
| Illumina | 0.002241 | 0.000024 | 0.000034 |
| HiFi | 0.001881 | 0.001072 | 0.001242 |
| ONT | 0.017890 | 0.006818 | 0.010617 |

**Table S3. Adjusted p-values calculated for t-test comparing error rates between motifs and controls and adjusted for multiple testing with the Benjamini-Hochberg correction.** P-values below 0.05 are shown in bold.

| Type | Tech | Filter level | adjusted P-value SNM | adjusted P-value Insertion | adjusted P-value Deletion |
|---|---|---|---|---|---|
| APhasedRepeats | Illumina | moderate | **4.75E-94** | 0.056 | **0.006** |
| DirectRepeats | Illumina | moderate | **0** | **0** | **0** |
| G4Motifs | Illumina | moderate | **0** | **7.21E-92** | **1.24E-82** |
| InvertedRepeats | Illumina | moderate | **0** | **2.47E-163** | **4.00E-117** |
| MirrorRepeats | Illumina | moderate | **0** | **0** | **0.00E+00** |
| ZDNAMotifs | Illumina | moderate | **0** | **0** | **1.01E-293** |
| APhasedRepeats | HiFi | moderate | 0.820 | **0** | **0** |
| DirectRepeats | HiFi | moderate | **4.35E-34** | **0** | **0** |
| G4Motifs | HiFi | moderate | **2.56E-25** | **0** | **0** |
| InvertedRepeats | HiFi | moderate | **0.0237** | **1.77E-293** | **0** |
| MirrorRepeats | HiFi | moderate | **2.07E-11** | **0** | **0** |
| ZDNAMotifs | HiFi | moderate | **5.39E-38** | **0** | **1.65E-282** |
| APhasedRepeats | ONT | moderate | **9.54E-124** | **0.000** | **1.19E-43** |
| DirectRepeats | ONT | moderate | **6.91E-270** | **7.59E-133** | **5.13E-73** |
| G4Motifs | ONT | moderate | **0** | **0** | **0** |
| InvertedRepeats | ONT | moderate | **0** | **9.61E-216** | **1.34E-18** |
| MirrorRepeats | ONT | moderate | **0** | **0** | **5.01E-122** |
| ZDNAMotifs | ONT | moderate | **6.74E-100** | **1.04E-209** | **0** |
| APhasedRepeats | Illumina | stringent | **1.73E-14** | **0.008** | **0.241** |
| DirectRepeats | Illumina | stringent | **3.55E-11** | **0.000** | **2.30E-12** |
| G4Motifs | Illumina | stringent | **2.41E-289** | **1.78E-05** | **1.36E-08** |
| InvertedRepeats | Illumina | stringent | **2.42E-93** | 0.068 | **9.35E-05** |
| MirrorRepeats | Illumina | stringent | 0.820 | 0.391 | 0.241 |
| ZDNAMotifs | Illumina | stringent | **6.84E-21** | 0.182 | 0.060 |
| APhasedRepeats | HiFi | stringent | 0.820 | **2.82E-43** | **1.00E-55** |
| DirectRepeats | HiFi | stringent | 0.820 | **0.18246421** | **1.91E-16** |
| G4Motifs | HiFi | stringent | 0.360 | **1.10E-49** | **0** |
| InvertedRepeats | HiFi | stringent | 0.820 | **5.31E-54** | **0** |
| MirrorRepeats | HiFi | stringent | 0.820 | **4.25E-26** | **1.46E-150** |
| ZDNAMotifs | HiFi | stringent | 0.820 | **9.73E-09** | **2.57E-57** |
| APhasedRepeats | ONT | stringent | **1.04E-31** | **0.006** | 0.241 |
| DirectRepeats | ONT | stringent | **0.000** | 0.103 | 0.619 |
| G4Motifs | ONT | stringent | **2.38E-101** | **0** | **0** |
| InvertedRepeats | ONT | stringent | **1.73E-93** | **5.59E-08** | **2.36E-179** |
| MirrorRepeats | ONT | stringent | **4.26E-16** | **4.27E-09** | **1.61E-63** |
| ZDNAMotifs | ONT | stringent | **6.57E-12** | **0.18246421** | **2.19E-47** |

**Table S4. Poisson regressions for different reduced models.** Shown are % deviance explained for the full model, and the % reduction in deviance explained when individual predictors are removed for different non-B motif types, error types, technologies, and filtering schemes.

| | | | | | Reduction in deviance explained when removing the following predictors | | | |
|---|---|---|---|---|---|---|---|---|
| **Non-B type** | **Error type** | **Technology** | **Filtering level** | **Deviance explained (full model)** | **Motif vs. control** | **Nucleotide composition** | **Motif length** | **Presence of homopolymers** |
| APhasedRepeats | SNM | Illumina | moderate | 1.70% | 0.00% | 89.37% | 0.06% | 0.01% |
| DirectRepeats | SNM | Illumina | moderate | 5.32% | 11.90% | 90.81% | 3.09% | 1.75% |
| G4Motifs | SNM | Illumina | moderate | 16.69% | 5.29% | 27.77% | 8.99% | 0.26% |
| InvertedRepeats | SNM | Illumina | moderate | 2.45% | 1.84% | 92.15% | 1.86% | 0.30% |
| MirrorRepeats | SNM | Illumina | moderate | 8.70% | 5.84% | 97.80% | 1.15% | 0.87% |
| ZDNAMotifs | SNM | Illumina | moderate | 4.61% | 31.52% | 37.56% | 0.19% | 0.59% |
| APhasedRepeats | Insertion | Illumina | moderate | 1.01% | 21.64% | 25.88% | 6.38% | 28.89% |
| DirectRepeats | Insertion | Illumina | moderate | 16.96% | 47.71% | 9.83% | 13.25% | 2.95% |
| G4Motifs | Insertion | Illumina | moderate | 9.03% | 1.51% | 32.38% | 2.48% | 1.08% |
| InvertedRepeats | Insertion | Illumina | moderate | 2.39% | 85.23% | 2.68% | 2.01% | 4.64% |
| MirrorRepeats | Insertion | Illumina | moderate | 20.46% | 39.48% | 6.60% | 38.15% | 5.30% |
| ZDNAMotifs | Insertion | Illumina | moderate | 21.54% | 83.83% | 4.23% | 31.07% | 0.00% |
| APhasedRepeats | Deletion | Illumina | moderate | 0.35% | 28.10% | 15.53% | 11.25% | 39.10% |
| DirectRepeats | Deletion | Illumina | moderate | 9.38% | 64.53% | 11.11% | 1.50% | 0.95% |
| G4Motifs | Deletion | Illumina | moderate | 3.10% | 1.98% | 11.63% | 2.90% | 1.68% |
| InvertedRepeats | Deletion | Illumina | moderate | 0.75% | 75.25% | 3.09% | 20.93% | 2.60% |
| MirrorRepeats | Deletion | Illumina | moderate | 9.71% | 47.26% | 4.61% | 49.44% | 2.71% |
| ZDNAMotifs | Deletion | Illumina | moderate | 16.34% | 83.20% | 3.96% | 23.72% | 0.08% |
| APhasedRepeats | SNM | HiFi | moderate | 0.08% | 37.94% | 66.03% | 13.56% | 1.09% |
| DirectRepeats | SNM | HiFi | moderate | 0.69% | 33.24% | 23.30% | 22.34% | 3.48% |
| G4Motifs | SNM | HiFi | moderate | 0.32% | 5.60% | 22.20% | 12.41% | 2.81% |
| InvertedRepeats | SNM | HiFi | moderate | 0.06% | 13.33% | 47.78% | 49.96% | 14.10% |
| MirrorRepeats | SNM | HiFi | moderate | 0.18% | 13.28% | 19.75% | 64.25% | 0.15% |
| ZDNAMotifs | SNM | HiFi | moderate | 0.66% | 53.26% | 4.37% | 3.55% | 0.28% |
| APhasedRepeats | Insertion | HiFi | moderate | 1.87% | 11.59% | 1.44% | 0.32% | 73.45% |
| DirectRepeats | Insertion | HiFi | moderate | 2.32% | 42.60% | 14.95% | 15.77% | 39.21% |
| G4Motifs | Insertion | HiFi | moderate | 2.36% | 1.42% | 22.45% | 1.40% | 19.71% |
| InvertedRepeats | Insertion | HiFi | moderate | 1.21% | 3.12% | 0.75% | 22.59% | 95.93% |
| MirrorRepeats | Insertion | HiFi | moderate | 2.07% | 44.92% | 15.64% | 50.95% | 26.84% |
| ZDNAMotifs | Insertion | HiFi | moderate | 4.61% | 93.49% | 0.55% | 33.89% | 6.00% |
| APhasedRepeats | Deletion | HiFi | moderate | 7.66% | 2.89% | 0.46% | 0.45% | 87.24% |
| DirectRepeats | Deletion | HiFi | moderate | 11.95% | 17.47% | 0.51% | 9.13% | 79.36% |
| G4Motifs | Deletion | HiFi | moderate | 11.86% | 5.11% | 2.31% | 4.88% | 30.93% |
| InvertedRepeats | Deletion | HiFi | moderate | 11.19% | 4.29% | 0.01% | 24.98% | 91.71% |
| MirrorRepeats | Deletion | HiFi | moderate | 11.67% | 19.91% | 0.86% | 39.06% | 57.85% |
| ZDNAMotifs | Deletion | HiFi | moderate | 4.78% | 18.57% | 0.94% | 20.04% | 66.09% |
| APhasedRepeats | SNM | ONT | moderate | 1.22% | 2.11% | 69.12% | 0.66% | 7.70% |
| DirectRepeats | SNM | ONT | moderate | 1.54% | 12.38% | 76.79% | 10.65% | 4.97% |
| G4Motifs | SNM | ONT | moderate | 2.94% | 1.06% | 39.33% | 0.14% | 0.07% |
| InvertedRepeats | SNM | ONT | moderate | 1.47% | 1.30% | 84.84% | 2.18% | 3.17% |
| MirrorRepeats | SNM | ONT | moderate | 2.30% | 15.83% | 68.10% | 14.18% | 0.29% |
| ZDNAMotifs | SNM | ONT | moderate | 1.54% | 70.59% | 37.91% | 55.07% | 0.35% |
| APhasedRepeats | Insertion | ONT | moderate | 2.77% | 7.21% | 88.27% | 0.01% | 21.94% |
| DirectRepeats | Insertion | ONT | moderate | 5.00% | 3.53% | 87.09% | 1.69% | 16.70% |
| G4Motifs | Insertion | ONT | moderate | 22.50% | 2.46% | 14.34% | 0.18% | 0.19% |
| InvertedRepeats | Insertion | ONT | moderate | 2.65% | 0.37% | 88.03% | 1.34% | 20.88% |
| MirrorRepeats | Insertion | ONT | moderate | 9.00% | 5.41% | 95.05% | 3.32% | 6.23% |
| ZDNAMotifs | Insertion | ONT | moderate | 2.36% | 63.92% | 21.22% | 51.44% | 4.13% |
| APhasedRepeats | Deletion | ONT | moderate | 8.60% | 4.45% | 46.27% | 0.15% | 68.56% |

| | | | | | | | | |
|---|---|---|---|---|---|---|---|---|
| DirectRepeats | Deletion | ONT | moderate | 10.99% | 1.12% | 57.36% | 5.07% | 53.80% |
| G4Motifs | Deletion | ONT | moderate | 23.72% | 0.67% | 22.64% | 1.91% | 12.09% |
| InvertedRepeats | Deletion | ONT | moderate | 9.16% | 0.82% | 51.86% | 12.39% | 61.75% |
| MirrorRepeats | Deletion | ONT | moderate | 12.50% | 2.25% | 75.26% | 5.63% | 34.81% |
| ZDNAMotifs | Deletion | ONT | moderate | 8.40% | 18.93% | 50.43% | 30.13% | 24.94% |
| APhasedRepeats | SNM | Illumina | stringent | 1.65% | 0.24% | 87.71% | 0.36% | 0.39% |
| DirectRepeats | SNM | Illumina | stringent | 2.48% | 9.10% | 94.94% | 0.97% | 4.64% |
| G4Motifs | SNM | Illumina | stringent | 9.56% | 2.72% | 32.66% | 3.11% | 0.24% |
| InvertedRepeats | SNM | Illumina | stringent | 1.37% | 1.96% | 85.89% | 0.33% | 1.02% |
| MirrorRepeats | SNM | Illumina | stringent | 3.03% | 2.73% | 97.33% | 3.22% | 1.11% |
| ZDNAMotifs | SNM | Illumina | stringent | 2.53% | 8.28% | 61.01% | 0.27% | 0.98% |
| APhasedRepeats | Insertion | Illumina | stringent | 5.54% | 22.76% | 39.04% | 1.38% | 0.80% |
| DirectRepeats | Insertion | Illumina | stringent | 6.82% | 82.13% | 16.52% | 8.80% | 3.37% |
| G4Motifs | Insertion | Illumina | stringent | 10.87% | 1.63% | 34.28% | 4.81% | 8.42% |
| InvertedRepeats | Insertion | Illumina | stringent | 0.76% | 20.91% | 4.31% | 11.23% | 34.39% |
| MirrorRepeats | Insertion | Illumina | stringent | 4.38% | 5.83% | 29.31% | 29.95% | 53.84% |
| ZDNAMotifs | Insertion | Illumina | stringent | 2.91% | 13.95% | 2.61% | 29.54% | 69.42% |
| APhasedRepeats | Deletion | Illumina | stringent | 0.56% | 7.17% | 31.63% | 23.29% | 27.05% |
| DirectRepeats | Deletion | Illumina | stringent | 6.14% | 87.87% | 13.57% | 17.51% | 1.42% |
| G4Motifs | Deletion | Illumina | stringent | 4.35% | 6.25% | 25.45% | 6.96% | 11.58% |
| InvertedRepeats | Deletion | Illumina | stringent | 0.45% | 31.86% | 4.99% | 5.66% | 67.91% |
| MirrorRepeats | Deletion | Illumina | stringent | 0.78% | 32.67% | 32.80% | 34.20% | 34.71% |
| ZDNAMotifs | Deletion | Illumina | stringent | 0.72% | 39.12% | 39.73% | 21.34% | 36.20% |
| APhasedRepeats | SNM | HiFi | stringent | 0.39% | 24.74% | 44.50% | 44.33% | 5.83% |
| DirectRepeats | SNM | HiFi | stringent | 0.15% | 44.92% | 24.98% | 43.35% | 1.46% |
| G4Motifs | SNM | HiFi | stringent | 0.67% | 2.26% | 32.34% | 11.40% | 4.15% |
| InvertedRepeats | SNM | HiFi | stringent | 0.09% | 5.07% | 75.74% | 19.41% | 24.65% |
| MirrorRepeats | SNM | HiFi | stringent | 0.40% | 13.49% | 28.95% | 60.26% | 14.97% |
| ZDNAMotifs | SNM | HiFi | stringent | 0.24% | 1.03% | 70.78% | 21.55% | 1.76% |
| APhasedRepeats | Insertion | HiFi | stringent | 1.73% | 10.31% | 0.24% | 0.00% | 73.24% |
| DirectRepeats | Insertion | HiFi | stringent | 1.36% | 2.67% | 5.06% | 3.48% | 98.07% |
| G4Motifs | Insertion | HiFi | stringent | 1.65% | 8.10% | 33.78% | 0.52% | 22.74% |
| InvertedRepeats | Insertion | HiFi | stringent | 1.61% | 1.74% | 0.47% | 8.35% | 90.11% |
| MirrorRepeats | Insertion | HiFi | stringent | 1.42% | 23.87% | 0.68% | 38.01% | 62.83% |
| ZDNAMotifs | Insertion | HiFi | stringent | 1.07% | 28.38% | 0.54% | 4.94% | 76.63% |
| APhasedRepeats | Deletion | HiFi | stringent | 7.31% | 1.56% | 0.41% | 0.09% | 90.15% |
| DirectRepeats | Deletion | HiFi | stringent | 9.12% | 6.24% | 0.07% | 2.45% | 93.51% |
| G4Motifs | Deletion | HiFi | stringent | 10.66% | 1.66% | 5.78% | 3.05% | 26.77% |
| InvertedRepeats | Deletion | HiFi | stringent | 12.29% | 4.85% | 0.06% | 9.28% | 90.57% |
| MirrorRepeats | Deletion | HiFi | stringent | 10.51% | 14.67% | 1.06% | 23.80% | 65.88% |
| ZDNAMotifs | Deletion | HiFi | stringent | 9.03% | 11.34% | 1.12% | 5.73% | 79.02% |
| APhasedRepeats | SNM | ONT | stringent | 1.55% | 0.23% | 71.12% | 0.14% | 0.41% |
| DirectRepeats | SNM | ONT | stringent | 1.74% | 3.16% | 91.73% | 2.73% | 0.07% |
| G4Motifs | SNM | ONT | stringent | 3.33% | 1.39% | 42.93% | 0.32% | 0.00% |
| InvertedRepeats | SNM | ONT | stringent | 1.73% | 0.48% | 85.01% | 1.76% | 2.83% |
| MirrorRepeats | SNM | ONT | stringent | 2.39% | 1.77% | 84.32% | 1.35% | 0.74% |
| ZDNAMotifs | SNM | ONT | stringent | 1.46% | 3.85% | 73.95% | 3.91% | 0.12% |
| APhasedRepeats | Insertion | ONT | stringent | 2.38% | 4.69% | 89.81% | 0.04% | 19.82% |
| DirectRepeats | Insertion | ONT | stringent | 3.21% | 0.03% | 92.67% | 0.93% | 16.29% |
| G4Motifs | Insertion | ONT | stringent | 16.98% | 1.62% | 14.51% | 0.38% | 0.05% |
| InvertedRepeats | Insertion | ONT | stringent | 1.60% | 2.91% | 88.01% | 1.15% | 24.75% |
| MirrorRepeats | Insertion | ONT | stringent | 4.31% | 8.23% | 95.56% | 6.47% | 6.00% |
| ZDNAMotifs | Insertion | ONT | stringent | 1.48% | 10.90% | 92.76% | 0.93% | 11.83% |

| | | | | | | | | |
|---|---|---|---|---|---|---|---|---|
| APhasedRepeats | Deletion | ONT | stringent | 8.57% | 3.14% | 40.26% | 0.31% | 74.95% |
| DirectRepeats | Deletion | ONT | stringent | 11.47% | 0.93% | 52.74% | 2.58% | 61.39% |
| G4Motifs | Deletion | ONT | stringent | 19.74% | 0.50% | 28.94% | 0.58% | 17.09% |
| InvertedRepeats | Deletion | ONT | stringent | 9.41% | 2.91% | 40.28% | 6.23% | 74.78% |
| MirrorRepeats | Deletion | ONT | stringent | 10.93% | 9.77% | 56.12% | 16.33% | 50.55% |
| ZDNAMotifs | Deletion | ONT | stringent | 7.40% | 0.71% | 49.01% | 4.35% | 53.33% |

**Table S5. SNM Error rates calculated from read pair overlaps.** Shown are error rates in different types of non-B motifs, calculated by detecting differences between overlapping mates in read pairs. The results for the randomized control interval

| Non-B Type | Errors | | bp of Overlap | | | | | | | |
|---|---|---|---|---|---|---|---|---|---|---|
| | Control | Motif | Control | Motif | Control | Motif | Increase | | χ² | p-value |
| APhasedRepeats | 60 | 61 | 119218 | 119424 | 0.0503 | 0.0511 | 1.02 | 1.6% | 0.007 | 0.935251 |
| DirectRepeats | 160 | 101 | 241588 | 164398 | 0.0662 | 0.0614 | 0.93 | -7.3% | 15.227 | 0.000095 |
| G4Motifs | 81 | 253 | 152530 | 132242 | 0.0531 | 0.1913 | 3.60 | 260.3% | 516.112 | < 0.00001 |
| InvertedRepeats | 791 | 676 | 1426180 | 1395058 | 0.0555 | 0.0485 | 0.87 | -12.6% | 7.764 | 0.00533 |
| MirrorRepeats | 279 | 264 | 508982 | 494961 | 0.0548 | 0.0533 | 0.97 | -2.7% | 2.376 | 0.123213 |
| ZDNAMotifs | 39 | 52 | 63404 | 44324 | 0.0615 | 0.1173 | 1.91 | 90.7% | 40.272 | < 0.00001 |

**Table S6. Sequencing success in non-B motif subregions.** In each row, aggregated (total number of errors divided by total number of nucleotides) and per-motif SNM error means are shown for each non-B motif type, the motif subregion,  error type, technology, and filtering level.

| Non-B type | Subregion | Error type | Technology | Filtering level | Overall mean | Per motif mean |
|---|---|---|---|---|---|---|
| APhasedRepeats | Repeat / stem | Deletion | HiFi | moderate | 0.0021 | 0.0019 |
| APhasedRepeats | Spacer / loop | Deletion | HiFi | moderate | 0.0007 | 0.0007 |
| APhasedRepeats | Repeat / stem | Insertion | HiFi | moderate | 0.0019 | 0.0019 |
| APhasedRepeats | Spacer / loop | Insertion | HiFi | moderate | 0.0016 | 0.0017 |
| APhasedRepeats | Repeat / stem | Single-nucleotide | HiFi | moderate | 0.0023 | 0.0013 |
| APhasedRepeats | Spacer / loop | Single-nucleotide | HiFi | moderate | 0.0033 | 0.0014 |
| DirectRepeats | Repeat / stem | Deletion | HiFi | moderate | 0.0017 | 0.0016 |
| DirectRepeats | Spacer / loop | Deletion | HiFi | moderate | 0.0015 | 0.0018 |
| DirectRepeats | Repeat / stem | Insertion | HiFi | moderate | 0.0013 | 0.0013 |
| DirectRepeats | Spacer / loop | Insertion | HiFi | moderate | 0.0015 | 0.0017 |
| DirectRepeats | Repeat / stem | Single-nucleotide | HiFi | moderate | 0.0024 | 0.0016 |
| DirectRepeats | Spacer / loop | Single-nucleotide | HiFi | moderate | 0.0029 | 0.0019 |
| G4Motifs | Repeat / stem | Deletion | HiFi | moderate | 0.0038 | 0.0036 |
| G4Motifs | Spacer / loop | Deletion | HiFi | moderate | 0.0006 | 0.0006 |
| G4Motifs | Repeat / stem | Insertion | HiFi | moderate | 0.0019 | 0.0019 |
| G4Motifs | Spacer / loop | Insertion | HiFi | moderate | 0.0022 | 0.0034 |
| G4Motifs | Repeat / stem | Single-nucleotide | HiFi | moderate | 0.0017 | 0.0017 |
| G4Motifs | Spacer / loop | Single-nucleotide | HiFi | moderate | 0.0018 | 0.0020 |
| InvertedRepeats | Repeat / stem | Deletion | HiFi | moderate | 0.0014 | 0.0014 |
| InvertedRepeats | Spacer / loop | Deletion | HiFi | moderate | 0.0012 | 0.0019 |
| InvertedRepeats | Repeat / stem | Insertion | HiFi | moderate | 0.0012 | 0.0012 |
| InvertedRepeats | Spacer / loop | Insertion | HiFi | moderate | 0.0012 | 0.0017 |
| InvertedRepeats | Repeat / stem | Single-nucleotide | HiFi | moderate | 0.0016 | 0.0014 |
| InvertedRepeats | Spacer / loop | Single-nucleotide | HiFi | moderate | 0.0016 | 0.0014 |
| MirrorRepeats | Repeat / stem | Deletion | HiFi | moderate | 0.0020 | 0.0020 |
| MirrorRepeats | Spacer / loop | Deletion | HiFi | moderate | 0.0011 | 0.0013 |
| MirrorRepeats | Repeat / stem | Insertion | HiFi | moderate | 0.0014 | 0.0014 |
| MirrorRepeats | Spacer / loop | Insertion | HiFi | moderate | 0.0011 | 0.0014 |
| MirrorRepeats | Repeat / stem | Single-nucleotide | HiFi | moderate | 0.0015 | 0.0015 |
| MirrorRepeats | Spacer / loop | Single-nucleotide | HiFi | moderate | 0.0013 | 0.0014 |
| APhasedRepeats | Repeat / stem | Deletion | Illumina | moderate | 0.0000 | 0.0000 |
| APhasedRepeats | Spacer / loop | Deletion | Illumina | moderate | 0.0000 | 0.0000 |
| APhasedRepeats | Repeat / stem | Insertion | Illumina | moderate | 0.0000 | 0.0000 |
| APhasedRepeats | Spacer / loop | Insertion | Illumina | moderate | 0.0000 | 0.0000 |
| APhasedRepeats | Repeat / stem | Single-nucleotide | Illumina | moderate | 0.0017 | 0.0017 |
| APhasedRepeats | Spacer / loop | Single-nucleotide | Illumina | moderate | 0.0024 | 0.0024 |
| DirectRepeats | Repeat / stem | Deletion | Illumina | moderate | 0.0001 | 0.0001 |
| DirectRepeats | Spacer / loop | Deletion | Illumina | moderate | 0.0001 | 0.0002 |
| DirectRepeats | Repeat / stem | Insertion | Illumina | moderate | 0.0001 | 0.0001 |
| DirectRepeats | Spacer / loop | Insertion | Illumina | moderate | 0.0001 | 0.0001 |
| DirectRepeats | Repeat / stem | Single-nucleotide | Illumina | moderate | 0.0024 | 0.0028 |
| DirectRepeats | Spacer / loop | Single-nucleotide | Illumina | moderate | 0.0028 | 0.0038 |
| G4Motifs | Repeat / stem | Deletion | Illumina | moderate | 0.0001 | 0.0001 |
| G4Motifs | Spacer / loop | Deletion | Illumina | moderate | 0.0000 | 0.0001 |
| G4Motifs | Repeat / stem | Insertion | Illumina | moderate | 0.0000 | 0.0000 |
| G4Motifs | Spacer / loop | Insertion | Illumina | moderate | 0.0000 | 0.0001 |
| G4Motifs | Repeat / stem | Single-nucleotide | Illumina | moderate | 0.0021 | 0.0021 |
| G4Motifs | Spacer / loop | Single-nucleotide | Illumina | moderate | 0.0068 | 0.0125 |

| | | | | | | |
|---|---|---|---|---|---|---|
| InvertedRepeats | Repeat / stem | Deletion | Illumina | moderate | 0.0000 | 0.0000 |
| InvertedRepeats | Spacer / loop | Deletion | Illumina | moderate | 0.0000 | 0.0000 |
| InvertedRepeats | Repeat / stem | Insertion | Illumina | moderate | 0.0000 | 0.0000 |
| InvertedRepeats | Spacer / loop | Insertion | Illumina | moderate | 0.0000 | 0.0000 |
| InvertedRepeats | Repeat / stem | Single-nucleotide | Illumina | moderate | 0.0020 | 0.0021 |
| InvertedRepeats | Spacer / loop | Single-nucleotide | Illumina | moderate | 0.0021 | 0.0022 |
| MirrorRepeats | Repeat / stem | Deletion | Illumina | moderate | 0.0001 | 0.0001 |
| MirrorRepeats | Spacer / loop | Deletion | Illumina | moderate | 0.0000 | 0.0001 |
| MirrorRepeats | Repeat / stem | Insertion | Illumina | moderate | 0.0001 | 0.0001 |
| MirrorRepeats | Spacer / loop | Insertion | Illumina | moderate | 0.0000 | 0.0001 |
| MirrorRepeats | Repeat / stem | Single-nucleotide | Illumina | moderate | 0.0021 | 0.0024 |
| MirrorRepeats | Spacer / loop | Single-nucleotide | Illumina | moderate | 0.0022 | 0.0029 |
| APhasedRepeats | Repeat / stem | Deletion | ONT | moderate | 0.0117 | 0.0111 |
| APhasedRepeats | Spacer / loop | Deletion | ONT | moderate | 0.0115 | 0.0117 |
| APhasedRepeats | Repeat / stem | Insertion | ONT | moderate | 0.0082 | 0.0084 |
| APhasedRepeats | Spacer / loop | Insertion | ONT | moderate | 0.0091 | 0.0095 |
| APhasedRepeats | Repeat / stem | Single-nucleotide | ONT | moderate | 0.0146 | 0.0124 |
| APhasedRepeats | Spacer / loop | Single-nucleotide | ONT | moderate | 0.0216 | 0.0180 |
| DirectRepeats | Repeat / stem | Deletion | ONT | moderate | 0.0105 | 0.0104 |
| DirectRepeats | Spacer / loop | Deletion | ONT | moderate | 0.0121 | 0.0132 |
| DirectRepeats | Repeat / stem | Insertion | ONT | moderate | 0.0073 | 0.0074 |
| DirectRepeats | Spacer / loop | Insertion | ONT | moderate | 0.0086 | 0.0099 |
| DirectRepeats | Repeat / stem | Single-nucleotide | ONT | moderate | 0.0171 | 0.0144 |
| DirectRepeats | Spacer / loop | Single-nucleotide | ONT | moderate | 0.0191 | 0.0169 |
| G4Motifs | Repeat / stem | Deletion | ONT | moderate | 0.0241 | 0.0229 |
| G4Motifs | Spacer / loop | Deletion | ONT | moderate | 0.0132 | 0.0137 |
| G4Motifs | Repeat / stem | Insertion | ONT | moderate | 0.0155 | 0.0160 |
| G4Motifs | Spacer / loop | Insertion | ONT | moderate | 0.0129 | 0.0179 |
| G4Motifs | Repeat / stem | Single-nucleotide | ONT | moderate | 0.0231 | 0.0227 |
| G4Motifs | Spacer / loop | Single-nucleotide | ONT | moderate | 0.0192 | 0.0189 |
| InvertedRepeats | Repeat / stem | Deletion | ONT | moderate | 0.0109 | 0.0109 |
| InvertedRepeats | Spacer / loop | Deletion | ONT | moderate | 0.0114 | 0.0142 |
| InvertedRepeats | Repeat / stem | Insertion | ONT | moderate | 0.0076 | 0.0076 |
| InvertedRepeats | Spacer / loop | Insertion | ONT | moderate | 0.0077 | 0.0107 |
| InvertedRepeats | Repeat / stem | Single-nucleotide | ONT | moderate | 0.0193 | 0.0154 |
| InvertedRepeats | Spacer / loop | Single-nucleotide | ONT | moderate | 0.0176 | 0.0169 |
| MirrorRepeats | Repeat / stem | Deletion | ONT | moderate | 0.0117 | 0.0117 |
| MirrorRepeats | Spacer / loop | Deletion | ONT | moderate | 0.0102 | 0.0107 |
| MirrorRepeats | Repeat / stem | Insertion | ONT | moderate | 0.0077 | 0.0076 |
| MirrorRepeats | Spacer / loop | Insertion | ONT | moderate | 0.0068 | 0.0078 |
| MirrorRepeats | Repeat / stem | Single-nucleotide | ONT | moderate | 0.0135 | 0.0132 |
| MirrorRepeats | Spacer / loop | Single-nucleotide | ONT | moderate | 0.0159 | 0.0154 |
| APhasedRepeats | Repeat / stem | Deletion | HiFi | stringent | 0.0020 | 0.0018 |
| APhasedRepeats | Spacer / loop | Deletion | HiFi | stringent | 0.0007 | 0.0007 |
| APhasedRepeats | Repeat / stem | Insertion | HiFi | stringent | 0.0019 | 0.0019 |
| APhasedRepeats | Spacer / loop | Insertion | HiFi | stringent | 0.0015 | 0.0016 |
| APhasedRepeats | Repeat / stem | Single-nucleotide | HiFi | stringent | 0.0012 | 0.0012 |
| APhasedRepeats | Spacer / loop | Single-nucleotide | HiFi | stringent | 0.0013 | 0.0013 |
| DirectRepeats | Repeat / stem | Deletion | HiFi | stringent | 0.0013 | 0.0013 |
| DirectRepeats | Spacer / loop | Deletion | HiFi | stringent | 0.0014 | 0.0014 |
| DirectRepeats | Repeat / stem | Insertion | HiFi | stringent | 0.0011 | 0.0011 |

| | | | | | | |
|---|---|---|---|---|---|---|
| DirectRepeats | Spacer / loop | Insertion | HiFi | stringent | 0.0013 | 0.0015 |
| DirectRepeats | Repeat / stem | Single-nucleotide | HiFi | stringent | 0.0014 | 0.0014 |
| DirectRepeats | Spacer / loop | Single-nucleotide | HiFi | stringent | 0.0015 | 0.0015 |
| G4Motifs | Repeat / stem | Deletion | HiFi | stringent | 0.0042 | 0.0040 |
| G4Motifs | Spacer / loop | Deletion | HiFi | stringent | 0.0007 | 0.0007 |
| G4Motifs | Repeat / stem | Insertion | HiFi | stringent | 0.0020 | 0.0019 |
| G4Motifs | Spacer / loop | Insertion | HiFi | stringent | 0.0021 | 0.0032 |
| G4Motifs | Repeat / stem | Single-nucleotide | HiFi | stringent | 0.0014 | 0.0015 |
| G4Motifs | Spacer / loop | Single-nucleotide | HiFi | stringent | 0.0015 | 0.0018 |
| InvertedRepeats | Repeat / stem | Deletion | HiFi | stringent | 0.0015 | 0.0015 |
| InvertedRepeats | Spacer / loop | Deletion | HiFi | stringent | 0.0015 | 0.0020 |
| InvertedRepeats | Repeat / stem | Insertion | HiFi | stringent | 0.0013 | 0.0013 |
| InvertedRepeats | Spacer / loop | Insertion | HiFi | stringent | 0.0014 | 0.0019 |
| InvertedRepeats | Repeat / stem | Single-nucleotide | HiFi | stringent | 0.0014 | 0.0013 |
| InvertedRepeats | Spacer / loop | Single-nucleotide | HiFi | stringent | 0.0014 | 0.0014 |
| MirrorRepeats | Repeat / stem | Deletion | HiFi | stringent | 0.0020 | 0.0021 |
| MirrorRepeats | Spacer / loop | Deletion | HiFi | stringent | 0.0012 | 0.0013 |
| MirrorRepeats | Repeat / stem | Insertion | HiFi | stringent | 0.0014 | 0.0014 |
| MirrorRepeats | Spacer / loop | Insertion | HiFi | stringent | 0.0011 | 0.0013 |
| MirrorRepeats | Repeat / stem | Single-nucleotide | HiFi | stringent | 0.0015 | 0.0015 |
| MirrorRepeats | Spacer / loop | Single-nucleotide | HiFi | stringent | 0.0013 | 0.0015 |
| APhasedRepeats | Repeat / stem | Deletion | Illumina | stringent | 0.0000 | 0.0000 |
| APhasedRepeats | Spacer / loop | Deletion | Illumina | stringent | 0.0000 | 0.0000 |
| APhasedRepeats | Repeat / stem | Insertion | Illumina | stringent | 0.0000 | 0.0000 |
| APhasedRepeats | Spacer / loop | Insertion | Illumina | stringent | 0.0000 | 0.0000 |
| APhasedRepeats | Repeat / stem | Single-nucleotide | Illumina | stringent | 0.0017 | 0.0018 |
| APhasedRepeats | Spacer / loop | Single-nucleotide | Illumina | stringent | 0.0023 | 0.0024 |
| DirectRepeats | Repeat / stem | Deletion | Illumina | stringent | 0.0001 | 0.0001 |
| DirectRepeats | Spacer / loop | Deletion | Illumina | stringent | 0.0001 | 0.0001 |
| DirectRepeats | Repeat / stem | Insertion | Illumina | stringent | 0.0000 | 0.0000 |
| DirectRepeats | Spacer / loop | Insertion | Illumina | stringent | 0.0000 | 0.0000 |
| DirectRepeats | Repeat / stem | Single-nucleotide | Illumina | stringent | 0.0023 | 0.0025 |
| DirectRepeats | Spacer / loop | Single-nucleotide | Illumina | stringent | 0.0025 | 0.0030 |
| G4Motifs | Repeat / stem | Deletion | Illumina | stringent | 0.0001 | 0.0001 |
| G4Motifs | Spacer / loop | Deletion | Illumina | stringent | 0.0000 | 0.0001 |
| G4Motifs | Repeat / stem | Insertion | Illumina | stringent | 0.0000 | 0.0000 |
| G4Motifs | Spacer / loop | Insertion | Illumina | stringent | 0.0000 | 0.0001 |
| G4Motifs | Repeat / stem | Single-nucleotide | Illumina | stringent | 0.0020 | 0.0020 |
| G4Motifs | Spacer / loop | Single-nucleotide | Illumina | stringent | 0.0060 | 0.0107 |
| InvertedRepeats | Repeat / stem | Deletion | Illumina | stringent | 0.0000 | 0.0000 |
| InvertedRepeats | Spacer / loop | Deletion | Illumina | stringent | 0.0000 | 0.0000 |
| InvertedRepeats | Repeat / stem | Insertion | Illumina | stringent | 0.0000 | 0.0000 |
| InvertedRepeats | Spacer / loop | Insertion | Illumina | stringent | 0.0000 | 0.0000 |
| InvertedRepeats | Repeat / stem | Single-nucleotide | Illumina | stringent | 0.0020 | 0.0021 |
| InvertedRepeats | Spacer / loop | Single-nucleotide | Illumina | stringent | 0.0022 | 0.0023 |
| MirrorRepeats | Repeat / stem | Deletion | Illumina | stringent | 0.0000 | 0.0000 |
| MirrorRepeats | Spacer / loop | Deletion | Illumina | stringent | 0.0000 | 0.0000 |
| MirrorRepeats | Repeat / stem | Insertion | Illumina | stringent | 0.0000 | 0.0000 |
| MirrorRepeats | Spacer / loop | Insertion | Illumina | stringent | 0.0000 | 0.0000 |
| MirrorRepeats | Repeat / stem | Single-nucleotide | Illumina | stringent | 0.0020 | 0.0022 |
| MirrorRepeats | Spacer / loop | Single-nucleotide | Illumina | stringent | 0.0022 | 0.0027 |

| | | | | | | |
|---|---|---|---|---|---|---|
| APhasedRepeats | Repeat / stem | Deletion | ONT | stringent | 0.0118 | 0.0111 |
| APhasedRepeats | Spacer / loop | Deletion | ONT | stringent | 0.0115 | 0.0117 |
| APhasedRepeats | Repeat / stem | Insertion | ONT | stringent | 0.0082 | 0.0083 |
| APhasedRepeats | Spacer / loop | Insertion | ONT | stringent | 0.0091 | 0.0094 |
| APhasedRepeats | Repeat / stem | Single-nucleotide | ONT | stringent | 0.0126 | 0.0124 |
| APhasedRepeats | Spacer / loop | Single-nucleotide | ONT | stringent | 0.0180 | 0.0180 |
| DirectRepeats | Repeat / stem | Deletion | ONT | stringent | 0.0109 | 0.0109 |
| DirectRepeats | Spacer / loop | Deletion | ONT | stringent | 0.0119 | 0.0130 |
| DirectRepeats | Repeat / stem | Insertion | ONT | stringent | 0.0073 | 0.0073 |
| DirectRepeats | Spacer / loop | Insertion | ONT | stringent | 0.0083 | 0.0093 |
| DirectRepeats | Repeat / stem | Single-nucleotide | ONT | stringent | 0.0164 | 0.0160 |
| DirectRepeats | Spacer / loop | Single-nucleotide | ONT | stringent | 0.0179 | 0.0176 |
| G4Motifs | Repeat / stem | Deletion | ONT | stringent | 0.0224 | 0.0209 |
| G4Motifs | Spacer / loop | Deletion | ONT | stringent | 0.0125 | 0.0128 |
| G4Motifs | Repeat / stem | Insertion | ONT | stringent | 0.0151 | 0.0156 |
| G4Motifs | Spacer / loop | Insertion | ONT | stringent | 0.0122 | 0.0166 |
| G4Motifs | Repeat / stem | Single-nucleotide | ONT | stringent | 0.0238 | 0.0234 |
| G4Motifs | Spacer / loop | Single-nucleotide | ONT | stringent | 0.0185 | 0.0183 |
| InvertedRepeats | Repeat / stem | Deletion | ONT | stringent | 0.0117 | 0.0117 |
| InvertedRepeats | Spacer / loop | Deletion | ONT | stringent | 0.0132 | 0.0155 |
| InvertedRepeats | Repeat / stem | Insertion | ONT | stringent | 0.0080 | 0.0080 |
| InvertedRepeats | Spacer / loop | Insertion | ONT | stringent | 0.0092 | 0.0118 |
| InvertedRepeats | Repeat / stem | Single-nucleotide | ONT | stringent | 0.0163 | 0.0160 |
| InvertedRepeats | Spacer / loop | Single-nucleotide | ONT | stringent | 0.0176 | 0.0175 |
| MirrorRepeats | Repeat / stem | Deletion | ONT | stringent | 0.0127 | 0.0127 |
| MirrorRepeats | Spacer / loop | Deletion | ONT | stringent | 0.0106 | 0.0112 |
| MirrorRepeats | Repeat / stem | Insertion | ONT | stringent | 0.0078 | 0.0078 |
| MirrorRepeats | Spacer / loop | Insertion | ONT | stringent | 0.0070 | 0.0082 |
| MirrorRepeats | Repeat / stem | Single-nucleotide | ONT | stringent | 0.0147 | 0.0146 |
| MirrorRepeats | Spacer / loop | Single-nucleotide | ONT | stringent | 0.0166 | 0.0163 |

**Table S7. Adjusted p-values for comparing SNM error rates between subregions of non-B motifs.** Shown are Benjamini-Hochberg corrected p-values calculated for t-test comparing SNM error rates between different subregions (repeat arm/ stem vs. spacer / loop) of non-B motifs.

| Type | Technology | Filtering level | adjusted P-value |
|---|---|---|---|
| APhasedRepeats | Illumina | moderate | **0** |
| DirectRepeats | Illumina | moderate | **1.12E-105** |
| G4Motifs | Illumina | moderate | **0** |
| InvertedRepeats | Illumina | moderate | **1.78E-73** |
| MirrorRepeats | Illumina | moderate | **2.65E-111** |
| APhasedRepeats | HiFi | moderate | **0.020** |
| DirectRepeats | HiFi | moderate | **2.22E-09** |
| G4Motifs | HiFi | moderate | **3.04E-11** |
| InvertedRepeats | HiFi | moderate | **1.55E-11** |
| MirrorRepeats | HiFi | moderate | 0.996 |
| APhasedRepeats | ONT | moderate | **0** |
| DirectRepeats | ONT | moderate | **9.99E-254** |
| G4Motifs | ONT | moderate | **0** |
| InvertedRepeats | ONT | moderate | **0** |
| MirrorRepeats | ONT | moderate | **0** |
| APhasedRepeats | Illumina | stringent | **5.04E-77** |
| DirectRepeats | Illumina | stringent | **0.000** |
| G4Motifs | Illumina | stringent | **0** |
| InvertedRepeats | Illumina | stringent | **1.47E-16** |
| MirrorRepeats | Illumina | stringent | **6.74E-05** |
| APhasedRepeats | HiFi | stringent | **0.995** |
| DirectRepeats | HiFi | stringent | 0.996 |
| G4Motifs | HiFi | stringent | **0.005** |
| InvertedRepeats | HiFi | stringent | **0.001** |
| MirrorRepeats | HiFi | stringent | 0.996 |
| APhasedRepeats | ONT | stringent | **0** |
| DirectRepeats | ONT | stringent | **7.58E-11** |
| G4Motifs | ONT | stringent | **4.86E-159** |
| InvertedRepeats | ONT | stringent | **1.19E-201** |
| MirrorRepeats | ONT | stringent | **1.63E-16** |

**Table S8. Expected number of false-positive SNVs in three existing datasets.**

| | | | | Expected number of FP SNVs | |
|---|---|---|---|---|---|
| **Dataset** | **Number of haploid genomes** | **MAF** | **Read depth (per haploid genome)** | **NonB** | **BDNA** |
| 1000Genomes | 5,008 | Singletons | 1 | 109,615,879 | 109,617,975 |
| SGDP | 600 | Singletons | 21 | 0 | 0 |
| gnomAD | 152,312 | Singletons | 15 | 11,044 | 2,480 |
| 1000Genomes | 5,008 | Tripletons | 1 | 109,372,266 | 109,484,576 |
| SGDP | 600 | Tripletons | 21 | 0 | 0 |
| gnomAD | 152312 | Tripletons | 15 | 0 | 0 |
| 1000Genomes | 5008 | 0.01 | 1 | 19 | 0 |
| SGDP | 600 | 0.01 | 21 | 0 | 0 |
| gnomAD | 152312 | 0.01 | 15 | 0 | 0 |
